# Sex-specific microRNA regulators of Parkinson’s disease: insights from cohort-stratified simulations of compensatory pathway dynamics

**DOI:** 10.1101/2025.09.29.679145

**Authors:** Ahmed Abdelmonem Hemedan, Marek Ostaszewski, Armin Rauschenberger, Lukas Pavelka, Pierre Klemmer, Enrico Glaab, Reinhard Schneider, Rejko Krüger, Venkata P. Satagopam, the NCER-PD consortium

**Author notes:** Ahmed Abdelmonem Hemedan and Marek Ostaszewski share the first authorship.

## Abstract

Parkinson’s disease (PD) exhibits sex differences in prevalence, symptom severity, and progression, suggesting distinct underlying molecular mechanisms. However, the pathophysiological mechanisms remain largely obscure, particularly in the context of the post-transcriptional regulations, where microRNAs (miRNAs) suppress the expression of multiple genes. Bulk transcriptomic data often blur these effects, especially when regulatory patterns vary by sex or cell type. miRNAs have emerged as key regulators of PD-related processes, but their complexity demands computational methods that enable capturing the functional impact at the pathway level.

In this study, we investigated how sex may affect miRNA-regulated PD pathways using Boolean modeling across two large PD cohorts, the Parkinson’s Progression Markers Initiative (PPMI) and the Luxembourg Parkinson’s Study (LuxPark). First, differential expression analysis identified significant variations in miRNA expression between the sexes across both cohorts. These miRNAs were analysed to identify molecular pathways that are over-represented among the predicted targets of the dysregulated microRNAs (i.e. enrichment analysis). The enriched pathways were used to build Boolean models to simulate the effects of sex-specific miRNA dysregulation. These simulations showed consistent male-specific impairment in mitochondrial biogenesis, and respiratory chain activity. Mitophagy and oxidative stress response pathways were also disrupted, alongside dysregulation of autophagy-related protein-folding mechanisms.

Our findings suggest that sex-specific miRNA dysregulation contributes to differences in molecular patterns in PD by influencing compensatory related pathways and responses. These results highlight the need for sex-stratified approaches in modeling, translational research, and precision medicine strategies for disease-modifying treatments.

## 1. Introduction

Parkinson’s Disease (PD) is a progressive neurodegenerative disorder, which exhibits notable sex-based differences in prevalence and progression. Epidemiological studies reveal a higher incidence in males, with a male-to-female ratio of approximately 1.5-2:1, suggesting that the biological sex influences susceptibility to the disease^1^. Emerging research highlights sex-specific microRNAs (miRNAs), small non-coding RNAs regulating gene expression, as key modulators of pathways implicated in PD^2^. These findings underscore the potential of miRNAs as contributors to sex-linked variability in PD onset, symptom severity, and progression.

The pathophysiology of PD involves different pathways, e.g. mitochondrial impairment, oxidative stress, synaptic dysfunction, and neuroinflammation, which collectively contribute to neurodegeneration^1^. However, clinical progression varies significantly between individuals, with sex-specific physiology likely contributing to this heterogeneity^3,4^. For instance, estrogen in females directly exhibits neuroprotective effects by maintaining mitochondrial resilience and mitigating oxidative stress, whereas testosterone in males directly influences dopamine receptor signaling, by upregulating dopamine transporter and D₂ receptor transcripts and reducing D₃ receptor expression in the nigrostriatal pathway^5,6^. Such hormonal disparities thus contribute to distinct molecular patterns in males and females.

Additionally, estrogen and testosterone indirectly shape molecular responses through modulation of miRNA expression, potentially increasing neuroprotective mechanisms in females and exacerbating inflammatory responses in males^7,8,9^. While miRNAs are critical post-transcriptional regulators of PD-related pathways, their sex-specific role remains largely unexplored^10,11,12^. Elucidating these mechanisms could explain why males experience higher PD risk and potentially faster progression, whereas females often present with delayed onset and distinct clinical features^7,13^.

To investigate the effect of sex-specific miRNAs, we first analyzed data from two well-characterized PD cohorts (LuxPark and PPMI) to identify sex-specific differentially expressed miRNAs. These miRNAs were analysed to identify PD-related pathways that are over-represented among the predicted targets of the dysregulated microRNAs using PD Map^14^. This process is referred to as pathway enrichment.

PD map encapsulates extensive knowledge about PD-related mechanisms, offering a crucial tool for visualizing and understanding molecular interactions implicated in the disease^14^. The PD map represents the largest repository of PD pathways available to date, including detailed knowledge about PD-related mechanisms. It serves as an essential tool for visualizing molecular interactions implicated in the disease. The pathway diagrams in the PD map are static but can be modeled and simulated to understand the dynamics of the represented mechanisms.

Based on these pathways, we constructed a computational model to simulate the effects of sex-specific miRNA dysregulation, applying the logical modeling approach called Probabilistic Boolean Networks (PBNs)^11,15^. Simulating the model and studying its dynamics allowed us to quantify the impact of miRNA dysregulation on key pathways (e.g., dopamine metabolism, mitochondrial function) across the LuxPark and PPMI cohorts^11,12^.

This study aims to investigate how sex-specific miRNAs regulate key molecular pathways in PD by combining differential expression analysis from two large cohorts (LuxPark and PPMI) with pathway enrichment and Boolean modeling (Figure 1). Our aim is to identify reproducible regulatory patterns that explain sex differences in disease mechanisms and progression. Our results reveal distinct sex-specific miRNA regulatory patterns influencing molecular pathway activity in PD. Males exhibited early engagement of compensatory mechanisms, whereas females displayed a more delayed response. These findings suggest that miRNA-mediated regulation contributes to sex-specific molecular adaptations in PD.

**Figure 1.**
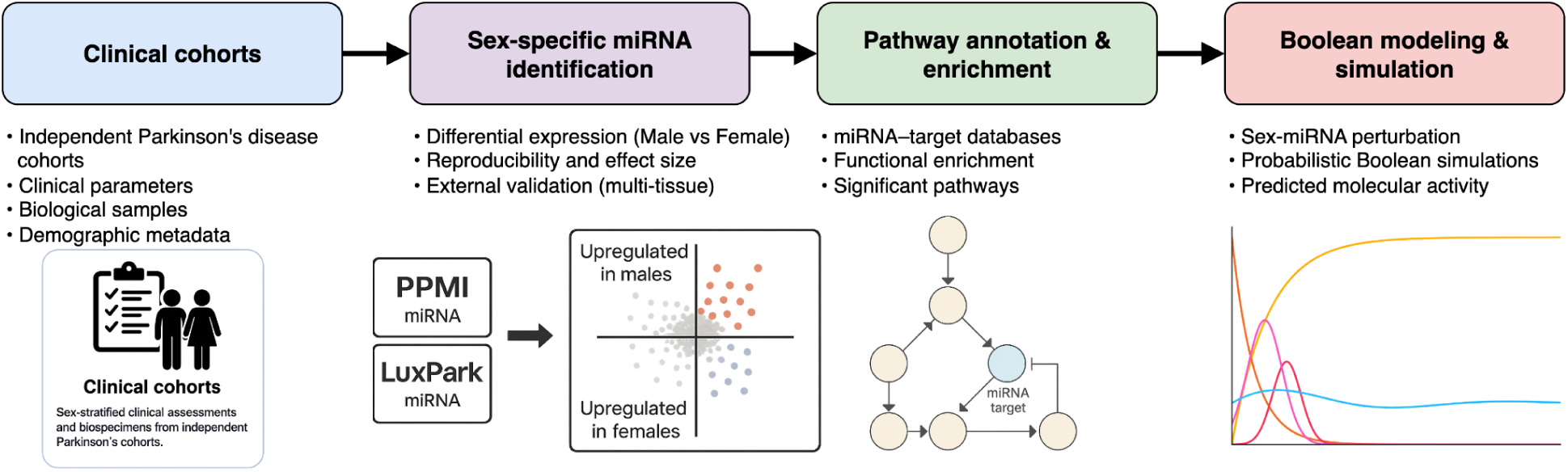
Schematic workflow and output from cohort data to simulated molecular activity. PD cohort data (PPMI, LuxPark) were used to identify sex-specific miRNAs, followed by pathway enrichment and Boolean modeling. Simulations predicted molecular activity patterns under sex-specific regulatory conditions.

The following sections detail (1) cohort characteristics and differential miRNA expression analysis, (2) Boolean modeling process and simulation results, and (3) a discussion on the results in the context of sex-dependent molecular mechanisms in PD highlights their relevance to disease progression and regulatory dynamics.

## 2. Results

### 2.1 Sex-dysregulated miRNAs in PD: pathway associations and disease-specific stratification

We first identified miRNAs with sex-specific expression in PD patients by performing differential expression analyses between males and females within LuxPark and PPMI cohorts (Figure 2). After correcting for multiple testing, 44 miRNAs in LuxPark (32 downregulated, 12 upregulated in males) and 67 in PPMI (41 downregulated, 26 upregulated in males) exhibited statistically significant sex-associated expression differences. Disease duration was not included as a covariate in the differential expression models, as it is intrinsically confounded with cohort design: PPMI participants were newly diagnosed and medication-naïve, while LuxPark patients had longer disease duration and ongoing treatment. To control for this, all analyses were performed separately within each cohort. This stratified approach preserved cohort-specific structure while allowing us to compare consistency of sex-associated miRNA signals across distinct clinical contexts.

**Figure 2:**
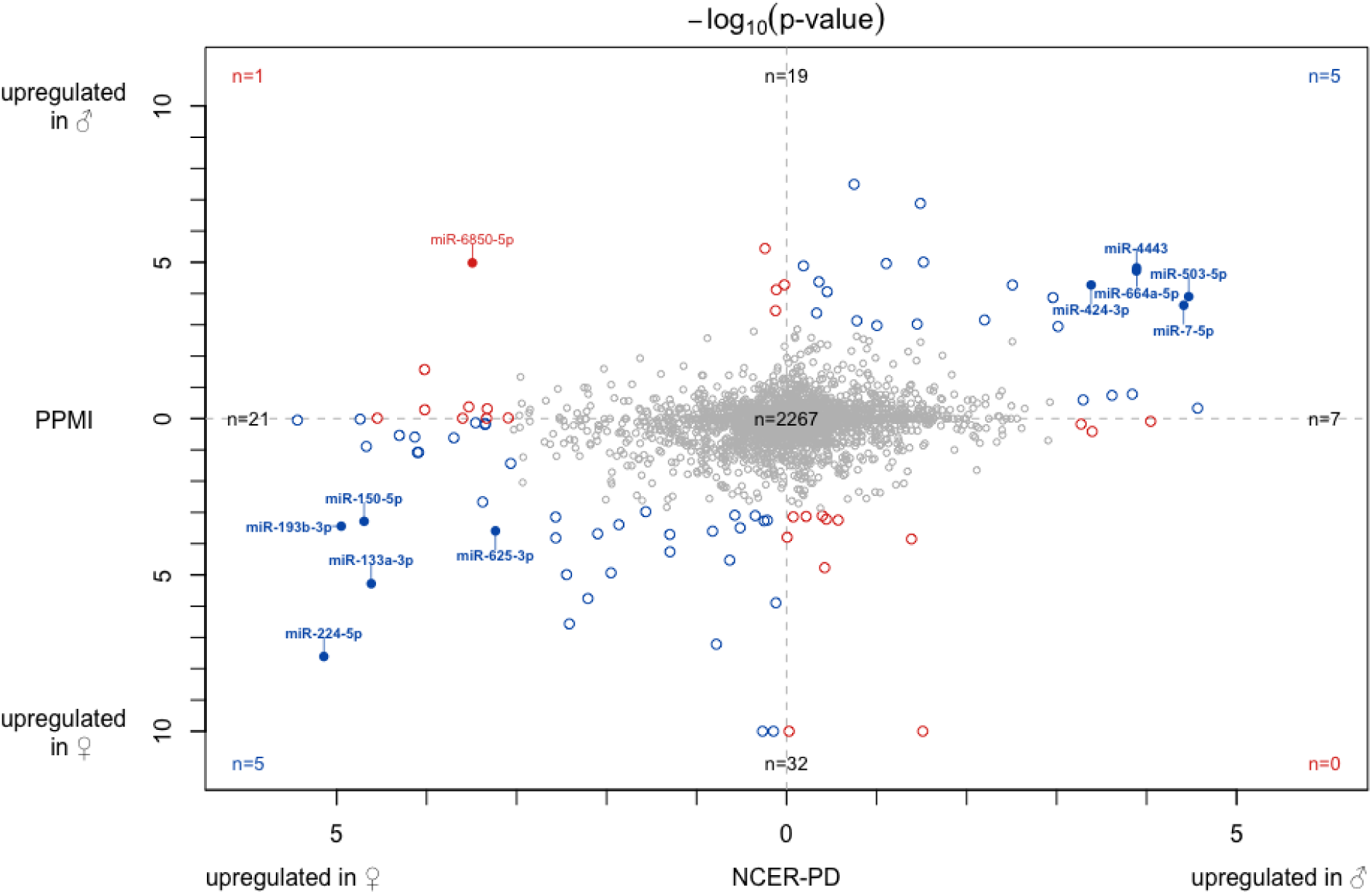
Sex-specific differential expression of miRNAs within PD patients from the LuxPark and PPMI cohorts. Each point represents a miRNA shared across both datasets (n = 2,357), plotted by its −log₁₀(p-value) from sex-specific comparisons: LuxPark on the x-axis and PPMI on the y-axis. Sex differences were assessed within each cohort (PD males vs PD females). The direction of differential expression is inferred from the fold change: right/up = higher in PD males; left/down = higher in PD females. Filled circles indicate miRNAs significant in both cohorts (FDR < 0.05). Color reflects directionality: blue for consistent sex bias (e.g., up in males in both cohorts), red for opposite direction. The numbers at each corner indicate counts of miRNAs in each quadrant. Extreme p-values (−log₁₀ > 10) were clipped for clarity.

To assess the disease-context specificity of these patterns, we compared miRNA sex effects in PD to those in healthy controls (HC). To refine the PD-specific signal, we compared sex differences in miRNA expression between PD and HC groups. For each miRNA, we classified the effect direction as positive (up in males), negative (up in females), or nonsignificant. We then excluded any miRNAs that showed a significant sex difference in HC alone, or were significantly sex-biased in both PD and HC with the same direction. This yielded a refined set of 26 HC-filtered miRNAs in LuxPark and 56 in PPMI, suggesting that many of the sex-associated miRNA changes in PD are amplified or uniquely present in the disease state.

A complete summary of differential expression results, including sex effects in PD, sex effects in HC, and sex-by-disease interaction terms, is provided in Table S10. This table also indicates which miRNAs passed disease-specific filters and were used in subsequent modeling.

Among the 2,357 miRNAs shared across both datasets, 10 were significantly differentially expressed between males and females in both LuxPark and PPMI (FDR < 0.05), with consistent direction of effect.

In Figure 2, the overlaid 3 x 3 contingency table shows the counts of significantly upregulated miRNAs in women (left or bottom) and men (right or top) for the two cohorts. The two-sided Fisher’s exact test for count data yields a p-value of 4.3 × 10^-11, rejecting the null hypothesis of independence. It shows that the direction of sex-specific miRNA expression is not random across cohorts. miRNAs upregulated in males or females in one cohort tend to show the same pattern in the other. This points to a consistent sex effect across both datasets.

Table 1 lists the 10 miRNAs that were significantly differentially expressed between males and females in both LuxPark and PPMI, with consistent direction of effect.

**Table 1:**
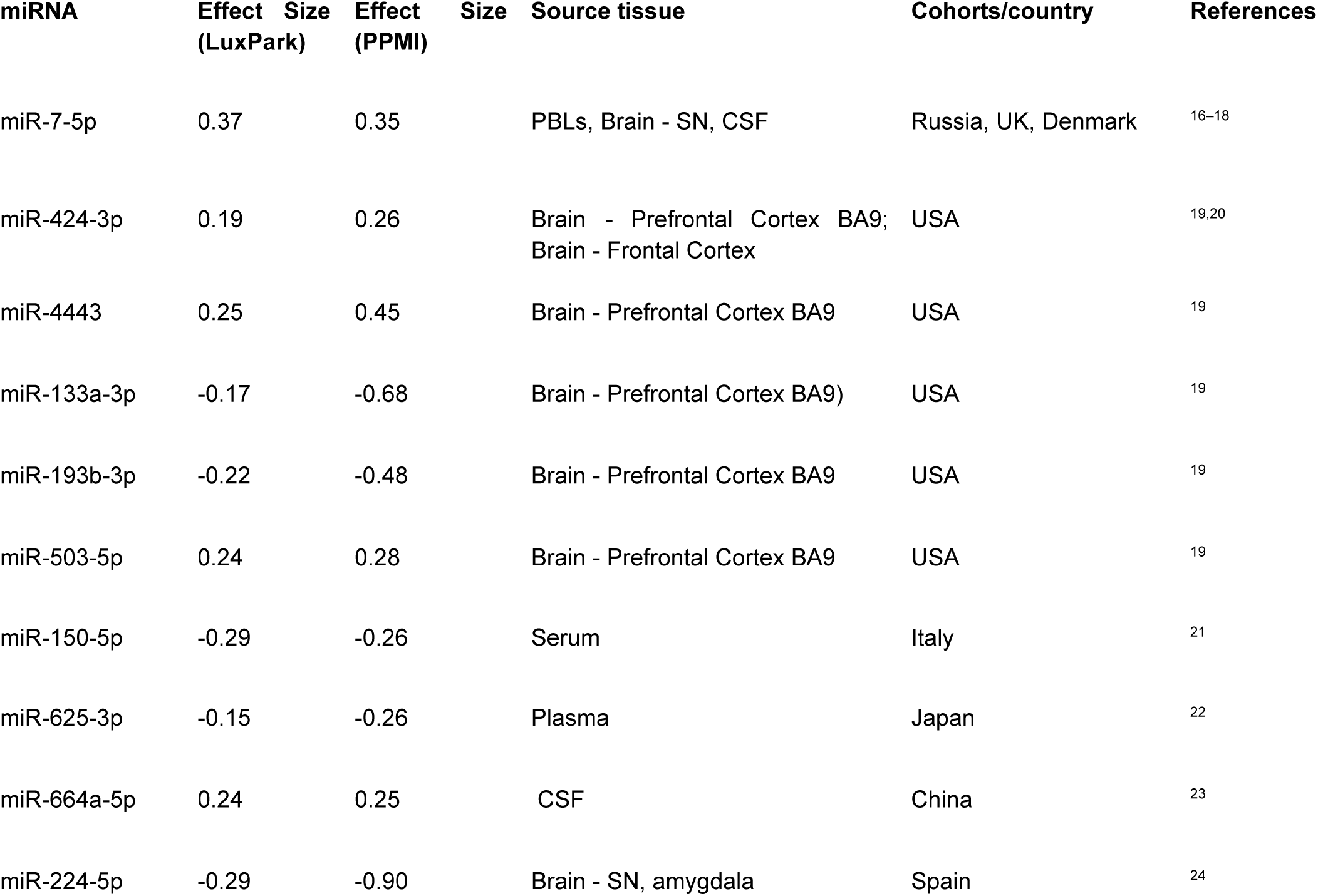
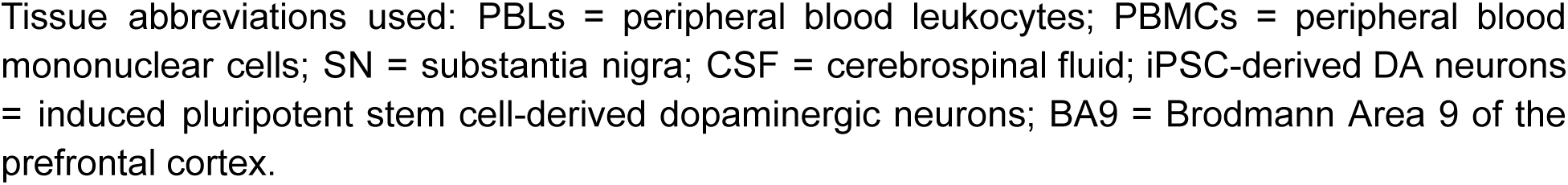
Overlapping significant sex-specific miRNAs between LuxPark and PPMI, used for enrichment and Boolean modeling. All miRNAs listed were consistently differentially expressed between males and females in both cohorts.

Tissue abbreviations used: PBLs = peripheral blood leukocytes; PBMCs = peripheral blood mononuclear cells; SN = substantia nigra; CSF = cerebrospinal fluid; iPSC-derived DA neurons = induced pluripotent stem cell-derived dopaminergic neurons; BA9 = Brodmann Area 9 of the prefrontal cortex.

Following the identification of sex-specific miRNAs within each cohort, we assessed their potential impact on disease-relevant pathways. Predicted miRNA targets were compiled using consensus across multiple reference databases (see Methods), then filtered for Parkinson’s disease–associated genes (Supplementary Table S11). These targets were mapped onto the Parkinson’s Disease Map (PD Map), a curated representation of PD-related molecular mechanisms.

This PD-specific enrichment step (Supplementary Table S12) defined the pathways used for Boolean model construction. For both LuxPark and PPMI cohorts, we built and simulated models parameterized with cohort-specific miRNA perturbations. This allowed us to quantify predicted differences in pathway activity under sex-stratified conditions and formed the central analytical framework of the study.

To preserve disease relevance, only PD map–derived pathways were used in the modeling. In parallel, broader enrichment analyses were conducted using (Kyoto Encyclopedia of Genes and Genomes) KEGG, Reactome, Gene Ontology (GO), and (Molecular Signatures Database)MSigDB. These results were not used for model construction but provided additional functional context, including recurrent enrichment of hormone-related and neuroendocrine signaling pathways (Supplementary Table S13).

To examine potential upstream regulators of the sex-specific miRNAs, we identified validated transcription factors (TFs) using the TransmiR database. Although not included in the simulation pipeline, these TFs were enriched for pathways related to dopaminergic signaling, hormonal regulation, and sex-specific transcriptional programs (Supplementary Table S14). Literature evidence confirmed that most of these TFs are modulated by sex hormones, suggesting a hormonal contribution to the observed patterns of miRNA dysregulation [miRNA-TF mapping and enrichment](https://docs.google.com/document/u/0/d/1MSy4j2rDHkDwa49R7cRpnga5ZI2CAxaTvHB0xfyIxmg/edit)

### 2.2 Cohort-specific differences in pathway activity patterns from Boolean simulations

Boolean models were constructed from the pathways previously identified through PD map enrichment (Supplementary Table S12). To assess molecular dysregulation linked to sex-specific miRNA profiles, we performed Boolean simulations under three conditions: using the full set of filtered miRNAs from each cohort independently (LuxPark, PPMI), as well as their shared overlap (Table 1). This allowed us to explore both distinct and convergent patterns of molecular activity across cohorts. Notably, even when shared miRNAs were used as inputs, the resulting simulations sometimes diverged across cohorts. This reflects differences in the magnitude of miRNA expression changes and target probabilities between LuxPark and PPMI, which shape pathway behavior even under partially overlapping regulatory inputs. The resulting activity probabilities revealed sex- and cohort-specific differences across multiple molecular processes (Figure 3), including mitochondrial function, oxidative stress, protein handling, and synaptic regulation. Extended results and additional simulation outputs are provided in the supplementary files. [NCER-PPMI Supplementary](https://docs.google.com/document/d/1mrjUDz5ySpiBKIJx_vlyYWQG7cVpq_HiAg0v0XMW6Ks/edit?tab=t.0) [Results- graphs](https://docs.google.com/document/d/14_8_a7J0FZ9aMOCEqHwoJA6voAB4Sx7GbE-wLgEPrmQ/edit?tab=t.0)

**Figure 3.**
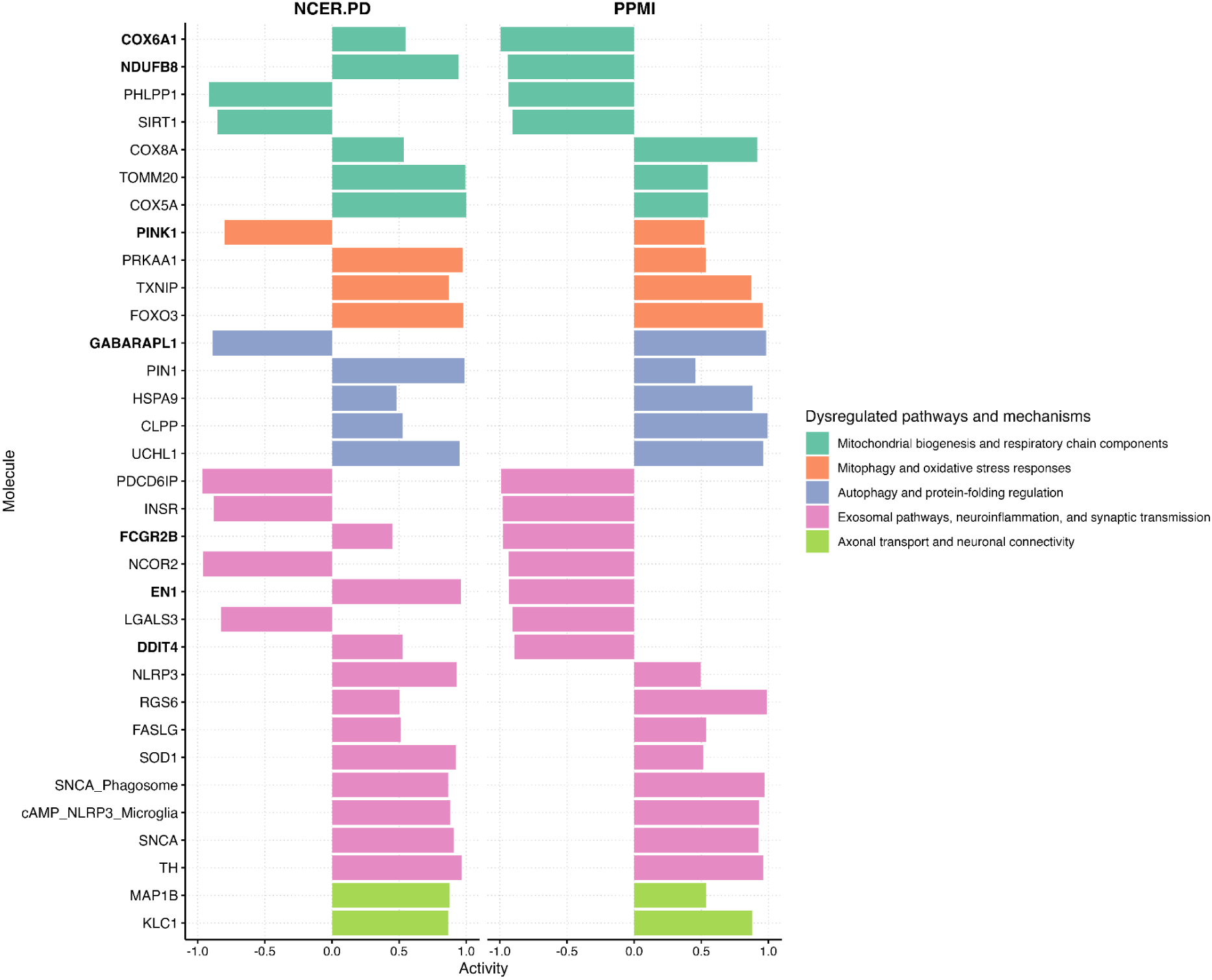
Boolean-simulated activity probabilities for key molecules in LuxPark and PPMI cohorts. Bar direction indicates activation or inhibition; colors reflect functional categories. Bold labels mark molecules with opposing activity between cohorts.

#### 2.2.1 Cohort and sex-specific molecular dysregulation

Simulations across both cohorts revealed Cohort- and sex-specific dysregulation within key biological domains relevant to PD. These included mitochondrial homeostasis, oxidative stress responses, protein quality control, exosomal signaling, and neuronal connectivity. While the same pathways were implicated in both cohorts, the direction and extent of activity changes varied, reflecting differences in sex-specific miRNA profiles and expression magnitudes. The following sections detail these pathway-level differences, organized by functional domain, to highlight shared patterns and divergence.

##### Mitochondrial biogenesis and respiratory chain components

Mitochondrial complex subunits (COX5A, COX8A, COX6A1, NDUFB8) and PD-associated regulators (PINK1, PRKN) showed decreased activity in both cohorts (Table S2). In simulations, males showed higher mitochondrial impairment compared to females. The miRNA profile of the PPMI cohort exhibited more downregulation of mitochondrial biogenesis factors (NFE2L2, SIRT1, PHLPP1) compared to LuxPark (Table S2). Activity of COX8A was increased in PPMI, whereas COX6A1 and NDUFB8 were decreased. Simulations indicate increased activity of PGC-1α and COX5A across both cohorts (Table S1).

##### Mitophagy and oxidative stress responses

The regulation of mitophagy showed cohort-level differences: activity of its regulators PRKN and GABARAPL1 increased in LuxPark, while in PPMI GABARAPL1 activity decreased and PRKN remained unchanged (Table S1). Mitophagy-related activity (PINK1, PRKN) appeared weaker in males compared to females across both cohorts. FOXO3 and MAPK9, regulators of oxidative stress response, were upregulated in both cohorts. The predicted TXNIP and SIRT2 activity is increased across both cohorts (Table S2).

##### Autophagy and protein-folding regulation

Regulation of autophagy differed across cohorts. In LuxPark, predicted activity of HSPA8 and HSPB1 was decreased, whereas activity of MAP1LC3B was increased (Table S3, Table S4). In contrast, in PPMI, HSPA8 activity was predicted to be increased in male patients. Specifically, the model revealed increased activation of HSPA8 and decreased activation of HSPB1. Simulations also predicted higher α-synuclein accumulation in males in both cohorts.

##### Exosomal pathways, neuroinflammation, and synaptic transmission

Increased misfolded protein load in simulations was accompanied by activation of exosomal pathways, including exosome-associated proteins LGALS3 and PDCD6IP, critical in synaptic function and protein clearance mechanisms. Both cohorts showed dysregulated activity patterns (Table S5). Simulation outcomes indicated higher rates of exosome-related protein activation and greater SNCA accumulation in males. Additionally, inflammatory marker NLRP3 was downregulated in LuxPark but upregulated in PPMI (Table S7).

##### Axonal transport and neuronal connectivity

Parallel to exosome-related changes, simulations predicted dysregulation in axonal transport pathways. Axonal transport regulators (KLC1, MAP1B) showed increased activity specifically in LuxPark (Table S6). Simulations revealed declines in axonal transport activity notably more severe in males relative to females.

In summary, the Boolean simulations revealed clear sex-specific molecular dysregulation in PD. Males show more mitochondrial impairment, less mitophagy, and reduced axonal transport compared to females. Although males activated oxidative stress and protein-folding responses, these mechanisms did not prevent increased α-synuclein accumulation and exosomal dysfunction, highlighting sex as a crucial factor in PD molecular pathology.

## 3. Methods

### 3.1 Sex-specific clinical parameters of PD cohorts description and clinical characteristics

We identified baseline miRNA expression data from two previously reported cohorts of individuals diagnosed with idiopathic Parkinson’s disease (PD): the Parkinson’s Progression Markers Initiative (PPMI; n = 350)^25^ and the Luxembourg Parkinson’s Study (LuxPark; n = 367)^26^. For PPMI, this subset reflects individuals with quality-controlled miRNA expression data available at baseline. For LuxPark, we used the same filtered PD sample as described in our previous research work^12^, which excluded individuals with known mutations or close familial relationships.

Participants included in this analysis had whole-blood or plasma miRNA profiles available at baseline and were clinically confirmed idiopathic PD cases. All clinical and demographic information reported below was extracted from the original cohort publications and is not derived from our own analyses. Detailed cohort descriptions are available in PPMI cohort study^25^ and LuxPark cohort study^26–28^.

To contextualize expression and modeling analyses, we compared selected demographic and clinical characteristics between the cohorts, including formal sex-stratified comparisons (Table S8). Both cohorts were male-dominated (65.5% in PPMI; 66.5% in LuxPark) but differed in recruitment strategy, disease stage, and treatment status. PPMI participants were newly diagnosed and medication-naïve (Levodopa Equivalent Daily Dose (LEDD) = 0 mg/day), while LuxPark participants were already under dopaminergic therapy (mean LEDD: 500 ± h410 mg/day).

LuxPark participants were older at inclusion (67.5 ± 11.2 years) than those in PPMI (61.7 ± 9.7 years), though age at disease onset was comparable (62.3 ± 11.8 vs. 61.7 ± 9.1 years), noting that this metric reflects age at diagnosis in LuxPark and age at first symptom appearance in PPMI. Disease duration reflected the design differences: PPMI participants were recently diagnosed (0.6 ± 0.5 years), whereas LuxPark patients had a longer disease duration (4.9 ± 5.2 years). Clinical severity was also greater in LuxPark, with higher MDS-UPDRS parts I–III scores and lower Montreal Cognitive Assessment (MoCA) scores compared to PPMI (Table 2). Sex-specific analysis revealed that in LuxPark, females presented with higher non-motor symptom burden (MDS-UPDRS I), more functional impairment (MDS-UPDRS II), and received higher levodopa equivalent doses adjusted for body weight. In contrast, MoCA scores were higher in females in both cohorts, consistent with preserved cognitive function.

**Table 2.** Clinical and sociodemographic characteristics of PPMI and LuxPark participants with baseline miRNA expression data.

| Variable | PPMI (n = 350)<sup>25</sup> | LuxPark (n = 367)<sup>26</sup> |
| --- | --- | --- |
| Sex (male) | 65.5% (277/423) | 66.5% (244/367) |
| Age at assessment (years) | 61.7 ± 9.7 | 67.5 ± 11.2 |
| Age at PD onset* (years) | 61.7 ± 9.1 | 62.3 ± 11.8 |
| Disease duration since diagnosis (years) | 0.6 ± 0.5 | 4.9 ± 5.2 |
| MDS-UPDRS I | 5.6 ± 4.1 | 10.6 ± 7.0 |
| MDS-UPDRS II | 6.1 ± 4.2 | 11.5 ± 8.4 |
| MDS-UPDRS III | 20.9 ± 8.9 | 35.2 ± 16.3 |
| MoCA | 27.1 ± 2.3 | 24.0 ± 4.6 |
| LEDD (mg/day) | 0 | 500 ± 410 |

Notes: Values are presented as mean ± SD or percentages. PPMI cohort participants were de novo and medication-naïve at baseline. \* The age at onset was defined as age at first symptom appearance in PPMI and as age at diagnosis in LuxPark. Abbreviations: MoCA = Montreal Cognitive Assessment; MDS-UPDRS = Movement Disorder Society–Unified Parkinson’s Disease Rating Scale; LEDD = Levodopa Equivalent Daily Dose.

These cohort- and sex-specific differences were considered during downstream modeling and interpretation. The primary focus of this study is cross-sectional. However, prior analyses of longitudinal PPMI follow-up data (+4 years) indicate that females may experience slower progression of motor function, which aligns with delayed dysregulation trajectories observed in our male-parameterized Boolean models. While time-resolved dynamics were not directly simulated, these external trends provide a coherent backdrop for interpreting the modeled compensatory breakdowns.

A summary of overall cohort characteristics is presented in Table 2. Formal sex-stratified comparisons are provided in Table S8, and supporting external evidence from additional studies is compiled in Table S9.

### 3.2 miRNA selection and statistical analysis

#### 3.2.1 Cohort overview

In PPMI, we included baseline samples from 350 idiopathic PD patients (118 females, 232 males) with available and quality-controlled baseline miRNA expression data. All participants underwent targeted genetic screening for major PD-related mutations, including LRRK2, GBA, SNCA, PRKN, and PINK1, using genotyping arrays with confirmatory sequencing for detected variants. PPMI did not exclude carriers from the overall study; however, for this analysis we retained only participants without known pathogenic variants, aligning the inclusion criteria with those applied in LuxPark. Samples were excluded if sequencing data were missing or did not meet quality criteria, such as low read counts or the miRNA being absent/very low in more than half of samples. The miRNA expression data were derived from Illumina HiSeq/NovaSeq sequencing platforms^29,30^.

In LuxPark, we included baseline samples from 367 idiopathic PD patients (116 females, 251 males) who had passed genetic screening using targeted resequencing of GBA1 gene via PacBio and via microarray NeuroChip, and for whom no known PD-causing mutations or close familial relationships (first- to third-degree relatives) were observed. The inclusion and exclusion criteria used were identical to those applied in our previous study with detailed description^12^.

#### 3.2.2 Statistical analysis

For each cohort, differential miRNA expression between males and females was assessed using the R package limma, which fits a linear model to each miRNA and applies a moderated t-test to evaluate the sex coefficient (coded as 0 = female, 1 = male). To account for platform-specific variability (microarray vs. sequencing), we applied variance stabilization^12^, sharing variance information across miRNAs. Effect sizes (regression coefficients) represented the estimated change in miRNA expression in males relative to females, with positive values indicating higher expression levels in males relative to females.

The false discovery rate (FDR) was controlled at 5% using the Benjamini-Hochberg procedure^33^ separately in each cohort.

#### 3.2.3 Filtering by disease context and interaction modeling

To focus on miRNA dysregulation that is specific to PD rather than significantly dysregulated solely based of sex, we incorporated into the analysis the age- and sex-matched healthy controls (HCs) from each respective cohort. The HCs were drawn from the same studies (PPMI and LuxPark) under identical recruitment protocols and were matched at the group level to the PD participants. In total, the PPMI dataset included 173 controls (85 males and 88 females), while the LuxPark dataset included 185 controls (92 males and 93 females).

For each miRNA, we computed the effect of sex separately within PD and HC groups, classifying the direction of effect as positive (higher in males), negative (higher in females), or non-significant. We then exclude shared baseline effects: If a miRNA showed a significant dysregulation associated with sex in both PD and HC groups in the same direction, it was removed from the further analysis.

Only miRNAs that displayed a significant dysregulation associated with sex inPD and having no comparable effects in HCs were retained for downstream modeling. The resulting lists, comprising 26 miRNAs in LuxPark and 56 in PPMI, were then used as inputs for pathway enrichment and Boolean simulations.

We explored upstream regulation of the filtered sex-specific miRNAs. We queried the TransmiR^34^ database for validated transcription factor (TF)-miRNA interactions, retaining only entries supported by literature evidence in humans. Unique TF symbols were mapped to Entrez Gene IDs and used as input for pathway enrichment analyses across Gene Ontology Biological Processes (GO-BP), KEGG, Reactome, and the MSigDB Hallmark (H) and curated (C2) gene sets.

### 3.3 Constructing boolean models from systems biology diagrams

We constructed Boolean models based on system biology diagrams from the Parkinson’s disease map hosted on the MINERVA Platform^35^. The MINERVA Platform provides the capacity to export selected segments of the map, and we refer to these parts as diagrams. The diagrams were selected through a pathway enrichment analysis of the miRNA targets from LuxPark and PPMI datasets using the Parkinson’s disease map^36^. Once significant pathways were identified, we exported these as diagrams in CellDesigner (RRID:SCR_007263) SBML format for modeling^37^. We converted these models into SBML-qual, a module of the SBML standard developed to represent Boolean models ^38^. To translate these diagrams into SBML-qual models, we used CaSQ (CellDesigner as SBML-qual)^39^. Additionally, we transformed the diagrams into the Simple Interaction Format (SIF) with CaSQ, and bnet format to facilitate the interoperability with other tools such as Rmut^40^ and BoolNet^41^.

### 3.4 Parameterisation using miRNA data

To parameterise the constructed models, we estimated the size of effects for each microRNA using the Cohen distance^42^. This measure provides a standardized mean difference between two distinct groups, taking into account the standard deviation. The calculated Cohen distances were used as a proxy for quantifying the biological impact of miRNA expression changes between case and control groups^11,12,43^. The greater the distance, the more likely that miRNA had a substantial role in influencing disease progression dynamics. To make these findings more comprehensible, we converted the Cohen distance into probability values using the Common Language effect size (CL) method^44^. The translated probabilities were assigned to the initial states of the corresponding miRNA targets within the SBML-qual models. These computed probabilities reflect the conditions of various disease subtypes. Simulations with random walks across PBMs were performed with pyMaBoSS to determine the likelihood of the outcomes (phenotypes) observed from these models. Each simulation run produced trajectories reflecting dynamics of disease pathways under different perturbations, giving us insights into the potential outcomes of various molecular interactions within the PD cohorts^11^. This allowed us to examine how certain molecular alterations can affect the likelihood of different disease phenotypes. To identify significant shifts within these simulations, we used a regression technique for detecting multiple change points, as outlined by Lindelov^45^.

### 3.5 Probabilistic boolean model simulation

The selected Boolean Models (BMs) were simulated using the pyMaBoSS framework, a Python API designed for the MaBoSS software for probabilistic Boolean modeling and simulation^46^. Random asynchronous transitions were applied within pyMaBoSS to discover steady states and complex attractors from predefined initial states. Each node (representing a biomolecule) was assigned an initial state determined by the input from the miRNA datasets, with probabilistic rules governing the state transitions reflecting the dynamics of molecular interactions within PD pathology. Nodes without direct miRNA input were initialized at the default pyMaBoSS value of 0.5, reflecting equal probability of activation and inhibition under baseline uncertainty. This approach allowed us to capture both deterministic and stochastic behaviors in disease progression. In the probabilistic Boolean simulation, each condition (i.e. configuration of initial states) was analyzed with 100 iterations and 1000 repetitions to ensure a robust representation of dynamic alterations. A high-performance computing (HPC) facility of the University of Luxembourg, specifically the Aion cluster with Intel Xeon Gold processors (20 cores, 192 GB RAM per node), was used for these simulations (https://hpc.uni.lu).

## 4. Discussion

In this work, we explored the impact of sex-specific miRNAs on molecular mechanisms in PD. The results showed that sex-specific miRNAs may influence mechanisms related to dopamine metabolism, mitochondrial dynamics, immune and inflammatory responses. Further, we demonstrated that the regulatory influences are not uniform across cohorts. This may result in cohort-specific variations that could contribute to the observed heterogeneity in PD progression^11,47^.

Our findings support earlier reports that miRNAs regulate neuroinflammatory responses, mitochondrial functions, and protein aggregation^10,11,12,48^. Many of these regulatory signals differed between males and females, indicating disease specific modulation rather than baseline differences. This pattern is consistent with the evidence that hormonal and immune factors shape molecular responses in PD^7,8^. While miRNAs are known to play a regulatory role in PD related mechanisms, the contribution of sex-specific miRNAs to molecular differences require further investigation^49,50^.

In this study, we identified differentially expressed miRNAs between male and female patients in both LuxPark and PPMI cohorts. Comparison of PD cases with healthy controls identified miRNAs exhibiting sex differences only in the disease state. This refined set is expected to reflect PD-specific molecular mechanisms rather than baseline physiological variation.

Our results identified a set of significantly dysregulated sex-specific miRNAs in PD, many of which are implicated in pathways relevant to neurodegeneration and disease progression. These include mechanisms associated with mitochondrial function, protein handling, inflammation, and synaptic regulation. After a systematic review of the literature relevant to these miRNAs, a consistent pattern of involvement in key molecular processes emerged.

Both miR-7-5p and miR-150-5p exhibited consistent downregulation in males compared to females (Table 1). These miRNAs align with prior studies, including miR-7-5p’s role in α-synuclein regulation in brain^17^ and miR-150-5p’s dysregulation in blood^21^. miR-145-5p and miR-335-3p emerged as key regulators of PD-associated pathways, with miR-145-5p linked to synaptic dysfunction in substantia nigra tissues^51^ and miR-335-3p associated with mitochondrial function^52^. Similarly, miR-423-3p, implicated in prefrontal cortex pathology^19^, displayed male-specific downregulation, and miR-224-5p demonstrated cohort-independent dysregulation (Table 1). These miRNAs have been associated with immune responses^24^ and dopamine receptor modulation^53^, highlighting their role in sex-specific PD mechanisms.

The sections that follow first summarize relevant clinical differences between sexes observed across the studied cohorts. Next, we discuss the physiological relevance of sex hormones, such as estrogen and testosterone, as context for interpreting our molecular findings. Finally, we detail the computationally inferred mechanisms and predicted sex-specific deregulations, integrating these insights with clinical observations.

### Clinical parameters and cohort differences

In both cohorts, males showed lower cognitive scores than females (see Table S8). In LuxPark, females also had lower non-motor (UPDRS I) and functional (UPDRS II) scores (*p* < 0.001), but better cognitive scores and higher weight-adjusted LEDD (*p* = 0.001). In PPMI, where patients were untreated, no sex differences were seen in UPDRS III (*p* = 0.34), and UPDRS I–II were not reported. Considering the age of onset (See Table S8), these data suggest early sex differences in cognitive performance, with broader symptom divergences emerging at later disease stages once treatment is introduced. This pattern is consistent with prior reports that male PD patients experience earlier and more rapid motor decline ^54,55^.

Longitudinal evidence from the Pacific Udall Consortium likewise indicates that males progress more rapidly to MCI (Mild Cognitive Impairment) or dementia than females, consistent with a faster erosion of neuronal resilience^56^. Complementing this, a cross-sectional Nijmegen study reported delayed disease onset and higher striatal dopamine levels in females^57^. These clinical observations underscore the importance of examining underlying physiological and molecular factors, discussed in detail in the following sections.

### Physiological relevance of hormonal factors

The clinical patterns align with previous studies reporting that while testosterone has a role in modulating the dopamine receptor expression and neuroinflammation, it does not provide the same degree of neuroprotection as estrogen^58,59^. Estrogen is known to protect pathways of mitochondrial homeostasis, mitigate oxidative stress and facilitate α-synuclein clearance^60,61^, and males might need additional compensatory mechanisms to handle dysfunction of these pathways^25,62,63^. This vulnerability in males aligns with our enrichment analyses, highlighting dysregulated estrogen-responsive pathways in the absence of hormonal protection (Table S13).

Results of our simulations are consistent with the hypothesis that sex-linked molecular differences shape the robustness of compensatory signaling in PD^64,65^. By this, we refer to the activation of stress-adaptive pathways that attempt to restore cellular homeostasis under sustained molecular dysregulation, such as mitochondrial, oxidative, and proteostatic systems. Both genders exhibit compensatory functions, with males showing sustained activation of these mechanisms, possibly due to limited estrogen^58,59^. These physiological insights provide important context for the sex-specific molecular deregulations predicted by our computational modeling

### Sex-specific pathway dysregulation and compensatory mechanisms

Mitochondrial biogenesis is one the mechanisms predicted to have higher activation in males (Figure 3; Table S1) across both PD cohorts. Simulations indicate increased activity of PGC-1α, a main regulator of mitochondrial biogenesis, and COX5A, a component of mitochondrial complex IV. This aligns with previous observations that biogenesis can be transiently activated in PD to meet energetic demands under oxidative stress^66,67^. However, in the PPMI context, this signal was less clear: activity of COX8A was increased, while COX6A1 and NDUFB8 decreased, suggesting a complex interplay between the electron transport chain components^68–70^.

Mitophagy and pathways compensating oxidative stress also show sex-specific dysregulation. In males, FOXO3-mediated pathways that maintain mitochondrial genome integrity and antioxidant defenses ^71–73,^, were predicted to have increased activity in both PPMI and LuxPark cohorts (Table S2). Correspondingly, TXNIP, SIRT2, and MAPK9 were predicted to increase in males, consistent with roles in oxidative stress responses^74,75^, promoting stress-induced mitochondrial biogenesis^76^, and supporting mitochondrial compensation^77,78^. However, the regulation of mitophagy showed cohort-level differences: the activity of its regulators PRKN and GABARAPL1 increased in the LuxPark, while in PPMI GABARAPL1 activity decreased and PRKN remained unchanged. Some of these discrepancies may reflect not only sex-specific miRNA regulation but also differences in treatment status, given that PPMI participants were untreated while LuxPark patients had ongoing dopaminergic therapy.

Regulation of autophagy differs across cohorts. In LuxPark, predicted activity of HSPA8 and HSPB1 (chaperones facilitating autophagy) was decreased, while the activity of MAP1LC3B was increased (Figure 3; Tables S2-S4). By contrast, in PPMI, HSPA8 activity was predicted to be increased in male patients. Notably, our simulations predict that α-synuclein accumulation increases in males in both cohorts. This aligns with clinical data showing α-synuclein buildup and autophagy failure in PD^79,^, which has been linked to rapid motor and cognitive decline due to proteinopathy^80^.

An increased misfolded protein load in our model is accompanied by activation of the exosomal pathway to support protein clearance^81^, including the removal of α-synuclein from neurons^82^- a process implicated in PD pathology^81^ (Table S5). Key molecules involved in exosome biogenesis and release, such as LGALS3 and PDCD6IP^83^, are predicted to be activated in males across both cohorts. This pattern suggests a compensatory response to alleviate intracellular proteotoxic stress^82,84^, aligning with reports of upregulated exosomal pathways in response to α-synuclein accumulation^82^.

While helping with neuronal protein load, exosomal activity facilitates the intercellular transmission of α-synuclein across neuronal networks^85^. Our simulations indicate that across both cohorts this process is increased in males (Table S5). This correlates with predicted activity of pro-inflammatory molecules, such as IFNG and CD36, which were increased in males in both cohorts. This simulated pattern matches findings of sustained inflammation in advanced PD^86,87^ and persistent immune activation accelerating neuron loss in PD^88^.

This shared pattern supports the view that exosomes, while compensatory at first, may become maladaptive as α-synuclein accumulates^89^. The spread of α-synuclein via exosomes exacerbates neuroinflammation and degeneration, expanding the pathology to previously unaffected brain regions^90^.

Parallel to these exosome-related changes, our model predicts dysregulation in axonal transport pathways. Specifically, we observed increased activity of MAP1B, a cytoskeletal stabilizer involved in axonal integrity and trafficking. Elevated MAP1B activity might represent an early compensatory response aiming to maintain neuronal connectivity under stress conditions. However, chronic or excessive stabilization could negatively influence axonal transport dynamics, potentially disrupting neuronal communication. Such axonal transport dysfunction is known to independently contribute to motor and cognitive deficits observed clinically in PD^91^. Thus, both exosomal transmission of α-synuclein and axonal transport dysregulation represent complementary and convergent pathways exacerbating neuronal dysfunction and disease progression.

### Perspectives and implications

To understand the sex-specific molecular differences in PD progression^92^, it is important to investigate their compensatory pathways. Our simulations show consistent activation of the pathways to maintain mitochondrial biogenesis, mitigate oxidative stress responses, and facilitate α-synuclein clearance. This aligns with the hypothesis that males, due to limited testosterone-mediated neuroprotection, must recruit these mechanisms earlier ^54,55^. However, these mechanisms are ultimately insufficient to control disease progression. In contrast, females benefit from estrogen, which may delay the need for such early activation until its decline in later disease stages ^93,94^.

Our model suggests that although compensatory pathways are activated under male-specific dysregulation, they do not achieve stable recovery. Molecules like UCHL1 and RHEB-mTORC1^95^, LGALS3 and PDCD6IP ^96,97^, and CD36^98^ remain persistently active, potentially reflecting unresolved cellular stress. This pattern points to the early stages of inflammatory signaling, where continued strain and impaired clearance begin to intersect^99^. It marks a shift from temporary adaptation to longer-term vulnerability, consistent with the greater clinical burden observed in males^39^.

Together, these findings support a broader hypothesis: motor symptoms in PD tend to emerge earlier and progress more rapidly in males^54,55^, while females may retain motor function longer but show earlier signs of cognitive disruption^94,100^. In males, cognitive symptoms often follow motor decline more quickly^94,101^. Research studies show that estrogen may contribute to this difference by supporting mitochondrial and redox resilience^102^, though this protection diminishes with age and hormonal decline^9,103^. Males may also be more vulnerable to protein misfolding and α-synuclein aggregation due to the lack of estrogen-mediated buffering^104^. This aligns with reports that PD males exhibit earlier neurostructural aging and are more prone to cognitive and motor decline^105^.

## 5. Limitations

Our study provides valuable insights into sex-specific miRNA regulatory patterns and their implications in PD, but it is also subject to methodological and conceptual considerations. First, the use of cross-sectional miRNA expression data inherently restricts our ability to fully capture dynamic molecular processes occurring over the disease course. Longitudinal studies would be essential to validate temporal relationships and better understand the trajectory of these molecular changes.

Second, differences in technological platforms, microarray for LuxPark and sequencing for PPMI, introduce variability in data generation. Although robust statistical normalization was applied to mitigate this, future studies employing unified sequencing approaches would help reduce platform-specific effects and enhance cross-cohort comparability.

Third, integrating additional layers of omics data (e.g., proteomics, transcriptomics, and metabolomics) in future computational approaches could refine model granularity and biological realism. Finally, inherent biological heterogeneity among patients, including individual differences in disease trajectory, medication status, and molecular responses, represents an ongoing challenge. In particular, all PPMI participants were de novo and drug-naïve at baseline (LEDD = 0), whereas LuxPark patients had already initiated dopaminergic treatment. The pharmacological difference is likely to influence circulating miRNA profiles, as dopaminergic therapy has been shown to affect miRNA expression in PD patients^16^. This effect can contribute to cohort-specific divergence in simulated molecule activity. While the distinction reflects real-world clinical variability, it also represents a limitation when interpreting cross-cohort differences purely as disease-related.

## 6. Conclusions

The study provides evidence for the role of sex-specific miRNAs in PD pathways dysregulation. We showed the molecular differences that shape compensatory mechanisms and disease dysregulation in males compared to females. The Boolean modeling highlights consistent sex-dependent regulatory patterns across pathways essential to mitochondrial biogenesis, mitophagy and oxidative stress responses, and neuroinflammation. The findings align with existing literature and underscore the reproducibility of key miRNA signatures, which consistently show significant modulation in both cohorts. These observations go beyond associations by demonstrating mechanistic predictions of pathway dynamics, offering new insights into how males and females respond to neurodegenerative stress.

Thus, miRNAs provide valuable insights into cross-cohort and sex-specific molecular mechanisms underlying Parkinson’s disease. However, the complexity and multi-target nature of miRNA signatures require computational frameworks, such as the Boolean modeling, to effectively analyse and interpret their functional implications at the level of pathway interactions and dynamics.

Our findings are supported by their cross-cohort consistency and literature validation, though further experimental validation is needed. Future experimental studies should focus on validating the regulatory roles of identified miRNAs through targeted perturbation assays, live-cell imaging, and sex-stratified animal models.. Additionally, longitudinal studies capturing the temporal changes in miRNA expression will be essential to map the progression of compensatory responses and their eventual collapse over time.

From a translational perspective, recognizing sex-specific differences in PD can guide future personalized therapeutic strategies. The computational predictions presented here highlight key molecular pathways that warrant further experimental and clinical validation. Validated pathways could inform targeted monitoring and early interventions. Integrating such insights into digital health platforms may help detect early compensatory changes and inflammatory responses. This approach could enable more precise patient management and improved clinical outcomes.

## Data availability

The LuxPARK clinical dataset (part of the Luxembourg Parkinson’s Study under NCER-PD) used in this work is not publicly available due to internal regulations. Access can be obtained upon reasonable request to the NCER-PD Data and Sample Access Committee. Data from the Parkinson’s Progression Markers Initiative (PPMI) were obtained from the PPMI database (www.ppmi-info.org; RRID: SCR_006431). PPMI data are freely available to qualified researchers via the PPMI website upon registration and acceptance of the PPMI data use agreement and publication policies journals.plos.org.

## Code availability

All analysis and modelling scripts used in this study are available in a public GitLab repository: https://gitlab.com/ahmed.hemedan/hemedan25-Sex-specific-microRNA-regulators-of-PD

## Acknowledgments

The authors acknowledge the Parkinson’s Progression Markers Initiative (PPMI) for providing the data used in this research. PPMI, a public-private partnership, is funded by the Michael J. Fox Foundation for Parkinson’s Research and funding partners https://www.ppmi-info.org/fundingpartners. We also acknowledge the joint effort of the NCER-PD consortium members contributing to the Luxembourg Parkinson’s Study. Data used in the preparation of this manuscript were obtained from the National Centre of Excellence in Research on Parkinson’s Disease (NCER-PD). The National Centre of Excellence in Research on Parkinson’s Disease (NCER-PD) was funded by the Luxembourg National Research Fund (FNR/NCER13/BM/11264123). We would like to thank all participants of the Luxembourg Parkinson’s Study for their important support to our research. Furthermore, we acknowledge the joint effort of the National Centre of Excellence in Research on Parkinson’s Disease (NCER-PD) Consortium members from the partner institutions Luxembourg Centre for Systems Biomedicine, Luxembourg Institute of Health, Centre Hospitalier de Luxembourg, and Laboratoire National de Santé generally contributing to the Luxembourg Parkinson’s Study.

## Author contributions

AAH was responsible for conceptualization, formal analysis and modeling, and writing of the original draft, as well as reviewing and editing. MO was responsible for conceptualization, methodology, and data interpretation, and participated in reviewing and editing. AR contributed statistical analysis together with writing and review. LP provided clinical expertise, supported conceptualization, and contributed to writing and review. PK performed enrichment analysis and contributed to writing and review. EG contributed to writing, review, and editing. RS contributed to writing, reviewing and editing. RK provided clinical expertise, supported conceptualization, and contributed to writing and review. VPS contributed to conceptualization, writing, review, and critical editing, and supervised the project. All authors read and approved the final version of the manuscript. AAH and MO contributed equally and share first authorship.

