## supplemental tables and files for "Sex-specific microRNA regulators of Parkinson’s disease: insights from cohort-stratified simulations of compensatory pathway dynamics": miRNA-TF mapping and enrichment.pdf

### Analysis of sex-specific microRNAs and their associated transcription factors in Parkinson's disease

#### 1. Background - Just to give some context to the document

We investigated how sex affects miRNA expression and the pathways they moderate in PD through differential expression analysis of two PD cohorts. From these differentially expressed miRNAs, we performed pathway enrichment analysis on the PD map and Boolean modelling based on downstream targets of the identified miRNAs.

Sex hormones have been shown to be involved in neuronal homeostasis more directly (e.g. estrogen's neuroprotective effects through mitochondrial resilience; testosterone's effects on dopamine receptor signalling) while also indirectly shaping molecular response through miRNA expression modulation. miRNAs are key post-transcriptional regulators of protein activity and as such have been implicated in PD, though their precise involvement as well sex-specific effects remain obscure.

As a complementary approach to downstream enrichment and modelling, we investigated the upstream transcription factors influencing the expression of the dysregulated miRNAs identified in the PD cohorts. Sex hormones may directly affect the expression and activity of these transcription factors, revealing upstream angles through which sex-specific differences in PD can be described further. We aim at a comprehensive analysis of sex-specific markers of PD through the lens of miRNAs, starting upstream with hormone-mediated TF activity leading to differential miRNA expression, modulation of miRNAs by sex hormones, and finally direct effects of sex hormones on PD-associated pathways.

#### 2. Methodology

We compiled a list of miRNAs determined to be significantly differentially expressed ($p < 0.5$) in both the NCER PD and PPMI cohorts and queried them in the TransmiR database[1] to retrieve known human transcription factors. The TransmiR database contains over 5000 TF-miRNA regulations from more than 2000 publications alongside around 6 million putative TF-miRNA regulations from ChIP-seq data[1]. We opted to use only literature-backed evidence to retain high confidence in our analysis results.
From this query, we obtain 84 transcription factors associated with 15 unique miRNAs selected from our differential expression analysis with an average of 9.6 transcription factors associated with each miRNA. TransmiR also details whether a given TF represses or activates its miRNA target, providing nuanced insights to regulatory mechanisms.

Next, we performed overrepresentation analysis based on the list of matched transcription factors to retrieve the biological processes and pathways these TFs participate in. Such enrichment analyses can provide pertinent insights to sex-based differential effects on selected transcription factors that are reflected in their miRNA targets. Reference gene sets were retrieved from Gene Ontology[2,3] Biological Process, KEGG[4,5], Reactome[6], and the Molecular Signatures Database (MSigDB)[7,8]. For MSigDB, we refined our search by selecting the C2 and H[9] collections encompassing chemical and genetic perturbations, canonical pathways, and hallmark gene sets. Enriched pathways from all sources were filtered for sex-specific keywords which were compiled into summary categories (table 1).

**Table 1 : Enrichment keywords and associated summary categories**

| **Keywords** | **Category** |
|---|---|
| Steroid hormone, sex hormone, steroid, estradiol, cortisol, progesterone | Hormonal signaling |
| Estrogen, ESR1, ESR2 | Estrogen pathway |
| Androgen, testosterone, AR | Androgen pathway |
| PGR | Progesterone pathway |
| NR3C1, NR4A1, GPER, PRLR, steroid receptor, sex hormone receptor | Steroid receptor signaling |
| Neuroendocrine, HPA axis, HPG axis, hypothalamus, pituitary, prolactin | Neuroendocrine axis |
| Dopaminergic, dopamine | Dopaminergic signaling |
| Sex-specific, sex differences | Sex-specific regulation |

To determine whether the selected transcription factors have some relation to sex-specific phenomena, we used BioKB[10], a text-mining based knowledge repository containing biological relations from over 5.5 million scientific publications. We queried BioKB for each TF and determined its sex-specific regulators by querying its incoming relationships for the keywords “testosterone”, “DHT”, “estrogen”, “estradiol”, and “progesterone” to retrieve relevant publications from which we then manually extracted pertinent information.

### 3. Results

**General enrichment results**

Table 2 shows summary statistics on the number of returned pathways by database.

**Table 2 : Enriched pathways by source**

| Database | Significantly enriched pathways | Significantly enriched sex-specific pathways [% of total] |
|---|---|---|
| Gene Ontology Biological Process | 1632 | 31 [6%] |
| KEGG | 128 | 12 [9%] |
| Reactome | 292 | 28 [10%] |
| MSigDB C2 | 1400 | 80 [6%] |
| MSigDB H | 6 | 0 |

Upon inspection of the significantly enriched pathways filtered for sex-specificity, we observe that our filter terms fail to fully distinguish sex-specific processes, returning some general hormonal- or receptor-associated processes. Given that we filter based on the brief description annotated to the pathways, we accept some off-topic matches to retain a sufficiently broad view.

Of the 84 selected transcription factors, 42 are enriched in sex-related pathways. Of these pathways, those associated with hormonal signaling (n = 29), androgen pathways (n = 23), estrogen pathways (n = 16) are linked to the most transcription factors. Neuroendocrine and dopaminergic signaling pathways are only linked to 12 and 9 transcription factors respectively. Transcription factors associated with dysregulated miRNAs as identified in both cohorts are therefore indeed closely linked to sex-specific pathways.

**Effects of sex hormones from transcription factors to associated miRNAs**

We perform a literature search to investigate how sex-related factors may affect expression and activity of the 42 TFs associated with selected pathways.

**Table 3 : Transcription factors associated with sex-specific pathways together with their miRNA targets and compiled sex-specific effects on TFs**

| Transcription factor | Associated processes | Targets | Sex-specific effects on TF |
|---|---|---|---|
| AGO2 | Estrogen Pathway | hsa-mir-150 | ERβ-AGO2 interaction modulates genome activity and miRNA loading in the RISC[11] |
| AKT1 | Neuroendocrine Axis, Estrogen Pathway, Dopaminergic Signaling, Androgen Pathway, Hormonal Signaling | hsa-mir-22 | ERβ represses Akt signaling[12] |
| AKT2 | Neuroendocrine Axis, Estrogen Pathway, Dopaminergic Signaling, Hormonal Signaling | hsa-mir-22 | ERα regulates PI3K/AKT2[13] |
| AKT3 | Neuroendocrine Axis, Estrogen Pathway, Dopaminergic Signaling, Hormonal Signaling | hsa-mir-22 | AKT3 expression may be negatively regulated by estrogen[14] |
| AR | Hormonal Signaling, Estrogen Pathway, Androgen Pathway | hsa-mir-150 | Testosterone and DHT mediate their effects by binding to the AR[15] |
| CFTR | Hormonal Signaling | hsa-mir-193b | CFTR expression is upregulated by estrogen and downregulated by progesterone [16,17] |
| CREB1 | Hormonal Signaling, Androgen Pathway, Estrogen Pathway, Dopaminergic Signaling | hsa-mir-495, hsa-mir-22, hsa-mir-433 | ERα mediates CREB phosphorylation[18], testosterone activates CREB[19] |
| CTNNB1 | Hormonal Signaling, Dopaminergic Signaling, Androgen Pathway | hsa-mir-150 | DHT promotes nuclear accumulation of CTNNB1 which |

| | | | |
|---|---|---|---|
| | | | associates with AR[20,21], estrogen and estradiol enhance up-regulation[22,23]and nuclear accumulation[24] of CTNNB1 |
| DNMT3A | Hormonal Signaling | hsa-mir-124 | Estradiol and dihydrotestosterone reduce DNMT3a mRNA expression[25] |
| E2F1 | Androgen Pathway | hsa-mir-224, hsa-mir-7 | Estrogen induces E2F1 expression and activation[26,27] |
| EGR1 | Hormonal Signaling | hsa-mir-124 | Estrogen induces EGR1[28] |
| EP300 | Hormonal Signaling, Androgen Pathway, Estrogen Pathway | hsa-mir-150 | Testosterone treatment downregulates EP300[29], estrogen indirectly promotes EP300[30] |
| ESR1 | Hormonal Signaling, Estrogen Pathway, Neuroendocrine Axis | hsa-mir-22, hsa-mir-7 | ESR1 is an estrogen receptor through which estrogen exerts its effects[31] |
| ESR2 | Hormonal Signaling, Estrogen Pathway, Neuroendocrine Axis | hsa-mir-7 | ESR1 is an estrogen receptor through which estrogen exerts its effects[32] |
| EZH2 | Hormonal Signaling | hsa-mir-22, hsa-mir-139 | Estradiol induces EZH2 transcription[33]; progesterone leads to increased EZH2 levels[34] |
| FOSB | Hormonal Signaling, Estrogen Pathway | hsa-mir-22 | No sex-specific data |
| HDAC1 | Hormonal Signaling, Androgen Pathway, Estrogen Pathway | hsa-mir-22, hsa-mir-503, hsa-mir-124, hsa-mir-424 | HDAC1 is induced by 17β-estradiol and progesterone[35] |
| HDAC2 | Androgen Pathway | hsa-mir-424, hsa-mir-503 | HDAC2 is induced by 17β-estradiol and progesterone[35] |
| HES1 | Androgen Pathway | hsa-mir-139 | DHT upregulates HES1[36], estradiol and |

| | | | |
|---|---|---|---|
| | | | estrogen decrease HES1 expression[37,38] |
| HIF1A | Dopaminergic Signaling | hsa-mir-424, hsa-mir-433, hsa-mir-150, hsa-mir-224 | DHT inhibits HIF1A production[39], progesterone increases HIF1A transcription[40] |
| JUN | Hormonal Signaling, Estrogen Pathway, Androgen Pathway, Neuroendocrine Axis | hsa-mir-139, hsa-mir-224, hsa-mir-22 | DHT induces expression of JUN[41]; estrogen stimulates transcription of JUN[42] |
| MYC | Estrogen Pathway, Androgen Pathway, Neuroendocrine Axis, Hormonal Signaling | hsa-mir-22, hsa-mir-150, hsa-mir-193b | DHT suppresses MYC expression[43,44], progesterone and estrogen stimulate MYC expression[45–47] |
| MYOD1 | Hormonal Signaling | hsa-mir-22, hsa-mir-133a, hsa-mir-150 | Estrogen[48], progesterone[49], testosterone[48], and DHT[50] positively regulate MYOD1 expression |
| MYOG | Hormonal Signaling | hsa-mir-133a | Testosterone promotes MYOG expression[51,52] |
| NFKB1 | Hormonal Signaling, Neuroendocrine Axis, Estrogen Pathway | hsa-mir-136, hsa-mir-424, hsa-mir-224, hsa-mir-150, hsa-mir-503 | Sex-specific differences in NFKB1 subunit expression[53,54] |
| PRMT5 | Androgen Pathway | hsa-mir-503 | Testosterone induces nuclear depletion of PRMT5[55] |
| PTEN | Androgen Pathway, Hormonal Signaling | hsa-mir-22 | DHT enhances PTEN expression[56]; estrogen inhibits PTEN[57,58], estradiol enhances PTEN phosphorylation[59] while testosterone inhibits PTEN phosphorylation[60] |
| RELA | Hormonal Signaling, Neuroendocrine Axis, Androgen Pathway | hsa-mir-22, hsa-mir-224, hsa-mir-150, hsa-mir-193b, hsa-mir-503 | DHT decreases RELA levels[54], progesterone inhibits RELA[61] |
| REST | Hormonal Signaling | hsa-mir-330, hsa-mir-124, | No sex-specific data |

| | | | |
|---|---|---|---|
| | | hsa-mir-139 | |
| SIRT1 | Hormonal Signaling, Androgen Pathway | hsa-mir-503, hsa-mir-424 | Testosterone, estrone, and estradiol induce SIRT1 expression[62] |
| SMAD2 | Androgen Pathway | hsa-mir-139 | Testosterone downregulates SMAD2[63,64], estrogen decreases levels of phosphorylated SMAD2[65,66] |
| SMAD3 | Androgen Pathway | hsa-mir-136, hsa-mir-424, hsa-mir-433, hsa-mir-503, hsa-mir-139 | Estrogen decreases levels of phosphorylated SMAD3[65,66], DHT indirectly downregulates SMAD3[67] |
| SMAD4 | Androgen Pathway | hsa-mir-424, hsa-mir-503, hsa-mir-139 | DHT increases SMAD4 mRNA levels[68] |
| SNAI1 | Hormonal Signaling | hsa-mir-22 | DHT mediates SNAI1 repression[69], context-dependent effects of estrogen on SNAI1[70,71] |
| SOX2 | Androgen Pathway, Dopaminergic Signaling | hsa-mir-22, hsa-mir-424 | SOX2 is upregulated by the X chromosome-linked gene USP9X[72] |
| SP1 | Estrogen Pathway, Androgen Pathway, Hormonal Signaling | hsa-mir-22, hsa-mir-193b | DHT inhibits SP1 expression[73], estrogen nuclear receptors mediate SP1 activity[74], estrogen negatively regulates SP1 activity[75] |
| SRSF1 | Hormonal Signaling | hsa-mir-7 | No observed sex-specific effects[76] |
| STAT1 | Neuroendocrine Axis, Androgen Pathway | hsa-mir-124, hsa-mir-7 | Progesterone downregulates STAT1[77], estrogen activates STAT1 and upregulates its mRNA expression[78,79], estradiol activates STAT1[80] |

| | | | |
|---|---|---|---|
| STAT3 | Hormonal Signaling, Neuroendocrine Axis, Androgen Pathway, Estrogen Pathway, Dopaminergic Signaling | hsa-mir-22, hsa-mir-124, hsa-mir-150 | Progesterone downregulates STAT3[77], estradiol activates STAT3[80,81] |
| STAT5A | Neuroendocrine Axis | hsa-mir-22 | Progesterone activates STAT5 mRNA and protein expression[82,83], estrogen induces STAT5A[84] |
| TGFB1 | Hormonal Signaling, Dopaminergic Signaling | hsa-mir-424, hsa-mir-433, hsa-mir-503, hsa-mir-224 | DHT reduces gene expression of TGFB1[85], progesterone downregulates TGFB1[86], estrogen increases TGFB1[87], estradiol upregulates TGFB1[88] |
| TP53 | Androgen Pathway | hsa-mir-124, hsa-mir-139, hsa-mir-150, hsa-mir-22, hsa-mir-224, hsa-mir-193b, hsa-mir-424 | Progesterone promotes TP53 activity[89,90], context-dependent effect of estrogen[89,91,92] |

#### 4. Discussion

Transcription factors associated with sex- and disease-specific miRNAs identified in the NCER-PD and PPMI cohorts show associations with sex-specific pathways involved in estrogen, androgen, and hormonal signaling. Furthermore, most of the selected transcription factors are modulated to some degree by sex hormones like estrogen or testosterone, suggesting that the observed miRNA dysregulation is partly due to differential effects of sex on transcription factor activity. Thus, it appears that sex plays key roles in PD pathology stemming from targeted effects on miRNA transcription and miRNA activity modulation leading to PD-relevant differential protein activity.

Oliva et al.[93] characterize a wide range of sex differences in the regulation of the human transcriptome, illustrating how 37% of all genes exhibit some kind of sex-biased expression. The miRNA data at hand is not appropriate to make specific statements on the interactions between sex hormones, transcription factors, and their miRNA targets, and despite the differing contexts in which it was collected, the literature evidence in table 3 suggests that sex hormones play significant roles in regulating these transcription factors and, as a consequence, their miRNA targets.
