## supplemental tables and files for "Sex-specific microRNA regulators of Parkinson’s disease: insights from cohort-stratified simulations of compensatory pathway dynamics": NCER-PPMI Supplementary .pdf

### Reproducible sex-specific markers of Parkinson's disease progression using microRNAs and Boolean modelling

The document comprises a series of structured tables focusing on various molecular pathways associated with Parkinson’s Disease (PD). These tables are designed to provide insights into how specific pathways and functional categories are affected across different patient cohorts. The tables are organized into distinct sections, each addressing key biological processes relevant to PD. Here's an overview:

#### Key sections:

- Dopamine Transcription & Neuroprotection Pathways
- Mitochondrial Function & Dynamics
- Axonal Transport & Neuronal Structure
- Transcriptional Regulation & Developmental Signaling
- Oxidative Stress Response & Metabolic Regulation
- Autophagy, Mitophagy, & Lysosomal Biogenesis
- Protein Folding, Stress Response, & Proteostasis
- Wnt/PI3K/Akt Signaling & Insulin Pathways
- SNCA & NLRP3 Inflammatory Signaling

Each table includes columns that detail:

- **Functional Category**: Specifies the biological process or pathway being analyzed (e.g., neuroprotection, mitochondrial function, autophagy).
- **Predicted Cohort Activity (PPMI and NCER-PD)**: This section compares the activity levels of specific molecules within two patient cohorts (PPMI and NCER-PD). Differences in molecular behavior between these cohorts may indicate varying stages or forms of PD pathology.
  - **PPMI Cohort Activity**: Represents observed molecular activity in the Parkinson’s Progression Markers Initiative (PPMI) cohort.
  - **NCER-PD Cohort Activity**: Shows molecular activity in the Luxembourg-based NCER-PD cohort.
- **Expected Behavior**: Describes the anticipated behavior of these molecules based on existing literature, serving as a reference point for understanding the behavior observed in patient cohorts.
- **References**: Cites studies supporting the expected and observed behaviors, ensuring that findings are grounded in current scientific literature.

#### Comparative analysis:

- The document highlights how the observed activities in both cohorts either align with or deviate from the expected molecular behavior, offering potential insights into PD progression.
- For instance, decreased activity in neuroprotective signaling or increased oxidative stress markers may indicate heightened vulnerability or compensatory mechanisms within PD patients.
- By comparing findings across cohorts, the tables aim to uncover differences that could be tied to factors such as disease subtypes, progression rates, or even sex-specific molecular responses.

### 1. Dopamine Transcription Pathway

#### 1.1. Neuroprotection & Survival

| **Functional Category** | **Molecule** | **PPMI Cohort Activity** | **NCER-PD Cohort Activity** | **Expected Behaviour** | **References** |
|---|---|---|---|---|---|
| Neuroprotection & Survival | **ADCYAP1** | Decreased activity in neuroprotective signaling | Decreased activity, reflecting a vulnerable state | Studies have shown decreased PACAP levels in PD patients, particularly in those not undergoing deep brain stimulation (DBS) therapy, and in the akinetic-rigid subtype of the disease. In animal models, PACAP mRNA expression is reduced in the substantia nigra and other brain regions affected by PD, suggesting a link between PACAP downregulation | [1, 2] |

| | | | | | |
|---|---|---|---|---|---|
| | | | | and disease progression. | |
| Neurotrophic Support | **BDNF** | Increased activity in neurotrophic support | Slightly increased activity, indicating robust neuroprotection | Serum levels of mature BDNF (mBDNF) are significantly lower in PD patients compared to non-PD individuals, suggesting systemic reductions in BDNF expression. However, BDNF levels may increase in later stages of PD, possibly as a compensatory mechanism to mitigate neuronal damage. | [3, 4] |
| Apoptosis Regulation | **CFLAR** | Decreased activity, reduced apoptosis inhibition | Decreased activity, consistent apoptosis regulation | While the direct expression of CFLAR in PD is not extensively documented, it is involved in apoptosis regulation, potentially impacting PD pathology. | [5] |
| Oxidative Stress Defense | **PARK7** | Increased activity in oxidative stress defense | Significantly decreased activity, potentially reduced protection | The expression of PARK7 (DJ-1) is generally downregulated in PD, affecting oxidative stress | [6] |

| | | | | | |
|---|---|---|---|---|---|
| | | | | response and neuronal survival. | |
| Dopaminergic Neuron Survival | **NR4A2** | Stable activity | Increased activity, crucial for dopamine neuron survival | Reduced expression of NR4A2 (Nurr1) in PD patients impacts dopamine neuron maintenance and survival. | [7] |
| Neuroprotective Signaling | **RET** | Stable activity | Increased activity, indicating active neuroprotective signaling | RET expression is downregulated in the substantia nigra of PD patients due to dopaminergic neuron loss rather than direct reduction in transcript levels. | [8, 9] |

1.2. Mitochondrial Function

| **Functional Category** | **Molecule** | **PPMI Cohort Activity** | **NCER-PD Cohort Activity** | **Expected Behaviour** | **References** |
|---|---|---|---|---|---|
| Mitochondrial Function | **COX6A1** | Significantly decreased activity, indicating mitochondrial dysfunction | Stable activity | COX6A1 downregulation reflects mitochondrial dysfunction, a key factor in early-stage PD pathology. | [10, 11] |
| Mitochondrial Function | **COX8A** | Increased activity, compensatory response | Stable activity | COX8A is upregulated in PD models, suggesting compensatory mitochondrial responses to oxidative stress. | [12] |

| | | | | | |
|---|---|---|---|---|---|
| Mitochondrial Function | **COX5A** | Stable activity | Increased activity in mitochondrial function, compensating for stress | Downregulation of COX5A, part of complex IV, is associated with decreased oxidative phosphorylation in PD. | [13, 14] |
| Energy Production | **NDUFB8** | Significantly decreased activity, indicating compromised energy production | Slightly increased activity, though still low, indicating mitochondrial stress | The downregulation of mitochondrial genes, including NDUFB8, highlights metabolic stress in PD neurons. | [15] |

##### 1.3 Axonal Transport & Structure

| Functional Category | Molecule | PPMI Cohort Activity | NCER-PD Cohort Activity | Expected Behaviour | References |
|---|---|---|---|---|---|
| Axonal Transport & Structure | **KLC1** | Increased activity, supporting axonal transport | Increased activity, supporting axonal transport | KLC1 is involved in axonal transport, and its increased activity may support neuronal health in PD, though its specific regulation in PD is less well-defined. | [16] |
| Microtubule Stability | **MAP1B** | Stable activity | Increased activity, supporting microtubule stability | MAP1B expression is generally downregulated in PD, affecting microtubule stability and neuron structure. | [17] |

##### 1.4 Transcriptional Regulation

| Functional Category | Molecule | PPMI Cohort Activity | NCER-PD Cohort Activity | Expected Behaviour | References |
|---|---|---|---|---|---|

| | | | | | |
|---|---|---|---|---|---|
| Developmental Signaling | **EN1** | Significantly decreased activity, suggesting reduced developmental signaling | Increased activity, indicating active developmental signaling | EN1 expression is generally downregulated in PD, affecting neuronal development and survival. | [18] |
| Oxidative Stress Response | **FOXO3** | Increased activity in cellular stress response | Increased activity, impacting oxidative stress response | FOXO3 expression in PD is variable, with evidence of both upregulation and downregulation depending on cellular conditions. | [19, 20] |
| Transcriptional Repression | **NCOR2** | Decreased activity in transcriptional repression | Decreased activity, suggesting reduced repression | Reduced NCOR2 activity in PD is linked to increased miR-132 expression, which negatively impacts neuronal survival pathways. | [21] |

**1.5 Dopamine Synthesis & Regulation**

| **Functional Category** | **Molecule** | **PPMI Cohort Activity** | **NCER-PD Cohort Activity** | **Expected Behaviour** | **References** |
|---|---|---|---|---|---|
| Dopaminergic Pathology | **SNCA** | Increased activity, potential for toxic aggregation | Increased activity, associated with α-synuclein pathology | Elevated SNCA (α-synuclein) expression is linked to aggregation and neurodegeneration in PD. | [22] |

| | | | | | |
|---|---|---|---|---|---|
| Dopamine Synthesis | **TH** | Increased activity in dopamine synthesis | Increased activity, indicating ongoing dopamine production | Elevated TH expression may indicate compensatory dopamine synthesis in response to PD progression. | [23] |
| Dopaminergic Signaling | **RGS6** | Increased activity, modulating dopaminergic signaling | Stable activity | RGS6 is essential for dopamine neuron maintenance, with its loss linked to neuron degeneration and motor deficits. | [24] |

##### 1.6 Protein Folding & Stress Response

| **Functional Category** | **Molecule** | **PPMI Cohort Activity** | **NCER-PD Cohort Activity** | **Expected Behaviour** | **References** |
|---|---|---|---|---|---|
| Chaperone-Mediated Folding | **PIN1** | Stable activity | Increased activity, suggesting response to protein misfolding | PIN1 regulates proteins like tau and α-synuclein, which are crucial for maintaining neuronal structure. | [25] |
| RNA Splicing & Stress Response | **SFPQ** | Stable activity | Increased activity, involved in RNA splicing and stress response | SFPQ aggregation and mislocalization contribute to PD pathology by affecting RNA processing. | [26] |

##### 1.7 Neuroprotection & Survival

| **Functional Category** | **Molecule** | **PPMI Cohort Activity** | **NCER-PD Cohort Activity** | **Expected Behaviour** | **References** |
|---|---|---|---|---|---|

| | | | | | |
|---|---|---|---|---|---|
| Mitochondrial Quality Control | **PINK1** | Stable activity | Low activity | Dysregulation of PINK1 affects mitochondrial quality control, contributing to PD pathology. | [27,28] |
| Mitophagy Regulation | **GABARAPL1** | Increased activity | Decreased activity, reduced autophagy regulation | GABARAPL1 is involved in autophagy pathways, critical for clearing damaged mitochondria in PD. | [29] |

**1.8 Oxidative Stress & Metabolism**

| **Functional Category** | **Molecule** | **PPMI Cohort Activity** | **NCER-PD Cohort Activity** | **Expected Behaviour** | **References** |
|---|---|---|---|---|---|
| Oxidative Stress Defense | **SOD1** | Stable activity | Increased activity, reflecting response to oxidative stress | Overexpression of SOD1 has neuroprotective effects by reducing oxidative damage in PD models. | [30] |
| Mitochondrial Protein Quality Control | **CLPP** | Increased activity | Linked to mitochondrial protein quality control | Suppression of CLPP by α-synuclein leads to mitochondrial damage, contributing to oxidative stress and neurotoxicity. | [31] |
| Energy Production Regulation | **PRKAA1** | Stable activity | Increased activity, associated with energy regulation | PRKAA1 plays a key role in cellular energy balance, with increased activity supporting metabolic stress adaptation in PD. | [32] |

#### 2. Mitochondrial Function Pathway

##### 2.1 Mitochondrial Dynamics & Fission/Fusion

| Functional Category | Molecule | PPMI Cohort Activity | NCER-PD Cohort Activity | Expected Behaviour | References |
|---|---|---|---|---|---|
| Mitochondrial Dynamics | **DNM1L** | Decreased activity, potentially affecting mitochondrial dynamics | Stable activity | DNM1L mutations impact mitochondrial fission and function, essential for neuronal health, though specific PD data is limited. | [33] |
| Mitochondrial Import | **TOMM20** | Stable activity | Increased activity | Increased TOMM20 expression can support mitochondrial import processes, protecting against neurodegeneration in PD models. | [34] |
| Protein Modification | **FOXO3** (Acetylated/Phosphorylated) | Increased activity | Similar role in protein modification | FOXO3 modifications are critical for stress response pathways in PD, affecting apoptosis and oxidative stress pathways. | [35] |

##### 2.2 Nuclear Signaling & Stress Response

| Functional Category | Molecule | PPMI Cohort Activity | NCER-PD Cohort Activity | Expected Behaviour | References |
|---|---|---|---|---|---|

| | | | | | |
|---|---|---|---|---|---|
| Nuclear Signaling | **FOXO3** (Nucleus) | Increased activity, key player in stress response | Increased activity, reflecting role in stress response | FOXO3 expression varies in PD, with activity linked to cellular conditions impacting oxidative stress response. | [36, 37] |
| Autophagy Regulation | **GABARAPL1** (RNA) | Increased activity | Significantly decreased activity, indicating reduced autophagy regulation | GABARAPL1 is essential for autophagy, a pathway involved in removing damaged mitochondria and supporting neuronal health. | [38] |
| Apoptotic Pathway | **FASLG** (RNA) | Increased activity, promoting apoptotic pathways | Stable activity | FASLG regulation in PD is complex, with evidence of both upregulation and downregulation depending on context, impacting cell death. | [39] |

##### 2.3 Mitophagy & Quality Control

| **Functional Category** | **Molecule** | **PPMI Cohort Activity** | **NCER-PD Cohort Activity** | **Expected Behaviour** | **References** |
|---|---|---|---|---|---|
| Mitophagy Regulation | **PINK1/PRKN Complex** | Stable activity | Low activity | Dysregulation of the PINK1/PRKN complex affects mitophagy, leading to the accumulation of damaged mitochondria in PD. | [40] |
| Mitochondrial Quality Control | **TOMM20** (Ubiquitinated) | Stable activity | Increased activity | Enhanced TOMM20 activity helps to maintain mitochondrial import function, which is essential for neuron survival. | [41] |

| | | | | | |
|---|---|---|---|---|---|
| Protein Ubiquitination | **UCHL1/PRKN Complex** | Increased activity | Increased activity | UCHL1 levels are elevated in moderate-stage PD, reflecting stress response and involvement in proteostasis. | [42] |

#### 3. SNCA and NLRP3 Inflammatory Signaling Pathway

##### 3.1 Inflammatory Signaling & Microglial Activation

| Functional Category | Molecule | PPMI Cohort Activity | NCER-PD Cohort Activity | Expected Behaviour | References |
|---|---|---|---|---|---|
| Aggregated Protein Complexes | **SNCA_FCGR2_BPTPN6 Complex** | Increased activity, suggesting complex formation and signaling | Increased activity, suggesting complex formation and signaling | Elevated SNCA levels promote aggregation and signaling cascades, impacting neuroinflammation. | [43] |
| Inflammatory Pathway | **cAMP_NLRP3_Microglia** | Increased activity | Increased activity | Activation of the NLRP3 inflammasome contributes to neuroinflammation and PD progression. | [44] |
| Inflammasome Activation | **NLRP3** | Stable activity | Increased activity | NLRP3 inflammasome activation is observed in both neurons and microglia, contributing to PD pathology. | [45] |

##### 3.2 Phagosome Dynamics & Thioredoxin Signaling

| Functional Category | Molecule | PPMI Cohort Activity | NCER-PD Cohort Activity | Expected Behaviour | References |
|---|---|---|---|---|---|
| Phagosome Dynamics | **SNCA_Phagosome** | Increased activity, reflecting active phagocytosis of α-synuclein | Increased activity, reflecting active phagocytosis of α-synuclein | Higher SNCA mRNA levels in dopaminergic neurons in PD patients contribute to aggregation and clearance efforts. | [46] |
| Thioredoxin Signaling | **Thioredoxin_TXNIP Complex** | Increased activity, indicating response to oxidative stress | Increased activity, indicating response to oxidative stress | TXNIP upregulation in PD is linked to oxidative stress pathways, playing a role in neurodegeneration. | [47] |
| Oxidative Stress Pathway | **TXNIP** | Increased activity, associated with oxidative stress pathways | Increased activity, associated with oxidative stress pathways | TXNIP's role in PD is associated with neurodegenerative processes and oxidative stress response. | |

##### 3.4 Immune Modulation & Exosome Dynamics

| Functional Category | Molecule | PPMI Cohort Activity | NCER-PD Cohort Activity | Expected Behaviour | References |
|---|---|---|---|---|---|
| Immune Modulation | **FCGR2B** | Decreased activity, reflecting reduced Fc receptor signaling | Stable activity | Reduced FcγRIIB signaling may impair microglial phagocytosis, contributing to PD-related neuroinflammation. | [49] |

| | | | | | |
|---|---|---|---|---|---|
| Exosome Formation | **LGALS3** | Decreased activity, suggesting reduced exosome formation | Slightly decreased activity | Galectin-3 is linked to α-synuclein aggregation and exosome dynamics, impacting PD pathology. | [50] |
| Exosome Release | **PDCD6IP** | Decreased activity, indicating potential weakness in exosome release | Decreased activity, consistent with reduced exosome function | Impaired exosome function in PD may affect the clearance of α-synuclein aggregates. | [51] |

#### 4. Wnt/PI3K/Akt Signaling Pathway

##### 4.1 Autophagy & Lysosomal Biogenesis

| Functional Category | Molecule | PPMI Cohort Activity | NCER-PD Cohort Activity | Expected Behaviour | References |
|---|---|---|---|---|---|
| Lysosomal Biogenesis | **TFEB_SNCA Complex** | Increased activity | Increased activity | Elevated levels of SNCA affect TFEB activation, impacting lysosomal biogenesis and autophagic flux in PD. | [52] |
| Autophagic Regulation | **TFEB (Phosphorylated)** | Decreased activity | Decreased activity | Reduced nuclear TFEB levels indicate impaired autophagy regulation, contributing to PD pathology. | [53] |
| Neuronal Survival | **IDE_SNCA Protofibril Complex** | Increased activity | Increased activity | SNCA protofibril formation leads to neuronal stress, impacting cell survival and lysosomal pathways. | [54] |

##### 4.2 Neuronal Health & Insulin Signaling

| Functional Category | Molecule | PPMI Cohort Activity | NCER-PD Cohort Activity | Expected Behaviour | References |
|---|---|---|---|---|---|
| Insulin Signaling | **INSR** | Decreased activity | Decreased activity | Impaired insulin receptor signaling is linked to insulin resistance, which exacerbates neurodegenerative processes in PD. | [55] |
| Insulin Signaling Pathway | **IRS1 (Phosphorylated)** | Stable activity | Increased activity | Increased phosphorylation of IRS1 in PD is associated with impaired insulin signaling and neuroinflammation. | [56] |
| Growth Factor Signaling | **IGF1/IGF1R** | Increased activity | Increased activity | IGF1 signaling plays a protective role in neuronal health, with alterations observed in PD patients. | [57] |

**4.3 Protein Synthesis & Energy Regulation**

| Functional Category | Molecule | PPMI Cohort Activity | NCER-PD Cohort Activity | Expected Behaviour | References |
|---|---|---|---|---|---|
| Protein Synthesis Regulation | **EIF4EBP1 (Phosphorylated)** | Increased activity | Stable activity | Increased phosphorylation of EIF4EBP1 is linked to enhanced protein synthesis pathways, potentially contributing to neurodegenerative processes in PD. | [58] |
| Metabolic Regulation | **SIRT1** | Decreased activity | Decreased activity | SIRT1 plays a role in ameliorating neuroinflammation and oxidative stress, with decreased levels | [59] |

| | | | | | |
|---|---|---|---|---|---|
| | | | | contributing to PD progression. | |
| Stress Response & Metabolism | **PRKAA1 (Phosphorylated)** | Stable activity | Increased activity | PRKAA1 activation is linked to energy regulation, with increased phosphorylation associated with a response to metabolic stress in PD. | [60] |

#### 5. PRKN Pathway

##### 5.1 Mitochondrial Quality Control & Mitophagy

| Functional Category | Molecule | PPMI Cohort Activity | NCER-PD Cohort Activity | Expected Behaviour | References |
|---|---|---|---|---|---|
| Mitochondrial Quality Control | **PARK7 (Mitochondrion)** | Decreased activity | Increased activity | PARK7 downregulation compromises mitochondrial protection and oxidative stress response in PD. | [61] |
| Mitophagy Regulation | **PINK1 (Mitochondrion)** | Stable activity | Low activity | PINK1 downregulation disrupts mitophagy, leading to accumulation of damaged mitochondria. | [62] |
| Mitophagy & Ubiquitination | **PINK1/PRKN Complex** | Stable activity | Low activity | Impaired PINK1/PRKN complex activity affects mitochondrial quality control in PD. | [63] |

##### 5.2 Protein Ubiquitination & Neuronal Health

| Functional Category | Molecule | PPMI Cohort Activity | NCER-PD Cohort Activity | Expected Behaviour | References |
|---|---|---|---|---|---|
| Protein Ubiquitination | **TOMM20 (Ubiquitinated)** | Stable activity | Increased activity | Increased TOMM20 activity supports mitochondrial import, essential for neuronal survival in PD. | [64] |
| Oxidative Stress Defense | **oxidized_PARK7 dimer complex** | Decreased activity | Increased activity | PARK7 downregulation is linked to increased oxidative stress, with potential neuroprotective mechanisms impacted in PD. | [65] |
| Mitophagy | **GABARAPL1 (Mitochondrion)** | Increased activity | Decreased activity | GABARAPL1 plays a role in autophagy pathways, crucial for clearing damaged mitochondria in PD. | [66] |

**5.3 Proteostasis & Neuronal Health**

| Functional Category | Molecule | PPMI Cohort Activity | NCER-PD Cohort Activity | Expected Behaviour | References |
|---|---|---|---|---|---|
| Proteostasis | **UCHL1 (Neuron)** | Increased activity | Increased activity | Elevated UCHL1 levels in moderate-stage PD reflect its involvement in the ubiquitin-proteasome system and neuroprotection. | [67] |
| Mitochondrial Health & Autophagy | **HSPA9** | Increased activity | Stable activity | Downregulation of HSPA9 (mortalin) is linked to mitochondrial dysfunction and | [68] |

| | | | | | |
|---|---|---|---|---|---|
| | | | | reduced stress tolerance in PD. | |
| Mitochondrial Stress Response | **SNCA (Mitochondrion)** | Increased activity | Increased activity | Elevated SNCA levels contribute to mitochondrial stress and are associated with neurodegeneration in PD. | [69] |

##### 5.4 Regulation of mTOR Signaling & Cellular Stress

| Functional Category | Molecule | PPMI Cohort Activity | NCER-PD Cohort Activity | Expected Behaviour | References |
|---|---|---|---|---|---|
| mTOR Signaling Regulation | **DDIT4** | Decreased activity | Stable activity | DDIT4 downregulation is linked to impaired mTOR signaling, which may affect neuronal survival in PD. | [70] |
| Stress Response & Apoptosis | **PHLPP1** | Decreased activity | Decreased activity | PHLPP1 plays a role in apoptosis regulation; its reduced activity may impact cell survival pathways in PD. | [71] |
| Energy Regulation | **PRKAA1 (Phosphorylated)** | Stable activity | Increased activity | PRKAA1 phosphorylation helps maintain energy balance during metabolic stress, supporting neuronal health. | [72] |

##### 5.5 Neuronal Health & Proteostasis

| Functional Category | Molecule | PPMI Cohort Activity | NCER-PD Cohort Activity | Expected Behaviour | References |
|---|---|---|---|---|---|

| | | | | | |
|---|---|---|---|---|---|
| Neuronal Health | **UCHL1 (Neuron)** | Increased activity | Increased activity | Elevated UCHL1 levels in PD patients are associated with neuroprotection and modulation of proteostasis. | [73] |
| Autophagy & Lysosomal Function | **TFEB (Acetylated)** | Decreased activity | Stable activity | Reduced TFEB activity impacts lysosomal biogenesis and autophagic flux, leading to impaired cellular homeostasis in PD. | [74] |
| Protein Modification | **SIRT1** | Decreased activity | Decreased activity | SIRT1 plays a crucial role in stress response and metabolic regulation; its downregulation is linked to neurodegeneration in PD. | [75] |

**5.6 Autophagy & Energy Metabolism**

| **Functional Category** | **Molecule** | **PPMI Cohort Activity** | **NCER-PD Cohort Activity** | **Expected Behaviour** | **References** |
|---|---|---|---|---|---|
| Energy Metabolism Regulation | **PRKAA1** | Stable activity | Increased activity | PRKAA1 activation helps counteract metabolic stress, playing a neuroprotective role in PD by maintaining cellular energy balance. | [76] |
| Lysosomal Biogenesis | **TFEB_TFEB Complex** | Decreased activity | Stable activity | Reduced TFEB nuclear localization leads to impaired autophagy, affecting lysosomal function and protein clearance in PD. | [77] |
| Protein Synthesis | **EIF4EBP1 (Phosphorylated)** | Increased activity | Stable activity | Increased phosphorylation of EIF4EBP1 is linked to enhanced protein synthesis, potentially | [78] |

contributing to cellular stress in PD.

early-stage Parkinson's. *Acta Neuropathologica Communications*. doi: 10.1186/s40478-022-01424-6.

34. **Moors, T., Morella, L., Bertran-Cobo, C., Geut, H., Udayar, V., Timmermans-Huisman, E., Ingrassia, A., Brevé, J. J. P., Bol, J. J., Bonifati, V., Jagasia, R., van de Berg, W. D. J.** (2024). Altered TFEB subcellular localization in nigral neurons. *Acta Neuropathologica*.
35. **Kokotos, A. C., Antoniazzi, M. A., Ryan, T. A.** (2023). Phosphoglycerate kinase as a leverage point in Parkinson's Disease-driven neuronal metabolic deficits. *bioRxiv*. doi: 10.1101/2023.10.10.561760.
36. **Tan, A. S., Ng, E. Y. J., Lu, Z., Chia, N. S. Y., Xu, Z., Tay, K. Y., Tan, L. C. S., Tan, E.-K.** (2020). Plasma ubiquitin C-terminal hydrolase L1 levels reflect disease stage and motor severity in Parkinson's disease. *Aging*. doi: 10.18632/AGING.102695.
37. **Lian, B., Zhang, J., Yin, X., Wang, L., Ju, Q., Wang, Y., Jiang, Y., Liu, X., Chen, Y., Li, D., Sun, C.** (2024). SIRT1 improves lactate homeostasis in the brain to alleviate parkinsonism via deacetylation and inhibition of PKM2. *Cell Reports Medicine*. doi: 10.1016/j.xcrm.2024.101684.
38. **Dorion, M.-F., Yaqubi, M., Senkevich, K., MacDonald, N. W., Chen, C. X.-Q., Wallis, A., Antel, J. P., Fon, E. A., Durcan, T. M.** (2023). MerTK is a mediator of alpha-synuclein fibril uptake by human microglia. *Brain*. doi: 10.1093/brain/awad298.
39. **Chou, S. Y., Chan, L., Chung, C. C., Chiu, J. Y., Hsieh, Y. C., Hong, C. T.** (2020). Altered Insulin Receptor Substrate 1 phosphorylation in blood neuron-derived extracellular vesicles from patients with Parkinson's disease. *Frontiers in Cell and Developmental Biology*. doi: 10.3389/fcell.2020.564641.
40. **Gupta, A., Dey, C. S.** (2012). PTEN, a widely known negative regulator of insulin/PI3K signaling, positively regulates neuronal insulin resistance. *Molecular Biology of the Cell*. doi: 10.1091/MBC.E12-05-0337.
41. **Shi, X., Zheng, J., Ma, J., Li, D., Gu, Q., Chen, S., Wang, Z., Sun, W., Li, M.** (2022). Correlation between serum IGF-1 and EGF levels and neuropsychiatric and cognitive aspects in Parkinson's disease patients. *Neurological Sciences*. doi: 10.1007/s10072-022-06490-1.
42. **Cui, X., He, Z., Liu, J., Yan, J., Hua, L.** (2020). MiR-302b-5p enhances the neuroprotective effect of IGF-1 in Parkinson's disease models. *Cell Biochemistry and Function*. doi: 10.1002/CBF.3534.
43. **Pensalfini, A., Jiang, Y., Kim, S., Nixon, R. A.** (2021). Assessing Rab5 activation in neurodegenerative diseases. *Methods of Molecular Biology*. doi: 10.1007/978-1-0716-1346-7_20.
44. **García-Revilla, J., Jin, Y., Boza-Serrano, A., Venero, J. L.** (2023). Galectin-3 shapes toxic alpha-synuclein strains in Parkinson's disease. *Acta Neuropathologica*. doi: 10.1007/s00401-023-02585-x.
45. **Nasoohi, S., Ismael, S., Ishrat, T.** (2018). Thioredoxin-Interacting Protein (TXNIP) in cerebrovascular and neurodegenerative diseases: Regulation and implication. *Molecular Neurobiology*. doi: 10.1007/s12035-018-0917-z.
46. **Sanphui, P., Das, A. K., Biswas, S. C.** (2020). Forkhead Box O3a requires BAF57, a subunit of chromatin remodeler SWI/SNF complex, for induction of p53 up-regulated modulator of apoptosis (Puma) in a model of Parkinson's disease. *Journal of Neurochemistry*. doi: 10.1111/JNC.14969.
