## supplemental tables and files for "Sex-specific microRNA regulators of Parkinson’s disease: insights from cohort-stratified simulations of compensatory pathway dynamics": Results- graphs .pdf

### Reproducible sex-specific markers of Parkinson's disease progression using microRNAs and Boolean modelling

The study investigates the role of sex-specific miRNAs in Parkinson’s Disease (PD) progression. To do so, we apply Boolean modeling to two independent cohorts, PPMI and NCER-PD, aiming to identify consistent and unique patterns in PD-related pathways. Differential expression analysis reveals sex-based differences in miRNA expression in males compared to females. Results demonstrate that these miRNAs modulate pathways such as dopamine metabolism, mitochondrial dynamics, and immune responses differently in males compared to females. Boolean simulations suggest that early compensatory mechanisms differ by sex, with males exhibiting increased compensatory responses in mitochondrial function and oxidative stress management. As compensatory mechanisms become weak, oxidative stress, protein misfolding, and neuroinflammation increasingly drive disease progression, with divergent effects on motor and cognitive symptoms between sexes. The findings indicate the need for sex-informed therapeutic strategies in PD, highlighting sex-specific miRNA modulation as a critical factor in disease progression and potential treatment targets.

### 1. Dopamine transcription pathway:

#### Axonal transport

Both cohorts show an initial increase followed by a decline in axonal transport activity. The increased activity is synchronised with an increased activity of KLC1 gene- A key player in vesical transport process, which maintan synaptic communication (Berth & Lloyd, 2023). In NCER-PD, the initial increase is further synchronised with MAP1B gene- A microtubeule- stabilizing protein that maintain axonal integrity (Wang et al., 2022). The activation of KLC1 and MAP1B may suggest that both cohorts initially employ compensatory mechanisms, sustaining the synaptic function during the PD progression.

However, the subsequent decline in axonal transport activity reflects the progressive nature of PD wherein the compensatory mechanisms begin to fail. In PPMI, the decline synchronises with decreased activity of COX6A1 and NDUFB8 which are critical molecules for mitochondrial function (Berth & Lloyd, 2023). The dysregulation of mitochondrial function can lead to deficits in energy production, affecting the transport process efficiency (Zhang et al., 2023. In NCER-PD, the impairment of the axonal transport is associated with the increased activity of COX5A which regulate mitochondrial respiration (Zhang et al., 2023).

Both cohorts initially manage to maintain the axonal transport, however, the undelying mitochondrial stress can ultimately hinder the associated compensatory roles of the dysregulated molecules.

#### Dopamine metabolism

Dopamine metabolism shows an initial decline in dopamine activity levels in both cohorts which is followed by a stable increase of its activty. The increase in dopamine metabolism is synchronised with the increased activity of BDNF, TH, and SNCA in both cohorts. BDNF is important to maintain the survival of the dopaminergic neurons, while TH is crucial for dopamine synthesis, and SNCA regulates synaptic vesicle related processes (Bae et al., 2019; Poston et al., 2016).

The consistent modulation of dopamine metabolism in both cohorts suggests that the key mechanisms for dopamine synthesis and regulations are conserved. The increase of dopamine metabolism in both cohorts may suggest early compensatory mechanisms, counteracting the neuronal loss. This response aligns with studies showing that a rapid decline in dopamine levels is followed by stabilisation which reflects a negative exponential pattern in disease progression (Jastrzębowska et al., 2019; Shine et al., 2018; Poston et al., 2016).

In the NCER-PD cohort, NR4A2 enhances the transcription of dopamine-related genes and maintains neuron survival (Bae et al., 2019; Poston et al., 2016). This may suggest a resilient dopaminergic system in this cohort. This resilience may provide a protective buffer against early stages of the PD, and highlight the importance of protective

#### Mitochondrial Biogenesis

Mitochondrial biogenesis, a critical process for high energy demanding cells such as neurons, shows an increased activity in both cohorts. This increase indicates an adaptive response to the mitochondrial dysfunction, which often resulted in neurodegenerative conditions.

In the PPMI cohort, the increased activity of mitochondrial biogenesis is synchronised with the activity of COX8A. However, the activity of COX6A1 and NDUFB8, which are critical for effective electron transport and ATP production, has decreased (Peng et al., 2015). This suggests that while there is a response to adapt the mitochondrial deficiency through increased COX8A activity, the overall mitochondrial function remains impaired. In turn, this can lead to decreased energy production over time (Heidorn-Czarna et al., 2021).

In the NCER cohort, the increased activity of mitochondrial biogenesis is observed along with the increased activity of COX5A, ENI, PARK7 and PGC-1α. These molecules are crucial to sustain energy production, indicating a more robust compensatory mechanism in response to mitochondrial dysfunction (Chandra et al., 2017; , Chuang et al., 2019,  Zhou et al., 2021).

Mitochondrial dysfunction has been identified to play a significant role in the pathogenesis of PD, because of energy deficiencies and enhanced oxidative stress, further accelerating neurodegeneration(Yan et al., 2013; Winklhofer & Haass, 2010). The FOXO3 signaling pathway controls  mitochondrial function by regulating the expression of mitochondrial genes while assuring protein quality control.

The modeling results showed an increase in mitochondrial biogenesis in both NCER-PD and PPMI cohorts. Such a trend might indicate that mitochondrial biogenesis gets actively upregulated as a compensatory response to the increased energetic demand and oxidative stress associated with PD (Compagnoni et al., 2020; Chen et al., 2021). This includes resident proteins of the mitochondrial matrix, such as MAPK9 (JNK2) and FOXO3. For example, one report demonstrated that MAPK9 may play a role in facilitating stress-induced mitochondrial biogenesis, a feature that may increase mitochondrial activities observed in these cohorts(Misko et al., 2012; Motori et al., 2020). Meanwhile, FOXO3 regulates the genes that participate in mitochondrial DNA replication and repair activities, critical for the maintenance of mitochondrial integrity under conditions of environmental or metabolic stress(Li et al., 2017; Gao & Zhang, 2018).

Higher activities of mitochondrial biogenesis in the NCER-PD cohort may reflect a more active mitochondrial response, hence offering better resistance to the mitochondrial dysfunction. This might delay severe symptom onset by sustaining ATP production and further mitigating the accumulation of ROS, one of the key players in neurodegeneration(Yan et al., 2013; Wang et al., 2019).

The increasing trend of mitochondrial biogenesis underlines the importance of the FOXO3 pathway supporting mitochondrial function in PD. The increased activity of FOXO3, especially within the mitochondrial matrix, will enhance mitochondrial resilience by reduced neuronal vulnerability to degeneration.

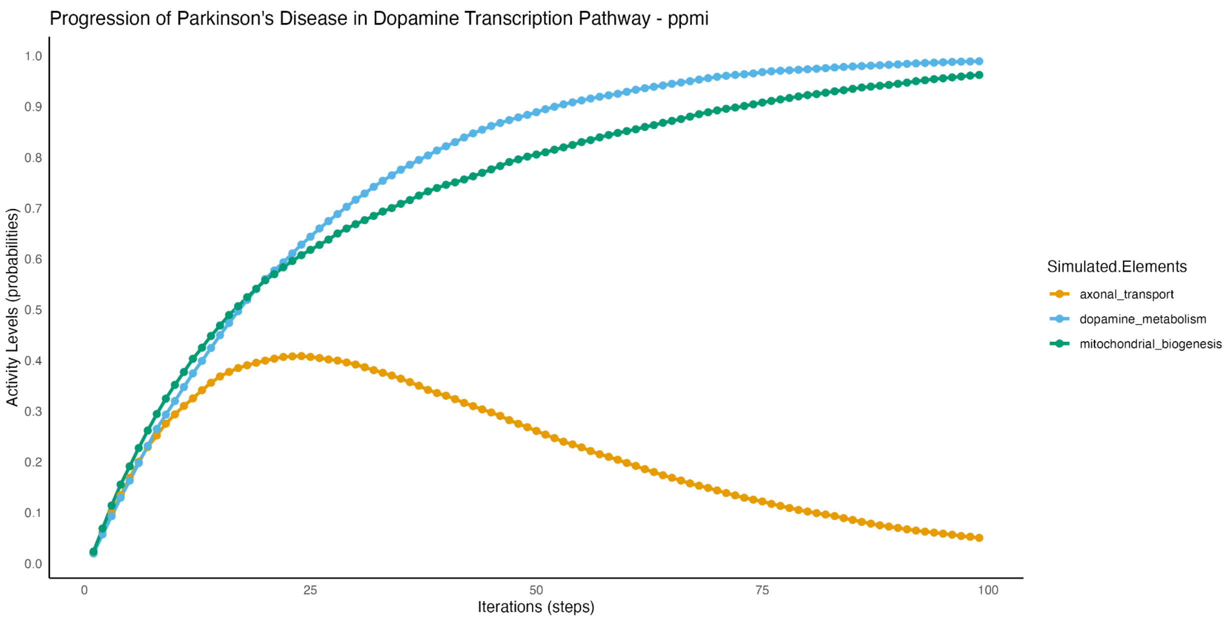


**Figure 1**: Simulation of Parkinson's Disease progression in the dopamine transcription pathway for the PPMI cohort. This graph illustrates how the activity levels of three critical elements—axonal transport (orange), dopamine metabolism (blue), and mitochondrial biogenesis (green) evolve over 100 simulation iterations. The results highlight that dopamine metabolism and mitochondrial biogenesis increase in activity over time, whereas axonal transport shows a decline, suggesting differential regulatory dynamics that could influence disease progression in this cohort..

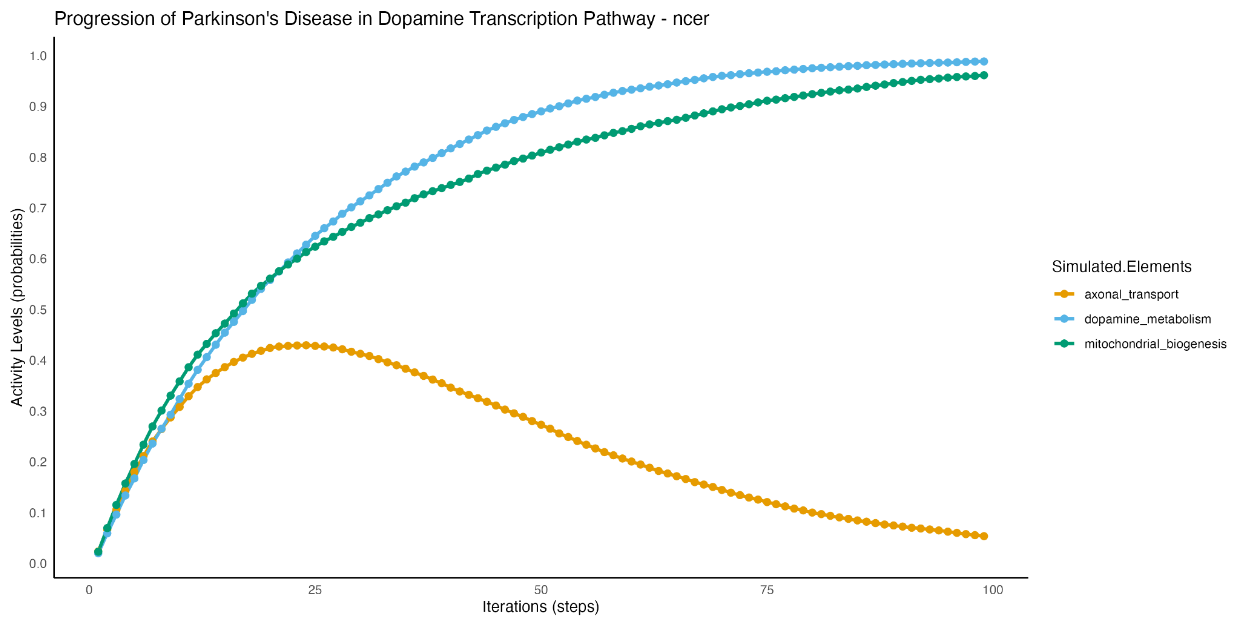


**Figure 2**: Simulation of Parkinson's Disease progression in the dopamine transcription pathway for the NCER cohort. The graph tracks changes in activity levels for axonal transport (orange), dopamine metabolism (blue), and mitochondrial biogenesis (green) across 100 iterations. The results reveal a similar increasing trend in dopamine metabolism and mitochondrial biogenesis, while axonal transport declines, indicating pathway disruptions specific to the NCER cohort. Comparing this with the PPMI cohort highlights potential cohort-specific regulatory mechanisms affecting these pathways.

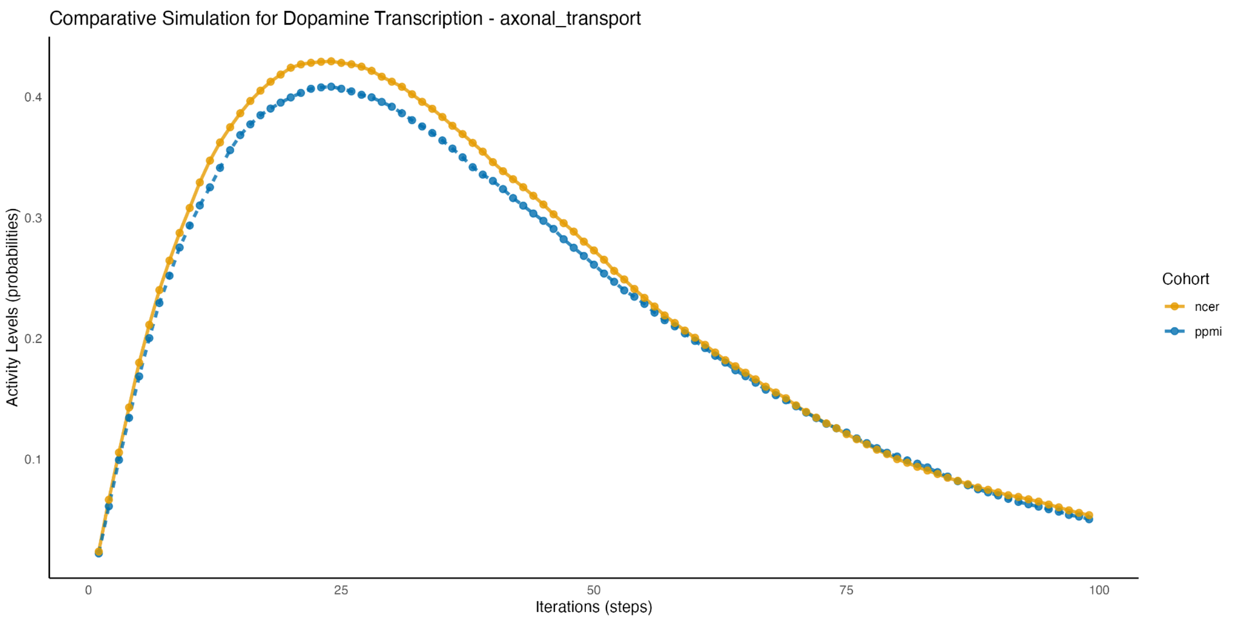


**Figure 3**: Comparative simulation analysis focusing on axonal transport within the dopamine transcription pathway between the PPMI (blue) and NCER (orange) cohorts. The plot demonstrates variations in axonal transport activity over 100 iterations, emphasizing how different cohort characteristics may contribute to distinct pathway disruptions. This comparison suggests that variations in axonal transport may underlie sex-specific or cohort-dependent differences in Parkinson's Disease progression.

### Chaperone-Mediated Protein Folding & Synucleinopathies

Maintaining cellular homeostasis is a critical process that helps to mitigate the accumulation of the misfolded proteins and oxidative stress. While differences in driver biomolecules and their cellular locations between the cohorts, the overall behavior of the pathways remains consistent. This indicates a robust and conserved response to neurodegenerative stress (Koon (2019), Ciechanover & Kwon, 2017).

In the NCER-PD cohort, cytoplasmic chaperone mediated proteins , such as HSPA8 and HSPB1, show a decreased activity, indicating a decreased capacity of folding misfolded proteins. However, this decrease is compensated by an increased activity of autophagic and lysosomal related molecules such as GABARAPL1 and MAP1LC3B acetylated (Su et al., 2020; , Magalhaes et al., 2016).

Further, the elevated activity of SCARB2 in both the endoplasmic reticulum and lysosomes supports that the NCER cohort may compensates for reduced chaperone activity by enhancing lysosomal degradation, specifically targeting misfolded proteins such as α-synuclein (Magalhaes et al., 2016).

In the PPMI cohort, the activity of the HSPA8 increased, indicating a more robust chaperone mediated response than autophagy to maintain the protein homeostasis.

Both cohorts show significant activity in lysosomal degradation of α-synuclein, a hallmark of PD pathology (Ciechanover & Kwon, 2017; , Baughman et al., 2018).

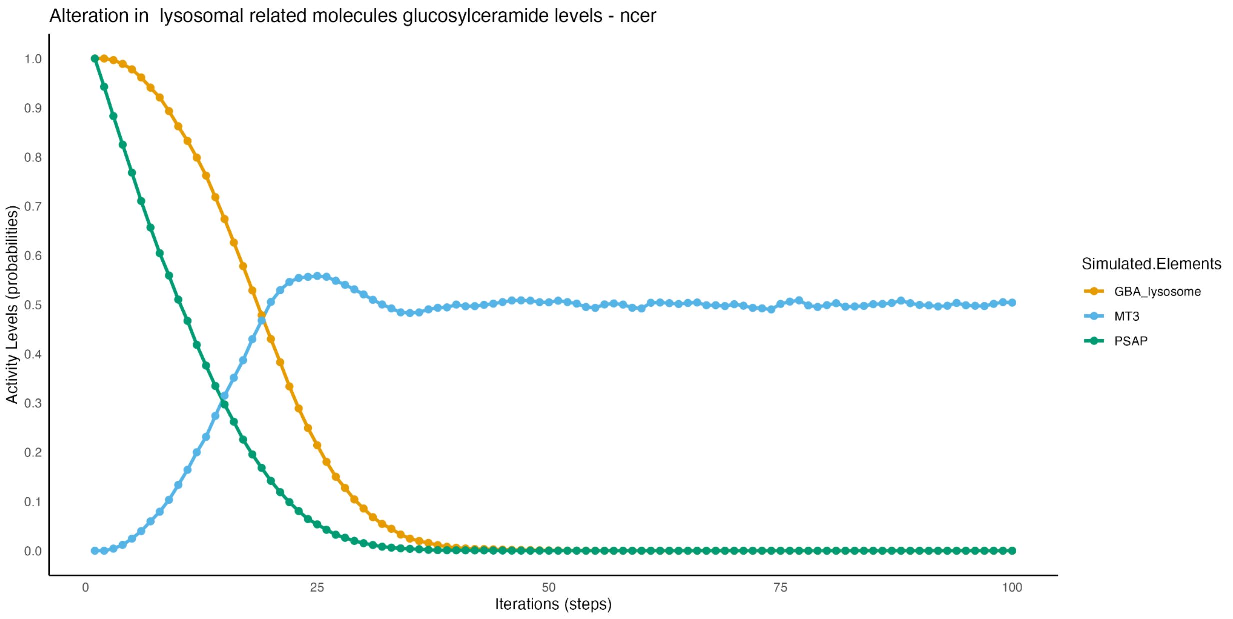


**Figure 4**: Simulation results showing alterations in lysosomal-related molecules (glucosylceramide levels) in the NCER cohort. The graph displays the dynamic activity levels of GBA, MT3, and PSAP over 100 iterations. The simulation indicates that while GBA and MT3 exhibit a rapid decline, PSAP maintains steady levels, suggesting specific disruptions in the lysosomal pathways that could influence disease progression in this cohort.

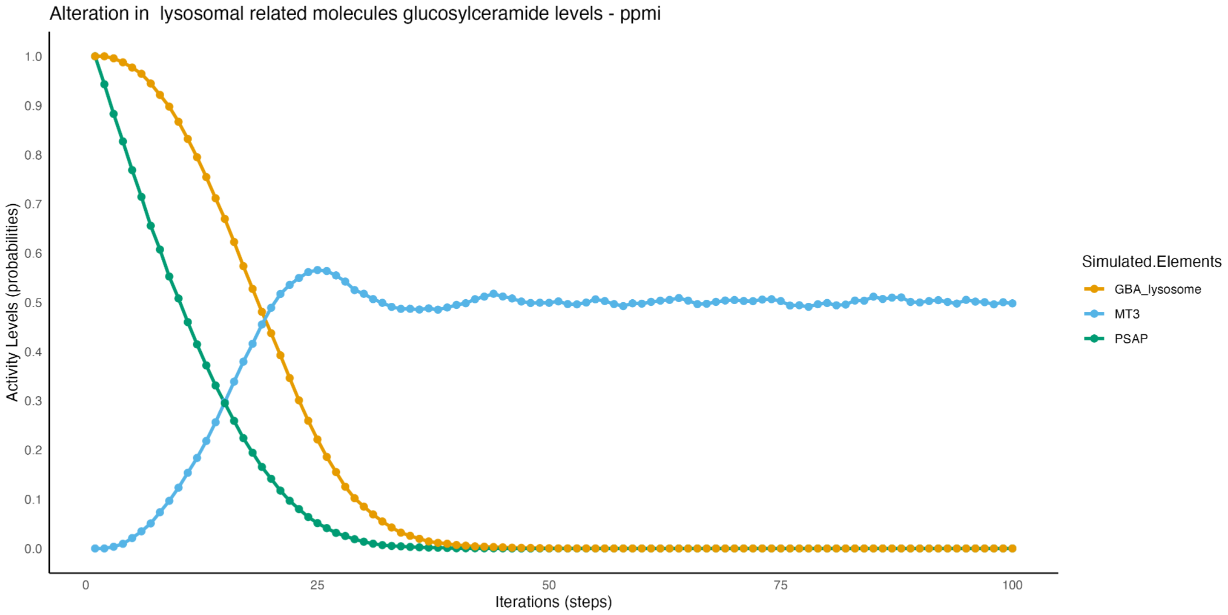


**Figure 5**: Simulation results for the PPMI cohort depicting changes in lysosomal-related molecules (glucosylceramide levels). Activity levels of GBA, MT3, and PSAP are tracked over 100 iterations. The results highlight a marked decline in GBA levels, with MT3 also decreasing, whereas PSAP remains relatively stable, indicating differential regulatory dynamics between the PPMI and NCER cohorts.

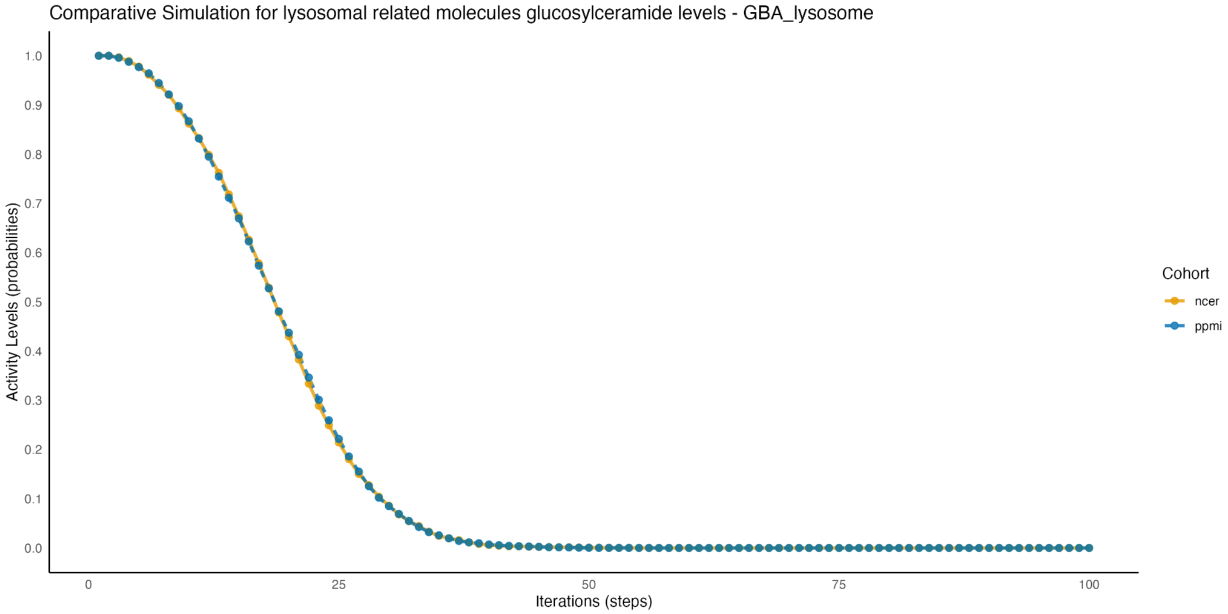

**Figure 6**: Comparative simulation focusing on alterations in GBA activity levels between the NCER and PPMI cohorts. The analysis shows consistent cohort-specific patterns, with the NCER.

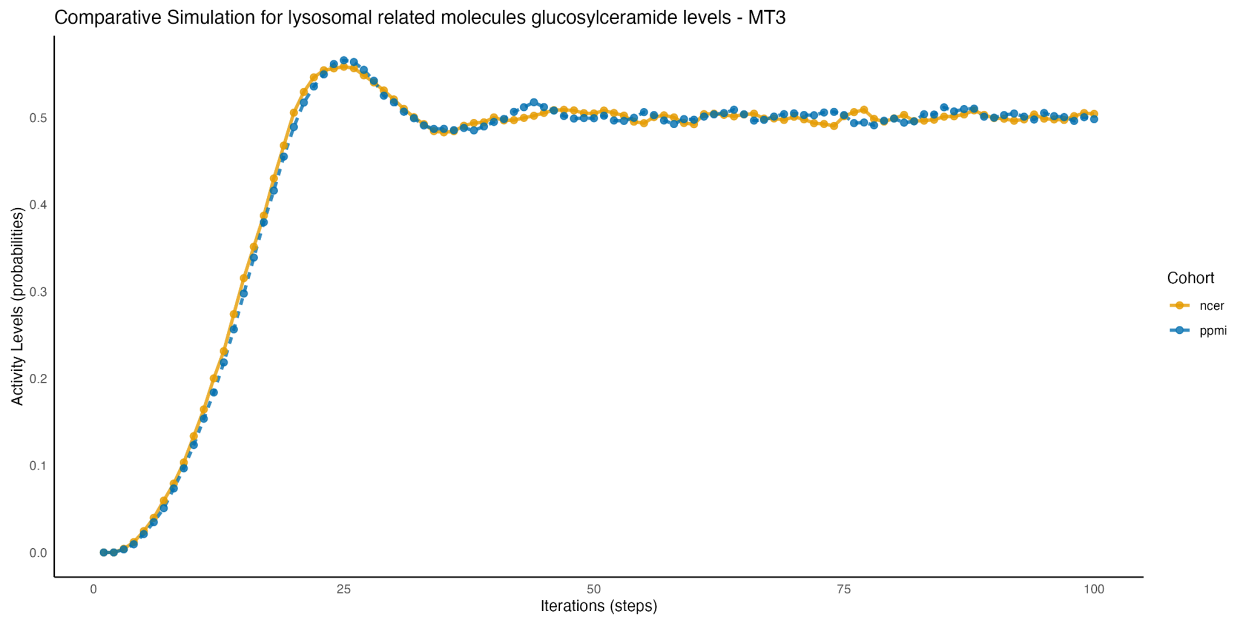


**Figure 7**: Comparative analysis of MT3 activity levels in the lysosomal pathway between the NCER and PPMI cohorts. The simulation reveals consistent in MT3 regulation.

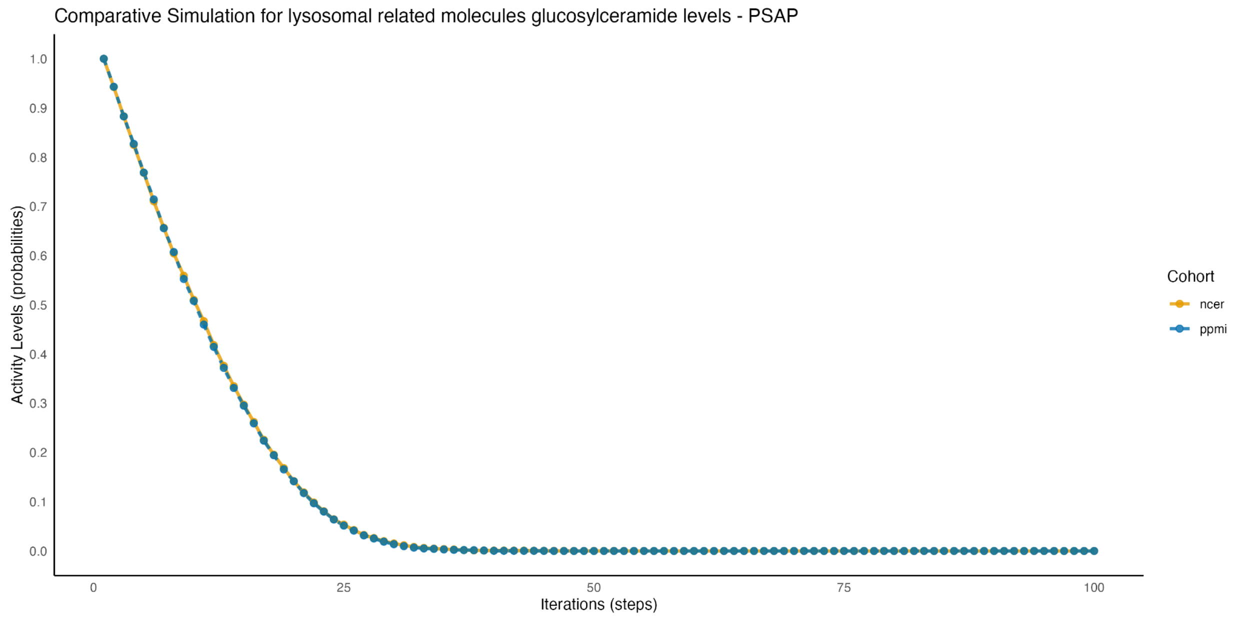

**Figure 8**: Comparative simulation of PSAP activity levels across the NCER and PPMI cohorts. PSAP levels remain consistent between cohorts, suggesting a conserved regulatory mechanism for this lysosomal component that may not be as susceptible to cohort-specific variables.

### FOXO3 Pathway & Oxidative Stress

FOXO3 plays a core role in cellular response against oxidative stress through the main regulation of antioxidant mechanisms. The results indicate that activities of the oxidative stress response in both cohorts have consistently increased, while there is a slight increase in activity level in the NCER-PD cohort. This relates to the heightened activities of TXNIP and SIRT2, important regulators in managing oxidative stress.

TXNIP has become recognized as an important player in cellular redox balance, a protein inducing oxidative stress by repressing antioxidant function from thioredoxin; this protein increases apoptosis upon ROS response. This hypothesis was further supported by Cui et al., 2019. Similarly, the sustained TXNIP activity across the two groups supported the consistent response towards oxidative stress, which is important in mitigating damage that ROS triggers, accelerating neurodegeneration in PD. SIRT2 has been shown to interfere with the cellular responses of oxidative stress, while increased activity was also driven by the involvement of the FOXO3 pathway in managing oxidative lesions.

The higher oxidative stress response in the NCER-PD indicates the involvement of a more robust mechanism of defense, thereby causing a delay in the onset of the oxidative damage. This hypothesis is based on  enhancement of FOXO3-mediated pathways would fortify the cell's defenses against ROS, reducing oxidative damage to neurons.

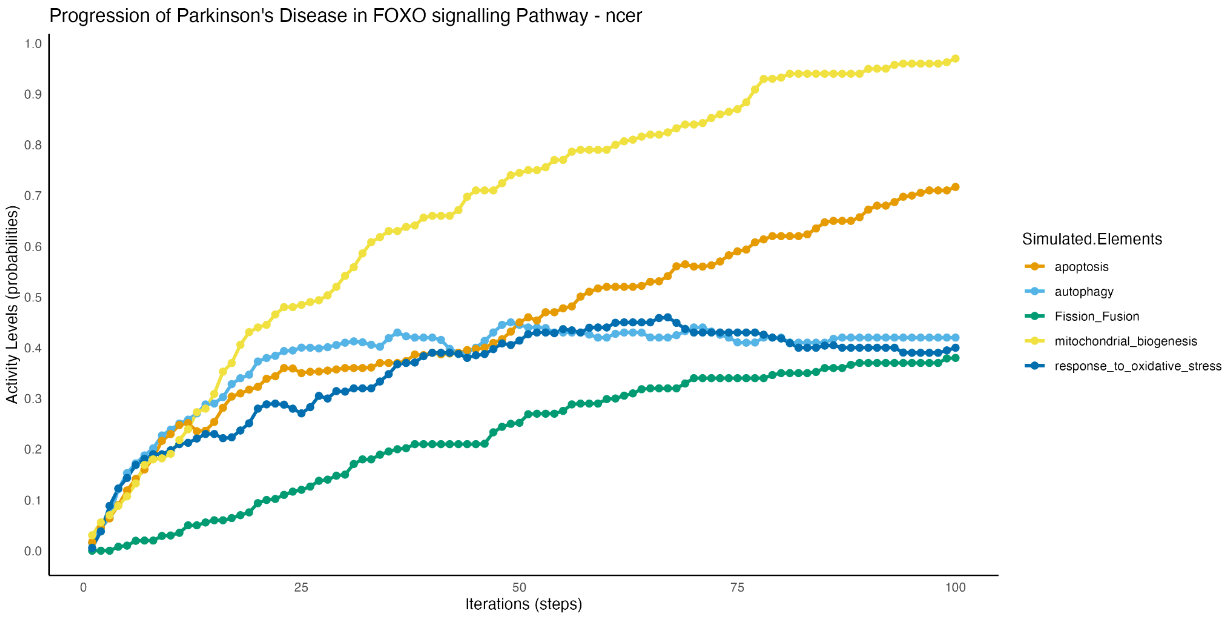


**Figure 9**: Simulation of Parkinson's Disease progression in the FOXO signaling pathway for the NCER cohort. The graph tracks activity levels over 100 iterations for key elements: apoptosis, autophagy, fusion/fission dynamics, mitochondrial biogenesis, and response to oxidative stress. The results reveal distinct temporal patterns, with apoptosis and autophagy showing continuous activation, while the response to oxidative stress remains stable.

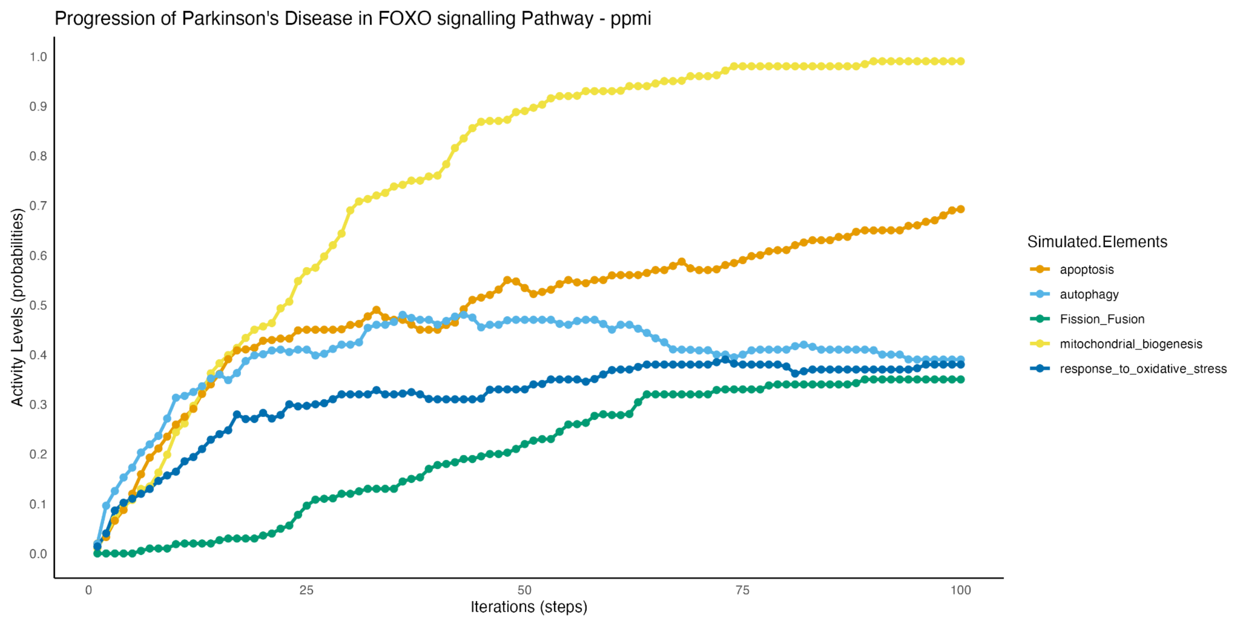


**Figure 10**: Simulation of Parkinson's Disease progression in the FOXO signaling pathway for the PPMI cohort. The plot illustrates how the activity levels of apoptosis, autophagy, fusion/fission dynamics, mitochondrial biogenesis, and response to oxidative stress change over time. The results highlight differences compared to the NCER cohort, suggesting cohort-specific variations in FOXO pathway regulation.

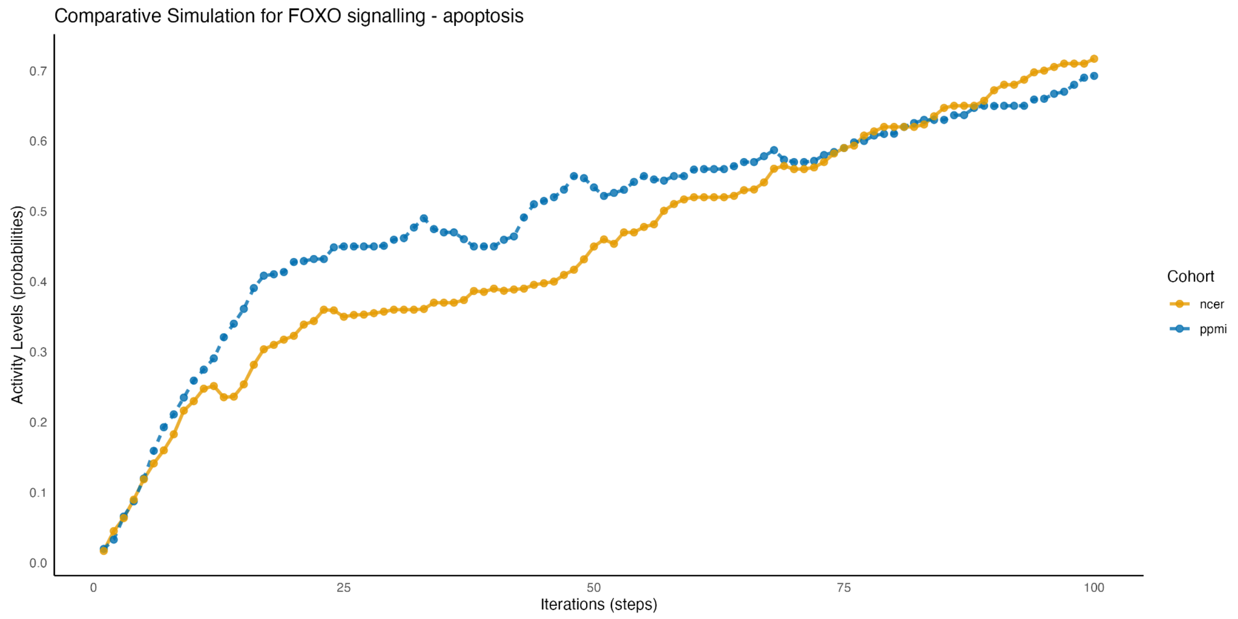

**Figure 11**: Comparative simulation analysis focusing on apoptosis activity within the FOXO signaling pathway across the NCER and PPMI cohorts. The simulation highlights differences in apoptosis activation patterns, indicating that cohort-specific factors may influence the apoptotic response, potentially impacting Parkinson's Disease progression.

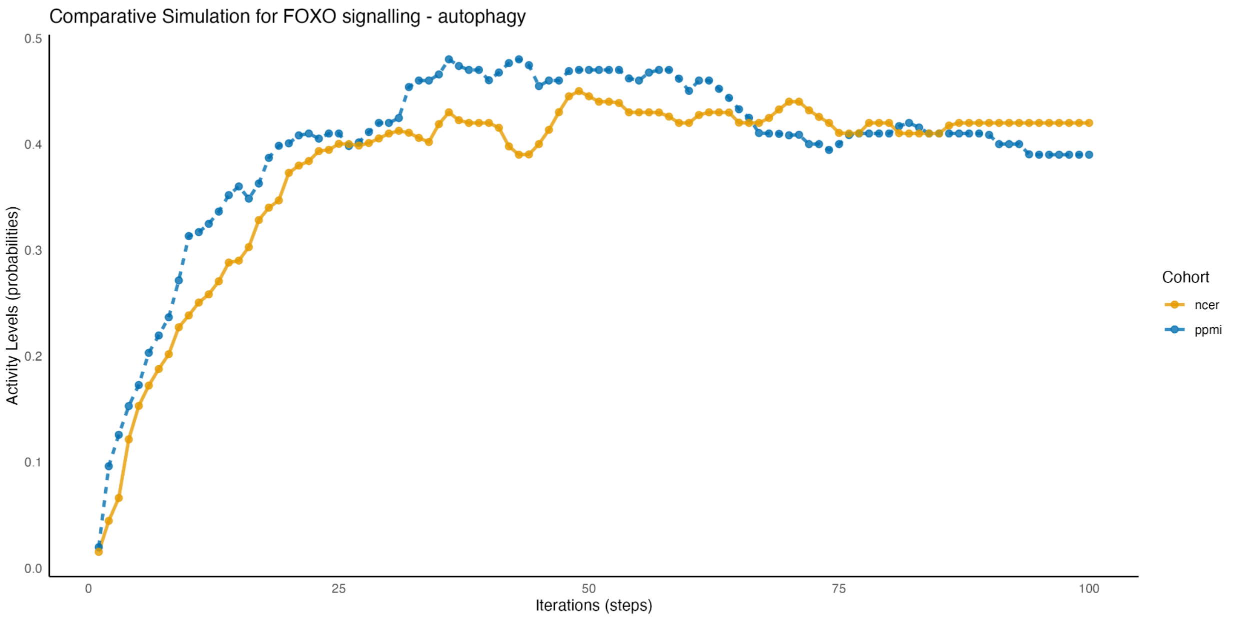


**Figure 12**: Comparative simulation of autophagy activity within the FOXO signaling pathway across the NCER and PPMI cohorts. The simulation results demonstrate distinct activity patterns, with the NCER cohort showing higher and sustained autophagy levels compared to PPMI. This difference may indicate cohort-specific regulation of autophagy processes in Parkinson's Disease.

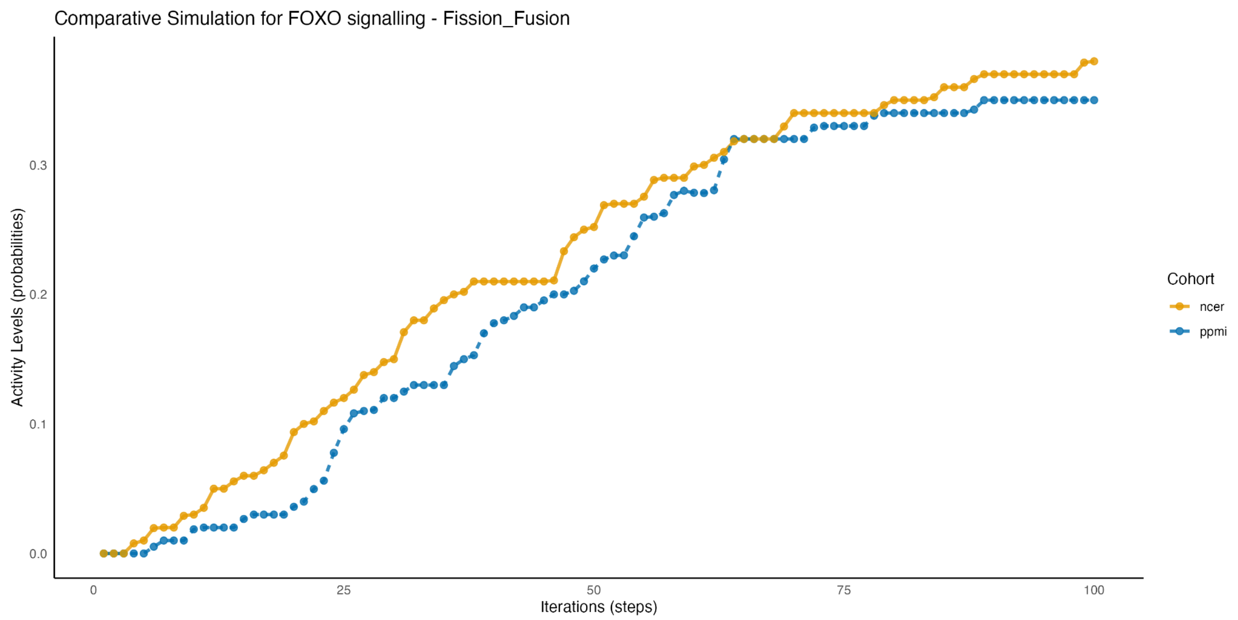


**Figure 13**: Comparative simulation of fusion/fission dynamics in the FOXO signaling pathway for the NCER and PPMI cohorts. The plot highlights differences in mitochondrial fusion/fission activity, suggesting variations in mitochondrial dynamics that could contribute to cohort-specific differences in disease progression.

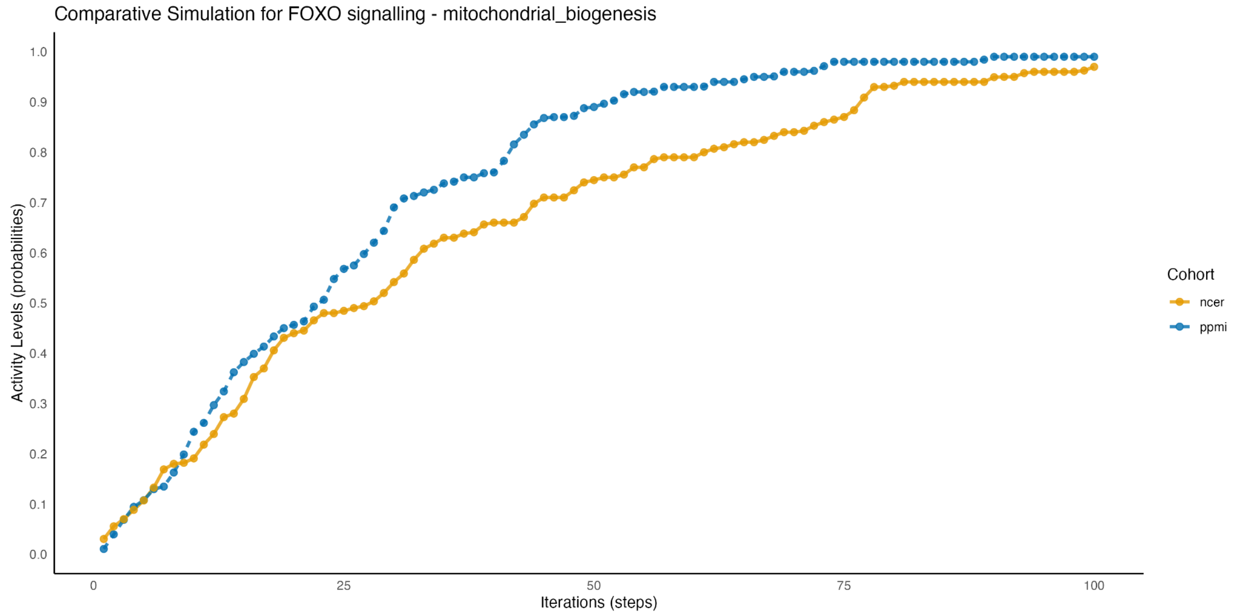


**Figure 14**: Comparative simulation of mitochondrial biogenesis within the FOXO signaling pathway between the NCER and PPMI cohorts. The analysis shows that the

PPMI cohort exhibits higher levels of mitochondrial biogenesis over time compared to NCER, suggesting differential regulation of mitochondrial function that could impact disease outcomes.

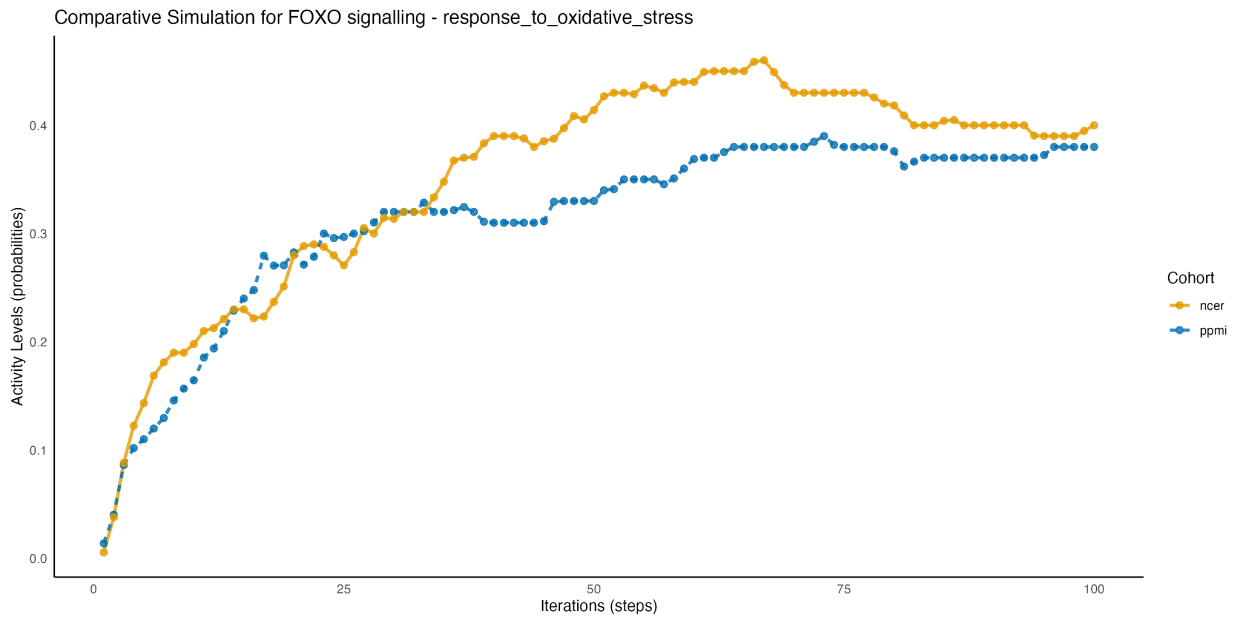


**Figure 15**: Comparative simulation of the response to oxidative stress within the FOXO signaling pathway for the NCER and PPMI cohorts. The plot shows how oxidative stress response activity evolves over 100 iterations, with the NCER cohort displaying a more pronounced and sustained increase compared to the PPMI cohort. These differences may highlight cohort-specific variations in oxidative stress management, potentially contributing to differential disease outcomes in Parkinson’s Disease

### Exosome Dynamics

Exosome dynamics play a critical role in the extracellular removal of misfolded proteins, including α-synuclein, which is central to the pathology of PD. LGALS3 (galectin-3) and PDCD6IP (ALIX) are key regulators of exosome biogenesis and release, regulating the export of aggregated proteins.

Botht al., 2014). These aggregates are implicated in the progression of PD when they disrupt cellular homeostasis and promote neuroinflammatory responses (Duijvesz et al., 2013). The gradual loss in exosome-related activity suggests that compensatory mechanisms may be involved, albeit not sufficiently robust to fully counterbalance the reduced exosome output (Kim, 2023).

Recent studies highlight the role of exosomes in the intercellular transmission of α-synuclein, further complicating the dynamics of protein clearance in PD. Exosomes originating from neurons might transport misfolded α-synuclein and facilitate its cell-to-cell transfer, thus mediating the spread of α-synuclein pathology (Yang et al., 2017; Mohamed et al., 2023). The amount of α-synuclein in exosomes has been shown to be increased in PD patients compared with controls and to correlate with the severity of the disease (Dinter et al., 2016).

The results suggest that the LGALS3 and PDCD6IP can enhance the extracellular clearance, reducing the intracellular accumulation of toxic protein aggregates and reducing the progression of PD. This is further supported by evidence that interventions targeted at enhancing exosomal release may reduce the neurotoxic effects of α-synuclein aggregates (Jiang et al., 2021; Bellingham et al., 2012). Exosome-based drug delivery systems represent an exciting approach for the design of neuroprotective therapies in PD (Cooper et al., 2014; Lashuel et al., 2012).

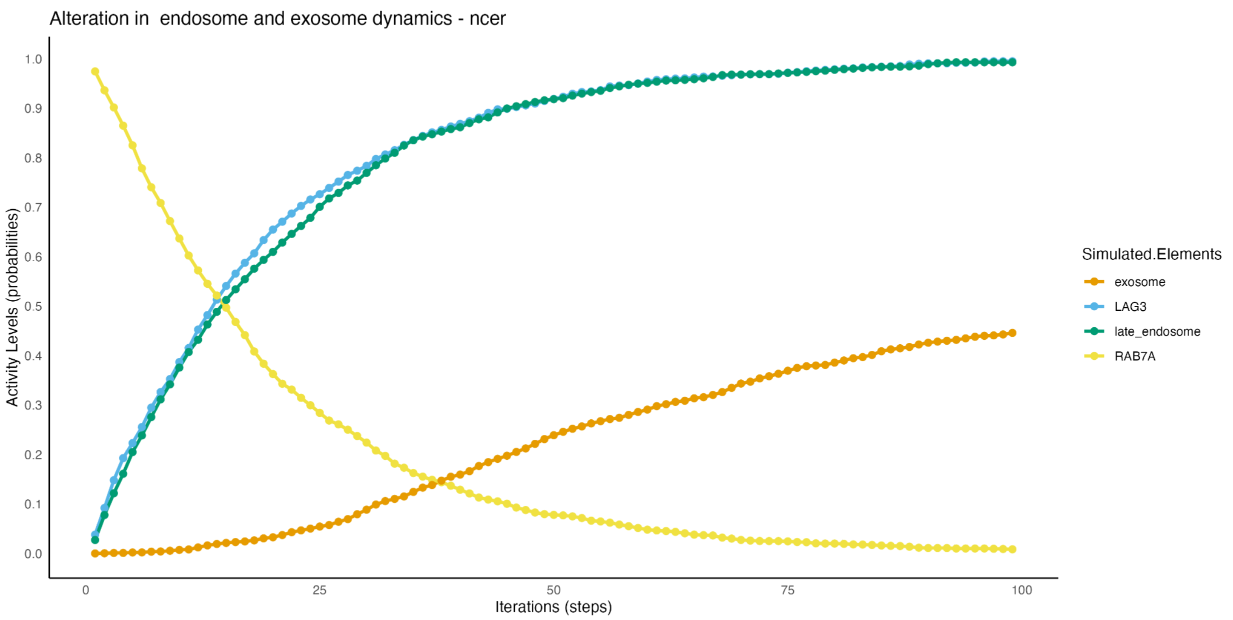


**Figure 16**: Simulation of alterations in endosome and exosome dynamics for the NCER cohort. The plot shows the activity levels of key elements (RAB7A, late endosome, LGALS3, and exosome) over 100 iterations. The results indicate that while RAB7A and late endosome activities increase over time, LGALS3 declines, and exosome activity shows a steady increase, suggesting a shift in vesicular transport dynamics in this cohort.

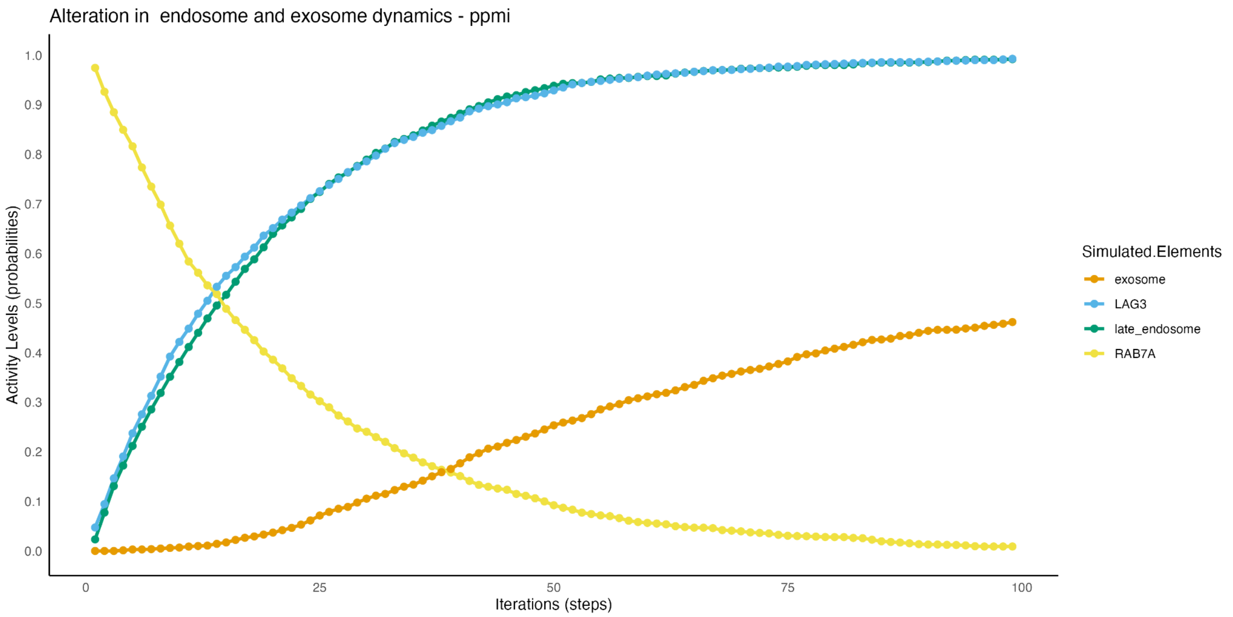

**Figure 17**: Simulation of alterations in endosome and exosome dynamics for the PPMI cohort. The graph tracks the activity levels of RAB7A, late endosome, LGALS3, and exosome components.

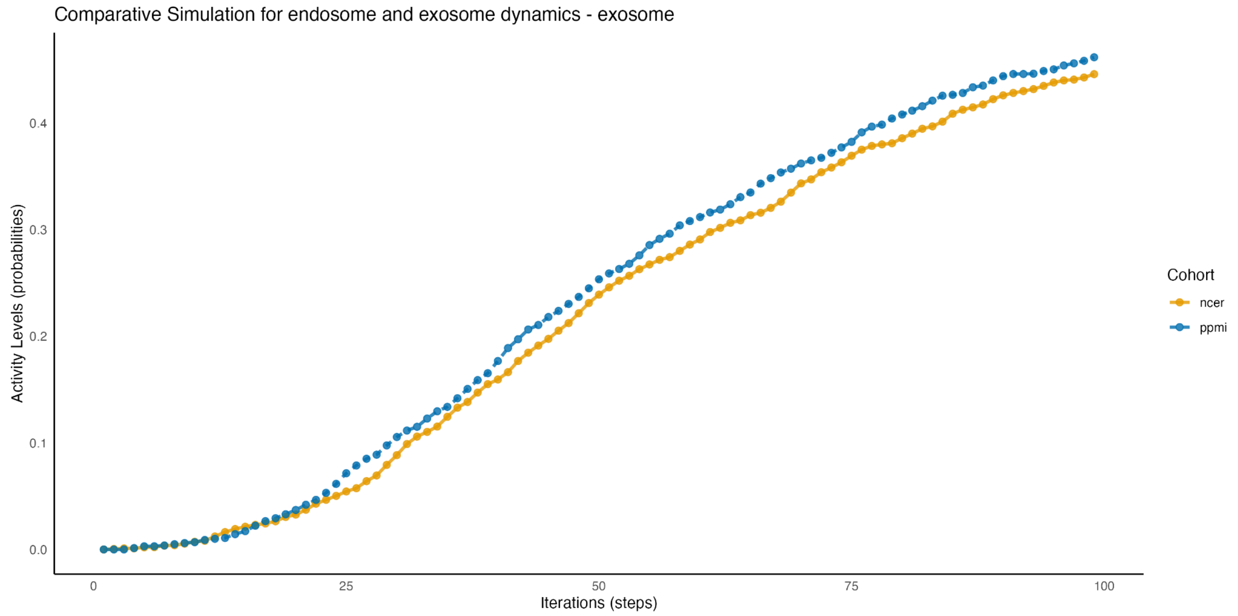


**Figure 18**: Comparative simulation focusing on exosome dynamics between the NCER and PPMI cohorts. The simulation reveals cohort-specific differences in exosome activity, with the PPMI cohort displaying a slightly higher rate of increase over 100 iterations. This difference may point to distinct exosomal processing behaviors linked to Parkinson’s Disease progression in these cohorts.

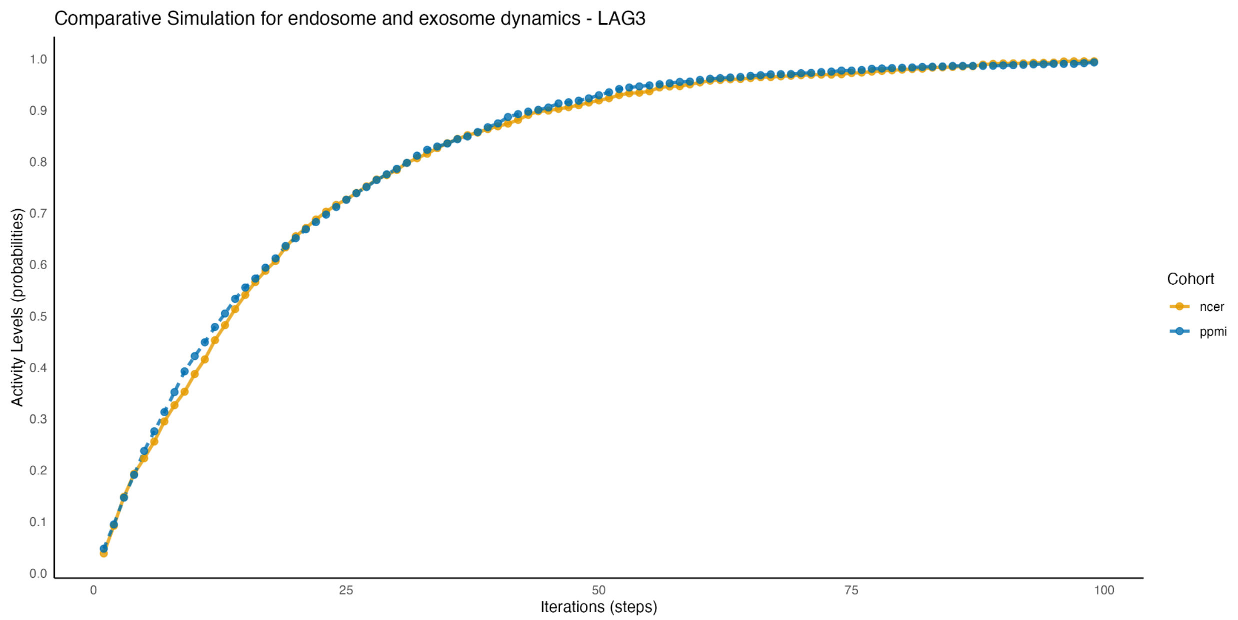


**Figure 19**: Comparative simulation of endosome and exosome dynamics focusing on LAG3 between the NCER and PPMI cohorts. The plot demonstrates that both cohorts exhibit a similar increasing trend in LAG3 activity over time, suggesting a conserved mechanism related to endosomal processing in Parkinson's Disease.

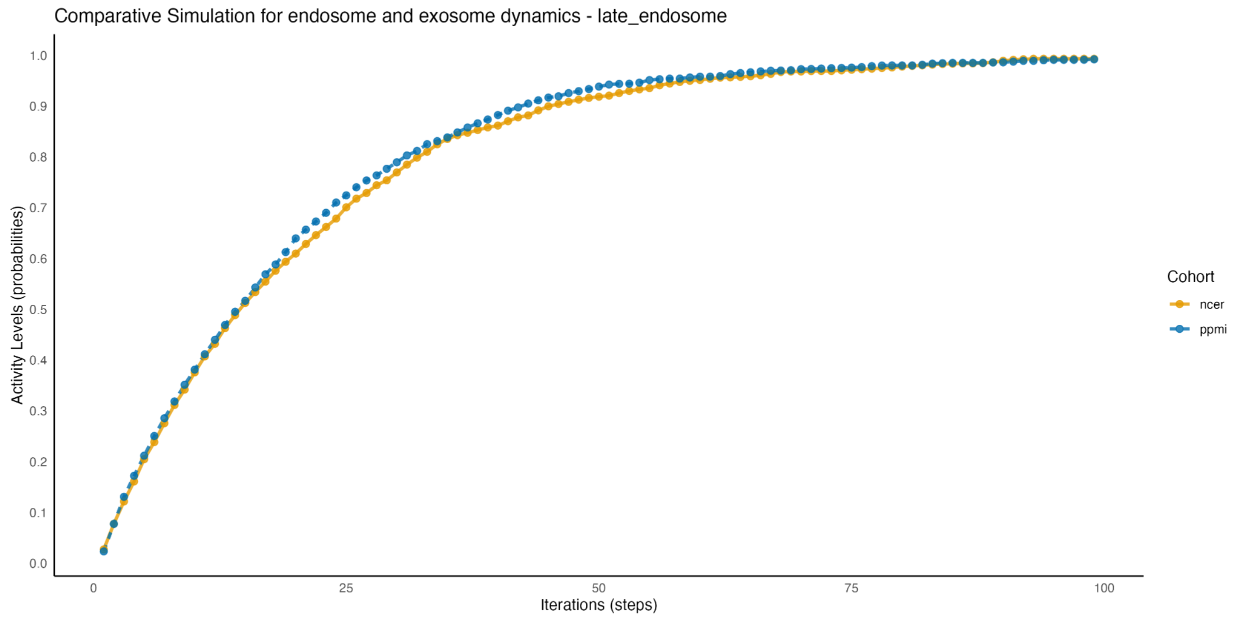


**Figure 20**: Comparative simulation of late endosome dynamics across the NCER and PPMI cohorts. The graph indicates that the activity levels of late endosome components

follow a consistent trajectory between the two cohorts, suggesting minimal cohort-specific differences in late endosome regulation.

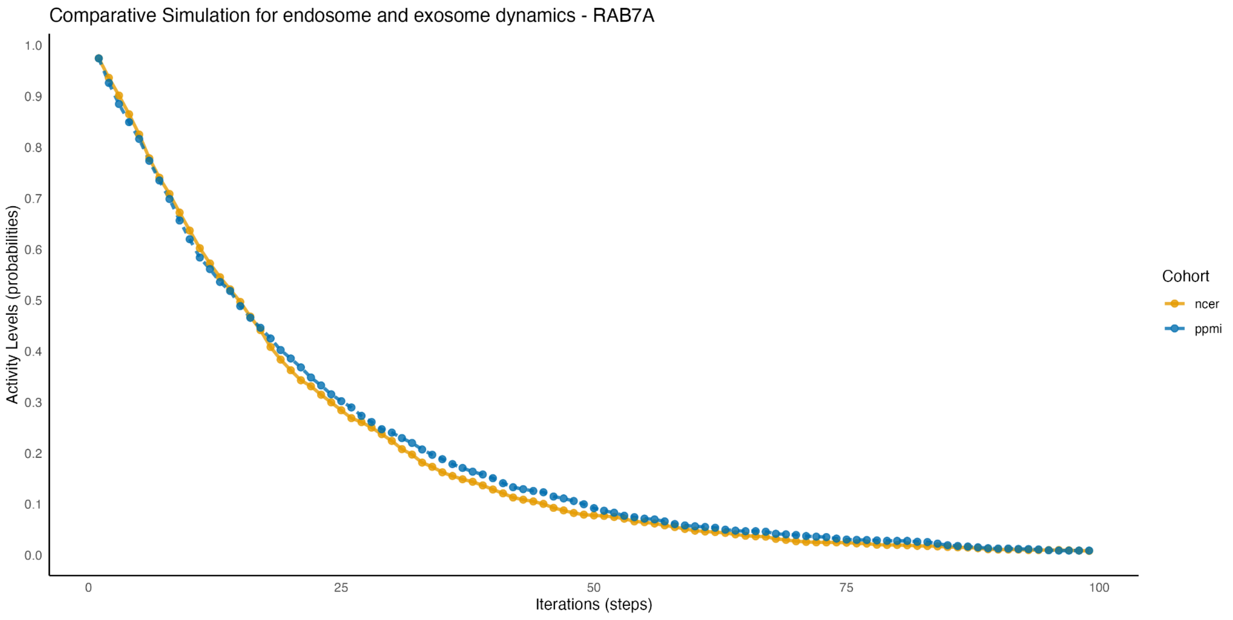


**Figure 21**: Comparative simulation of RAB7A dynamics in endosome and exosome pathways for the NCER and PPMI cohorts. The results show a rapid decline in RAB7A activity across both cohorts, indicating a potentially shared mechanism in the regulation of endosomal trafficking.

### Inflammatory Signaling & Microglial Activation

#### CD36 activity

CD36 has an important role in immune modulation, facilitating the phagocytosis of apoptotic cells and oxidised lipids. The results show an increase in CD36 activity in both cohorts with a slight higher activity in NCER than PPMI. In NCER, this increase suggests a compensatory response to decreased activity of immune-related molecules such as LGALS3 and MERTK (Dorion et al., 2023; Janda et al., 2018).

LGALS3 is involved in exosome formation and immune related responses. The reduced activity can impair exosome-mediated clearance mechanisms, hence, requiring upregulation of CD36 activity to control the neurotoxic debris (Iridoy et al., 2018; Yan, 2024). This result is aligned with studies showing that LGALS3 is dysregulated in different neurodegenerative diseases including PD, where its dysregulation can serve as a biomarker (Puigdellívol et al., 2020; Yan et al., 2023). Further, MERTK, a key molecule implicated in the clearance of apoptotic cells, shows a decreased activity in the NCER-PD cohort. The reduced activity can hinder the clearance process, reinforcing the hypothesis that CD36 upregulation acts as a compensatory mechanism in response to the impaired clearance mechanism(Dorion et al., 2023; Joers et al., 2017).

In the PPMI cohort, LGALS3 shows a decreased activity without dysregulation of MERTK. Alternative pathways might be partially compensating for the loss of the LGALS3 function, albeit with reduced efficiency(Dorion et al., 2023; Janda et al., 2018). This could explain the slight lower activity in CD36 compared to the NCER-PD cohort.

Both cohorts represent complex interplay of immune responses in PD, where the dysregulation of phagocytic mechanisms by microglia lead to neuroinflammation and neurodegeneration(Lecca et al., 2018; Butler et al., 2021). The results suggest that the increase in CD36 activity acts as a compensatory mechanism to counter the reduced activity of LGALS3 and MERTK, particularly in the NCER-PD cohort. Targeting these molecules could help to reduce the neuroinflammation and mitigate the progression of the PD by enhancing the clearance of the neurotoxic substances(Janda et al., 2018; Butler et al., 2021).

#### IFNG (Interferon Gamma) activity:

Both cohorts show an increased activity of the IFNG activity, suggesting an increased inflammatory response(Han et al., 2019; Lee et al., 2018). The increased activity of the IFNG is synchronized with elevated levels of NLRP3 and NLRP3 inflammasome. Studies show that NLRP3 inflammasomes are activated by aggregated SNCA(Han et al., 2019; Herrmann et al., 2018; Sarkar et al., 2017). The results suggest that IFNG could increase the inflammatory response, contributing to the neurodegenerative process(Lee et al., 2018; Chen et al., 2021).

The increased IFNG activity in the NCER-PD cohort is synchronized with the reduced activity of anti-inflammatory regulators such as SIRT1. SIRT1 suppress inflammatory responses and its decreased activity allows for increased inflammation(Herrmann et al., 2018).

The PPMI cohort shows a lower IFNG activity than the NCER-PD cohort. The results indicate that PRKAA1 and its phosphorylated form is activated, mitigating the inflammatory stress associated with the PD progression. This explains activating PRKAA1 molecules could modulate IFNG activity, preventing it from reaching the same peak levels as observed in the NCER-PD cohort (Zhang et al., 2020).

The results show that the IFNG activity is driven by the increased activity of NLRP3 signalling activity particularly in the NCER-PD cohort. Further, the reduced activity of SIRT1 can amplify this response. Modulating the IFNG activity could be crucial where uncontrolled inflammation may contribute to the disease progression (Cook et al., 2011).

#### MHC Class II receptor activity

MHC Class II receptors have an important role in antigen presentation and the activation of adaptive immunity. We observed a decline in MHC Class II receptor activity in the both cohorts. This decline is synchronised with a reduced activity of immune-modulator molecules such as a FCGR2B- Known for its inhibitory signalling in immune response. The downregulation of FCGR2B suggests an impairment of the regulatory feedback mechanism. This impairment can lead to the disturbance in the immune environment, exacerbating the decline of the MHC Class II activity(Kannarkat et al., 2013; Bélarbi et al., 2020).

In the NCER-PD cohort, the TF TFEB complex shows a decreased activity which is crucial for lysosomal biogenesis and function. This reduction may hinder antigen processing and presentation, thereby accelerating the decline in MHC Class II receptor activity.

Further, the PPMI cohort exhibits a reduced activity in NFE2L2 which may affect the immune balance. NFE2L2 is a key transcription factor that regulates the expression of antioxidant and cytoprotective genes in PD(Pajares et al., 2016; Gui et al., 2016; Otter et al., 2010).

The decline of the MHC Class II receptor activity and immune modulators suggests a progressive impairment in antigen presentation.Therapeutic targeting of these molecules could improve immune surveillance. The results are consistent with the fact that the immune role in PD is multifaceted, involving innate and adaptive immune response, and alteration in these responses can significantly impact the PD progression(Kannarkat et al., 2013; Heavener & Bradshaw, 2022).

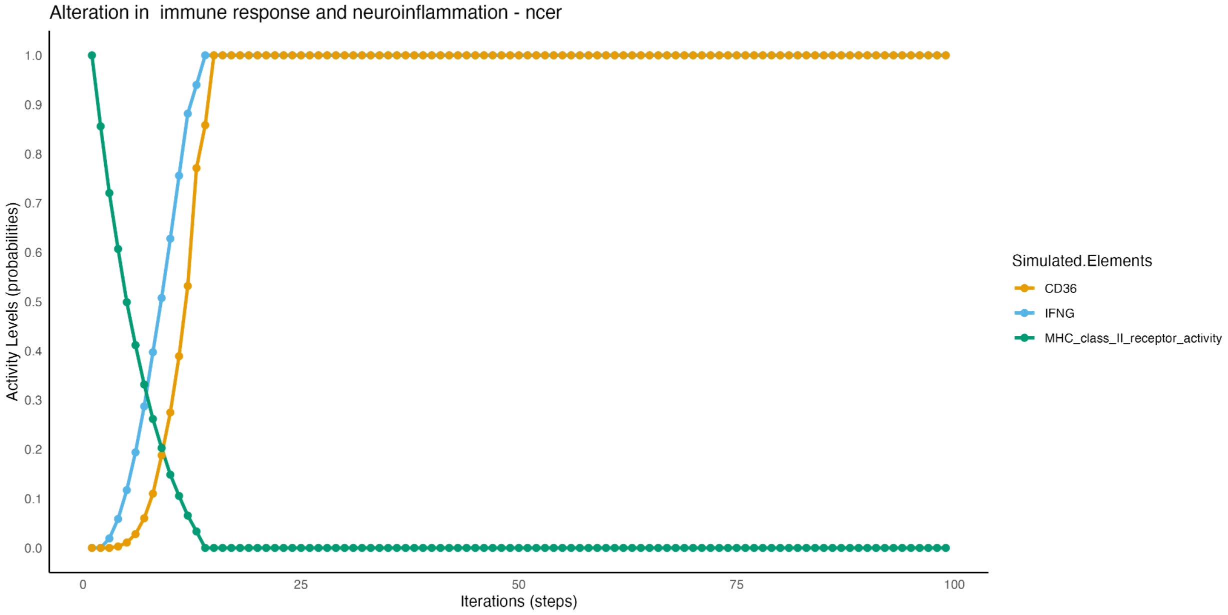


**Figure 22**: Simulation of alterations in immune response and neuroinflammation dynamics for the NCER cohort. The plot shows the activity levels of CD36, IFNG, and MHC class II receptor activity over 100 iterations. The results highlight a rapid increase in CD36 and IFNG, while MHC class II receptor activity rapidly decreases, suggesting cohort-specific immune response patterns.

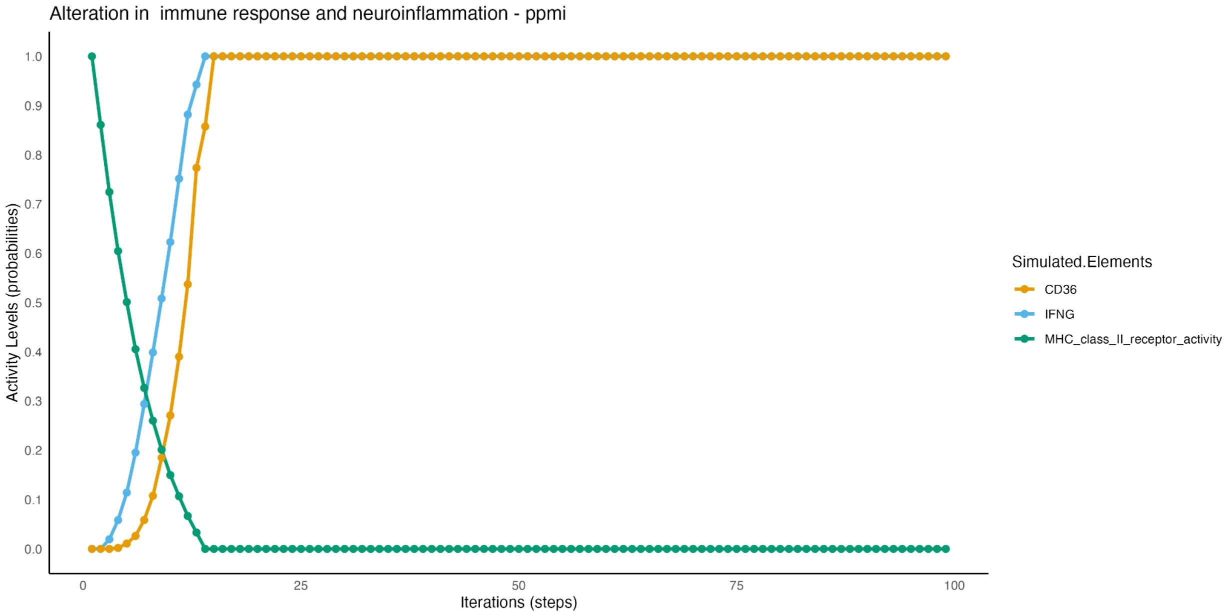


**Figure 23**: Simulation of alterations in immune response and neuroinflammation dynamics for the PPMI cohort. The graph tracks activity levels of CD36, IFNG, and MHC class II receptor activity over time. The patterns indicate similar dynamics to the NCER cohort, but with slight variations in the timing and extent of immune activation.

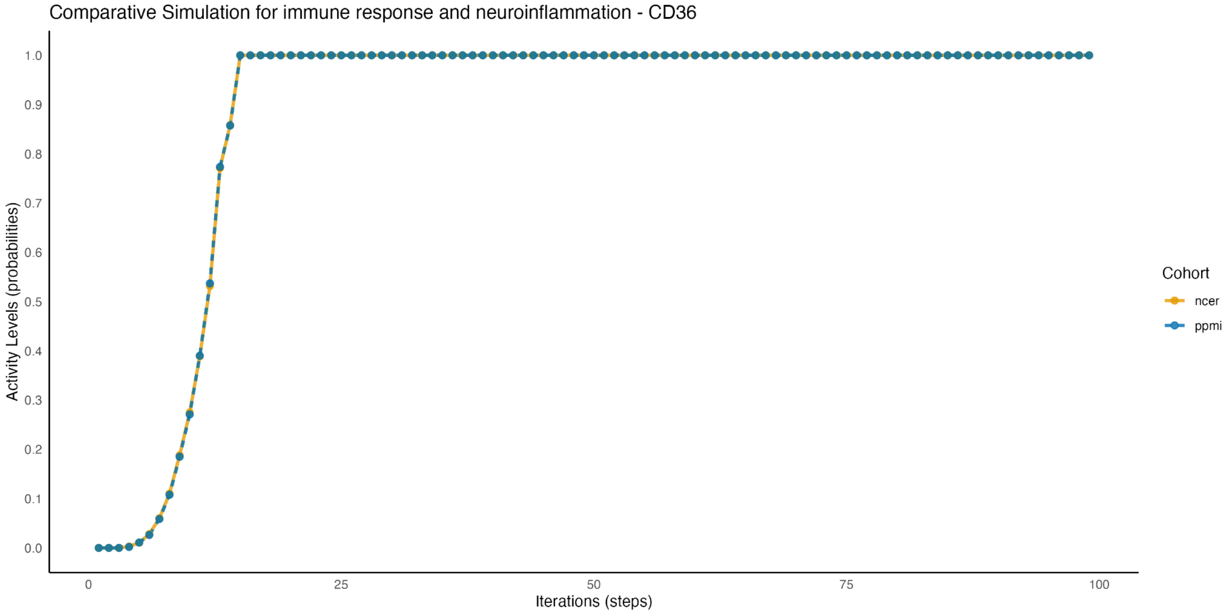

**Figure 24**: Comparative simulation focusing on CD36 activity between the NCER and PPMI cohorts. The plot shows a rapid increase in CD36 activity across both cohorts, with minimal cohort-specific differences, suggesting a conserved response in neuroinflammation.

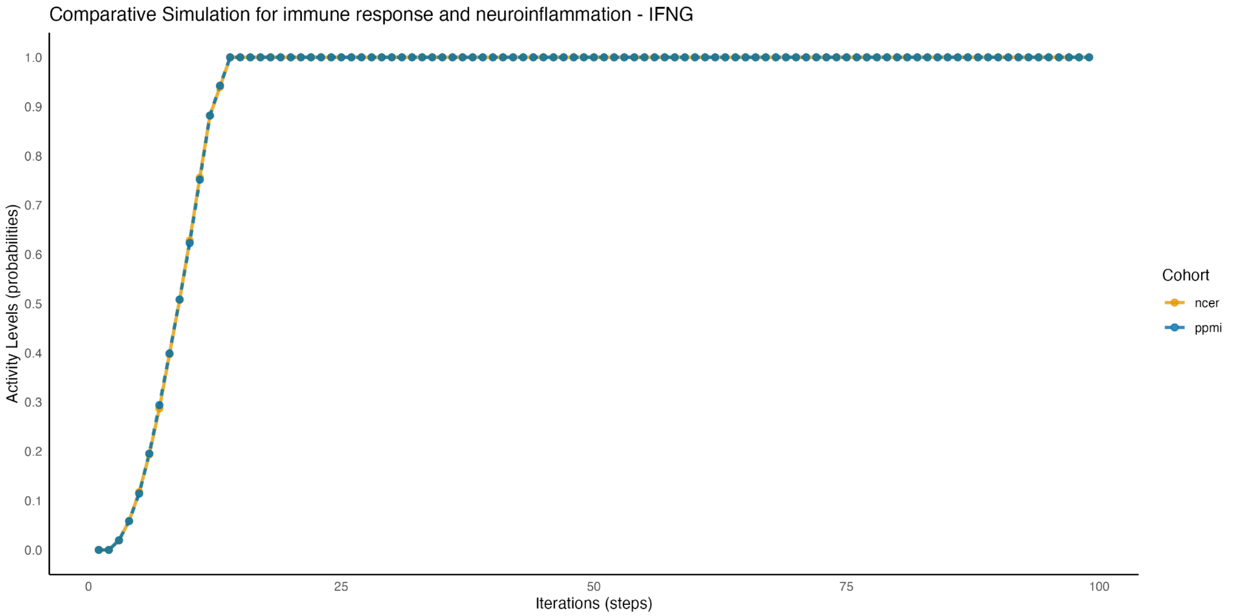


**Figure 25**: Comparative simulation of IFNG activity in immune response pathways for the NCER and PPMI cohorts. The results indicate consistent IFNG activation across both cohorts, highlighting a shared immune response pattern.

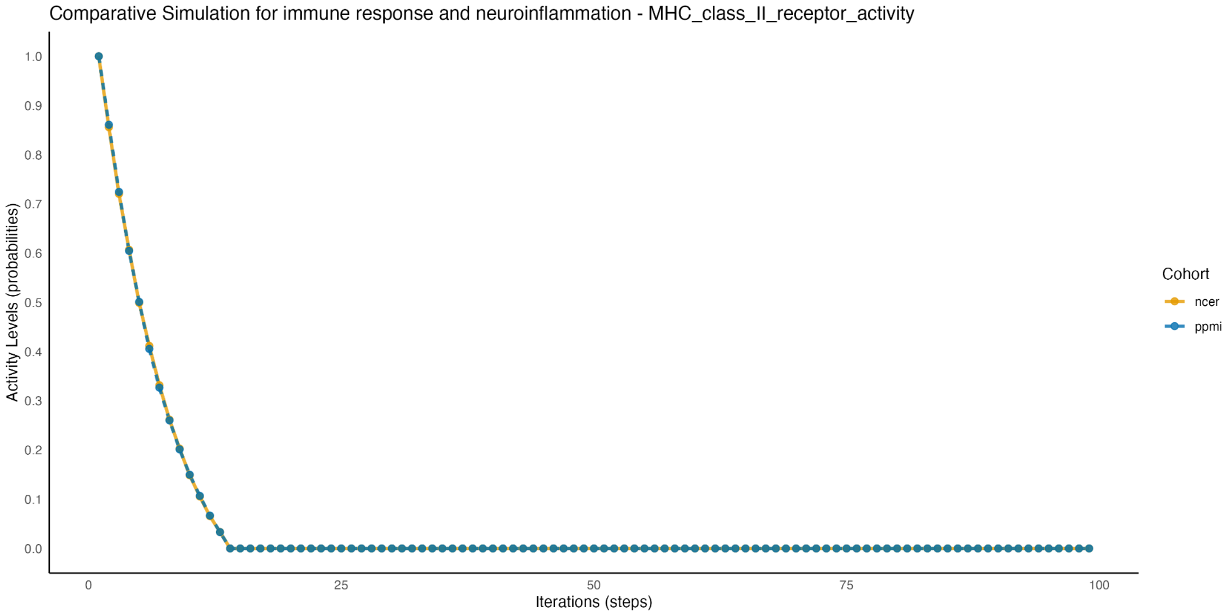


**Figure 26**: Comparative simulation of MHC class II receptor activity between the NCER and PPMI cohorts. The simulation demonstrates a rapid decline in MHC class II receptor activity in both cohorts, suggesting potential similarities in their neuroinflammatory responses.

### mTOR Pathway

#### AKT activity

Both cohorts show an elevated activity in AKT signalling. AKT signaling is critical in neuroprotection against the cytotoxic agents(Tasaki et al., 2012). The observed increase in AKT activity is synchronized with the elevated UCHL1 and UCHL1/PRKN complex in neurons, both are vital for neuronal health and proteostasis (Tasaki et al., 2012; , Greene et al., 2011).This suggests that the upregulation of these molecules may contribute to the enhanced Akt activity in both cohorts.

The NCER-PD cohort shows slightly higher AKT activity than the PPMI cohort. This behaviour is attributed to the reduced activity of DDIT4 and its phosphorylated form, which serve as inhibitors of mTOR signalng. The decreased inhibition role of DDT4 could facilitate a more activation of AKT signaling in the NCER cohort, improving cell cervical(Greene et al., 2011).

The PPMI cohort, while showing an increase in AKT signaling, shows a decreased activity of SIRT1- Known for its dual role in metabolism and inflammation. The reduced activity of SIRT1 may limit the AKT pathway activation, resulting in a lower peak compared to the NCER-PD cohort (Kim et al., 2011).

The increased AKT activity in both cohorts could improve the neuronal survival in PD since AKT signaling is crucial for the integrity of dopaminergic neurons (Tönges et al., 2012). This may indicate a compensatory response to control the inhibitory regulation. Targeting the Akt pathway may require it to be carefully calibrated to avoid the overactivation, leading to adverse effects (Jia et al., 2014; , He et al., 2021).

#### RHEB-Lysosome activity

RHEB is a key GTPase which activates the mTORC1 at lysosome- a hub for nutrient sensing and cellular metabolism. Both cohorts exhibit a consistent increase in RHEB-lysome activity. This observation aligns with the increased activity of the UCHL1-PRKN complex, which is known for its involvement in the mitochondrial control. The UCHL1-PRKN complex indirectly facilitates RHEB activation, increasing cellular energy homeostasis. Thereby, maintaining the metabolic functions essential for cell survival (Yu et al., 2018;, Lupše et al., 2021).

The NCER-cohort exhibits an increased activity of RHEB along with a reduced activity of PHLPP1. The PHLPP1 molecule is a phosphatase that negatively regulates AKT and mTOR signaling. PHLPP1 dephosphorylates AKT at critical sites (Ser473 and Thr308), modulating the cellular metabolism and growth (Du et al., 2013; , Liu et al., 2011). The lower PHLPP1 activity may increase the RHEB activity, resulting in an activation of mTORC1 at the lysosome (Yu et al., 2013; , Chen et al., 2012). Furthermore, the presence of SIRT1 in the PPMI cohort, albeit at reduced levels, may exert a modulatory effect on RHEB activity, contributing to the slightly lower peak observed in this cohort (Sharma & Dey, 2022).

Improving cellular metabolism and autophagy requires a regulation of RHEB-lysosome activity which improves mTORC1 activation in PD. The balance between the activation and the regulation is required to prevent potential overactivation, which could lead to cellular stress and compromise the cellular viability (Bradley et al., 2015; , Haque et al., 2021).

#### Glycolysis activity

Both cohorts show a notable increase in glycolytic activity. The elevated activity is consistent with the increased UCHL1 in neurons- Known for fulfilling the energy requirements of neurons (Lundgaard et al., 2015). The NCER-PD cohort shows an increased activity in glycolysis, which may be attributed to the reduced activity of DDIT4 and its phosphorylated form. DDIT4 inhibits mTOR signaling that regulates glycolysis, thereby its reduced activity could facilitate an increased glycolytic flux (Li et al., 2023).
The PPMI cohort shows a reduced activity in SIRT1 and PHLPP1 that could affect the glycolytic activity (Lu et al., 2022).
Maintaining glucose metabolism could be a viable strategy to maintain neuronal energy requirements in PD, ensuring adequate ATP production to meet cellular demands (López-Fabuel et al., 2022). The elevated glycolytic activity in both cohorts may indicate a more aggressive metabolic response, likely due to the reduced inhibitory effects of DDIT4 on mTOR signaling (Shamsi et al., 2021)(Manzo et al., 2019). The consistent and excessive glycolytic activity could lead to oxidative stress and neuronal damage.

##### 6.2.4 Catabolism activity

Both cohorts exhibit an increase in catabolic activity. This observation aligns with findings suggesting the increased catabolic processes could be a compensatory response to cellular stress (Zhou et al., 2019; Costello et al., 2023; Sanchez-Mirasierra et al., 2022).
The elevated catabolic activity coincides with a reduction in PHLPP1 and SIRT1- Molecules that inhibit mTOR signaling. The decreased activity may reduce the inhibitory role on mTOR, leading to enhanced mTOR signaling (Zhou et al., 2019; Soukas & Zhou, 2019). While mTOR activation generally improves the anabolic processes such as proteins and lipid synthesis, the dysregulation in this context paradoxically leads to

an increased catabolic activity, possibly as a compensatory response to cellular stress (Soukas & Zhou, 2019; Aman et al., 2021). This reflects the complexity of mTOR's function to maintain the metabolic homeostasis. Such aberrant signaling can disrupt the balance between anabolic and catabolic processes, contributing to disease progression(Costello et al., 2023; Basit & Vries, 2019).

The results show that the increased catabolic activity indicates a need for cellular breakdown and recycling. Metabolic shifts are often a response to cellular stressors (Costello et al., 2023; Zhang et al., 2017; Sanchez-Mirasierra et al., 2022). However, uncontrolled metabolic shifts lead to metabolic dysfunction, which commonly results in neurodegenerative diseases (Zhou et al., 2019; Li et al., 2017; Mazzulli et al., 2011). Developing therapeutic strategies can be beneficial to control the shifts, limiting the increased degeneration of cellular components (Sanchez-Mirasierra et al., 2022; Aman et al., 2021).

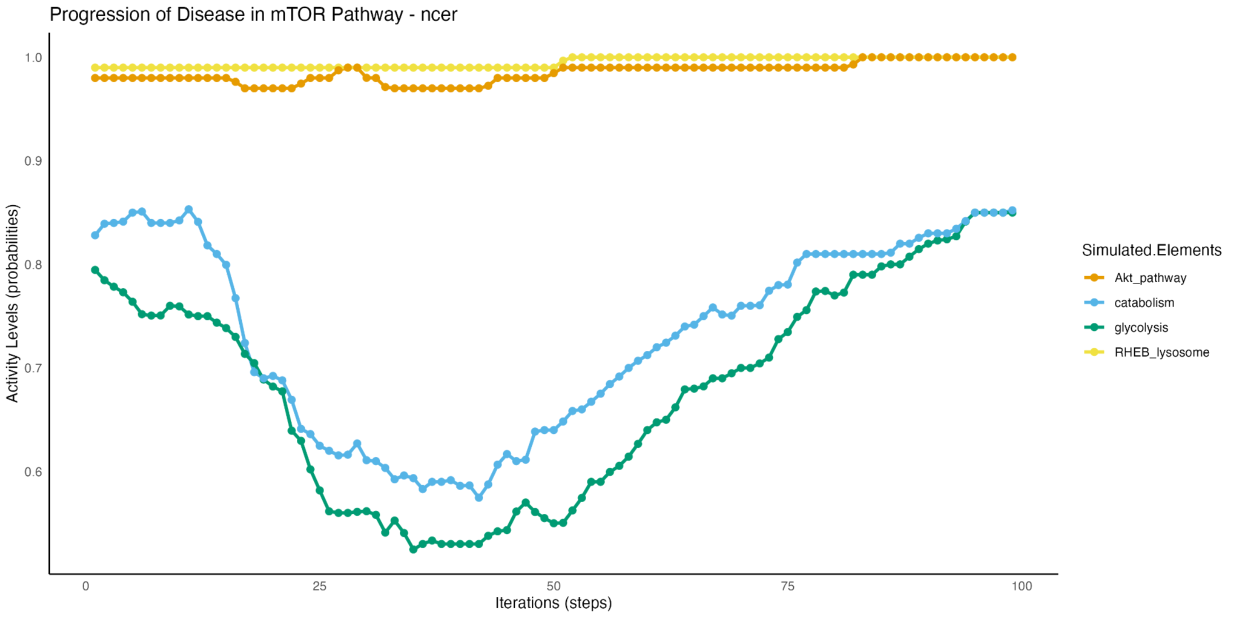


**Figure 27**: Simulation of Parkinson's Disease progression in the mTOR pathway for the NCER cohort. The graph tracks activity levels of key elements: AKT pathway, catabolism, glycolysis, and RHEB lysosome over 100 iterations. The results reveal that while the AKT pathway remains consistently active, there are fluctuations in glycolysis

and catabolism, with a notable dip followed by a recovery, indicating potential disruptions in metabolic regulation.

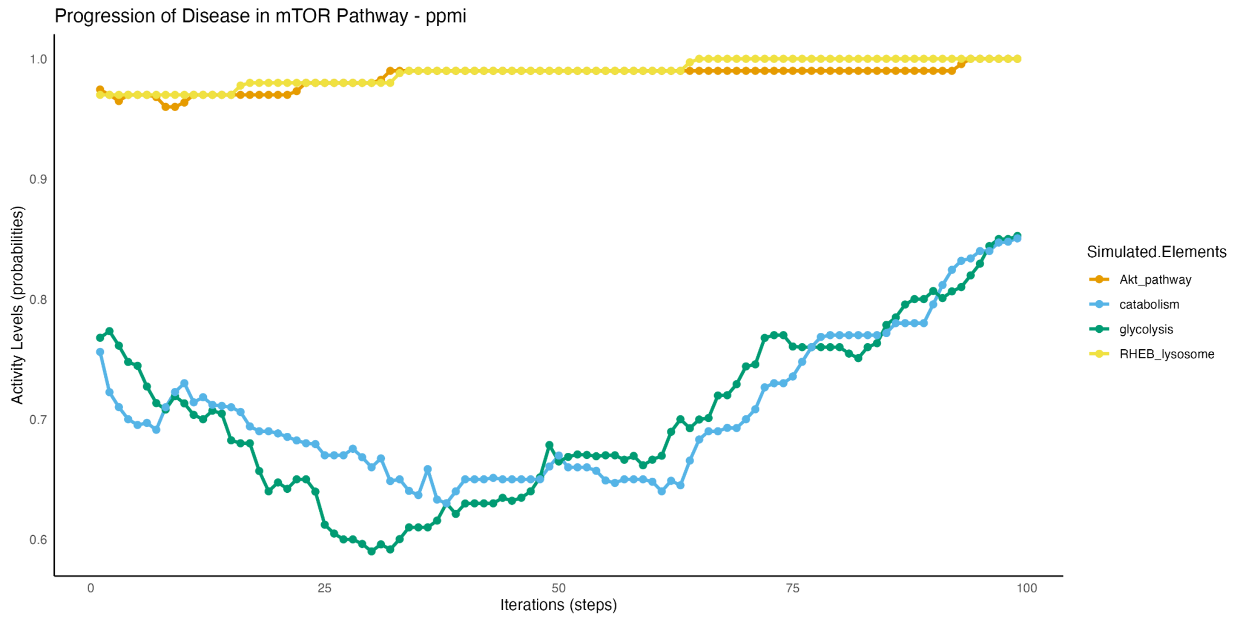


**Figure 28**: Simulation of Parkinson's Disease progression in the mTOR pathway for the PPMI cohort. The plot shows activity levels for the AKT pathway, catabolism, glycolysis, and RHEB lysosome. Similar to the NCER cohort, the AKT pathway remains steady, while fluctuations are observed in catabolism and glycolysis, suggesting metabolic dysregulation specific to Parkinson’s Disease progression in the PPMI cohort.

### Wnt/PI3K/Akt Signaling Pathway

#### TFEB_SNCA complex activity

Both cohorts show an increased activity of the TFEB SNCA complex. The elevated activity of TFEB SNCA complex, in both cohorts, coincides with the increased activity of SNCA. TFEB SNCA complex has a significant role to maintain cellular homeostasis by improving the autophagic responses and lysosomal biogenesis. The result aligns with the findings indicating that the improved autophagic responses can control the accumulation of the neurotoxic proteins and manitan cellular integrity (Banerjee et al., 2010; Menzies et al., 2017; Wong & Cuervo, 2010).

In the NCER cohort, the increased activity is synchronized with the increased activity of the GFRA1 GDNF complex beside the IGF1 and IGF1R. These molecules are known to activate autophagy and lysosomal dysfunction (Djajadikerta et al., 2020).

However, in the PPMI cohort, the increased activity is influenced by the increased activity of the PTEN and the IRS1, both limiting the TFEB activation (Dehay et al., 2015).

###### 8.2.2 TFEB phosphorylation activity

Both cohorts exhibit a decreased activity in the TFEB phosphorylation process. The phosphorylation process of the TFEB (Transcription Factor EB) is a critical mechanism that influences the activity and localisation of TFEB in the cell. The phosphorylated TFEB is sequestered in cytoplasm, while the dephosphorylation allows its translocation to the nucleus. This translocation leads to the activation of autophagic responses (Settembre et al., 2012; , Roczniak-Ferguson et al., 2012; , Napolitano & Ballabio, 2016).

### Insulin resistance activity

Both cohorts exhibit an elevated activity of insulin resistance which is recognized as a significant factor contributing to metabolic dysfunction and neurodegeneration in PD (Chohan et al., 2021; , Monte, 2013). The results show a decrease in the activity of the insulin receptor (INSR) and the INSR INS complex, indicating a predisposition to insulin resistance (Chohan et al., 2021; , Ormazábal et al., 2018).

Both cohorts show elevated mitophagy-related activity, suggesting a compensatory response within the mitochondrial dysfunction (McWilliams et al., 2018; Watzlawik, 2023). In the PPMI cohort, the increased activity coincides with increased levels of PARK7 within the mitochondria. PARK7 could protect mitochondria from oxidative damage (Argueti-Ostrovsky et al., 2021; Imberechts et al., 2022). Further, the PPMI cohort shows increased ubiquitination of TOMM20, helping in the recruitment of autophagic machinery and enhancing mitochondrial clearance (Teixeira et al., 2016). Additionally, elevated PINK1 activity in the PPMI cohort facilitates the recruitment of PRKN, leading to more efficient mitochondrial degradation (Rita et al., 2018; Zhang et al., 2014). The NCER-PD cohort exhibits increased mitophagy activity, and shows lower levels of key protective proteins such as PARK7 and PINK1. This may result in less effective mitochondrial clearance, which could contribute to ongoing mitochondrial stress (Wang et al., 2015; Zhu et al., 2016).

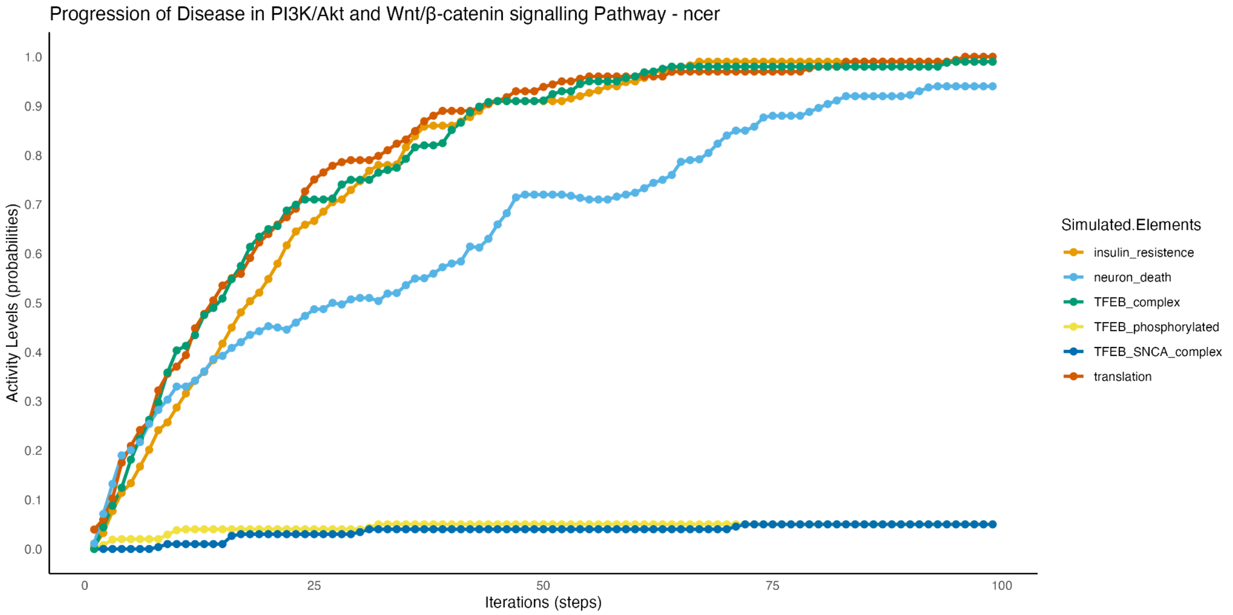


**Figure 29**: Simulation of Parkinson's Disease progression in the PI3K/Akt and Wnt/β-catenin signaling pathway for the NCER cohort. The plot shows activity levels of key elements, including insulin resistance, neuron death, TFEB complex, TFEB phosphorylated, TFEB-SNCA complex, and translation over 100 iterations. The results indicate a rapid activation of insulin resistance and translation processes, while TFEB-related activities increase gradually, highlighting potential disruptions in signaling pathways that may influence neurodegeneration.

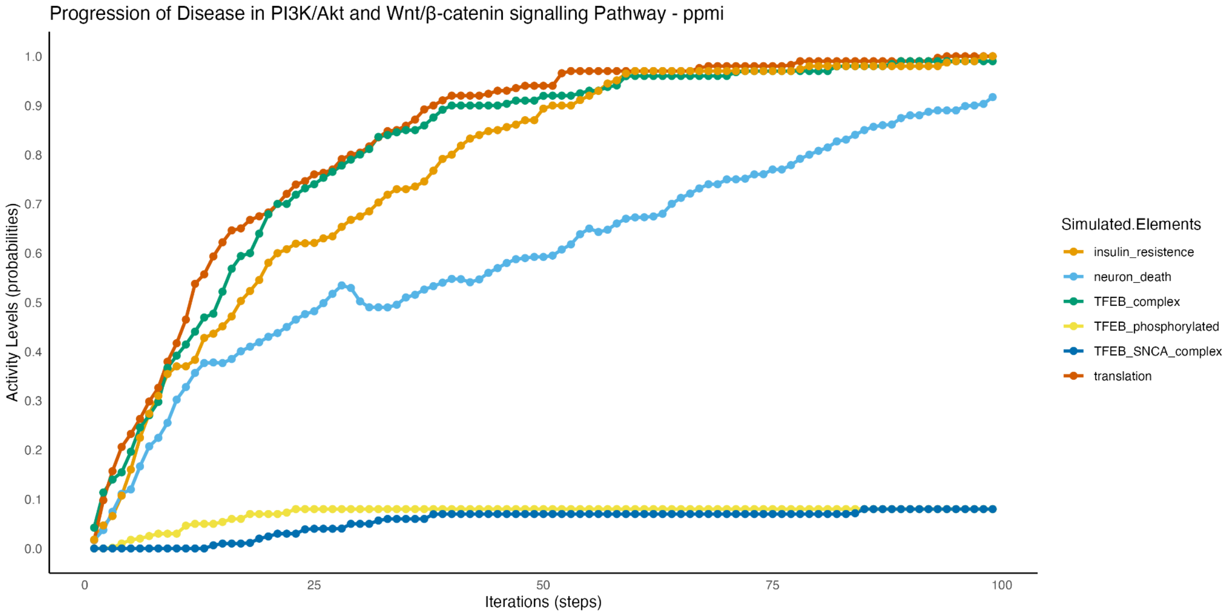


**Figure 30**: Simulation of Parkinson's Disease progression in the PI3K/Akt and Wnt/β-catenin signaling pathway for the PPMI cohort. The graph tracks changes in the activity of insulin resistance, neuron death, TFEB complex, TFEB phosphorylated, TFEB-SNCA complex, and translation. Similar trends are observed as in the NCER cohort, with some variations in the timing and extent of pathway activation, indicating cohort-specific regulatory differences in neurodegenerative mechanisms.

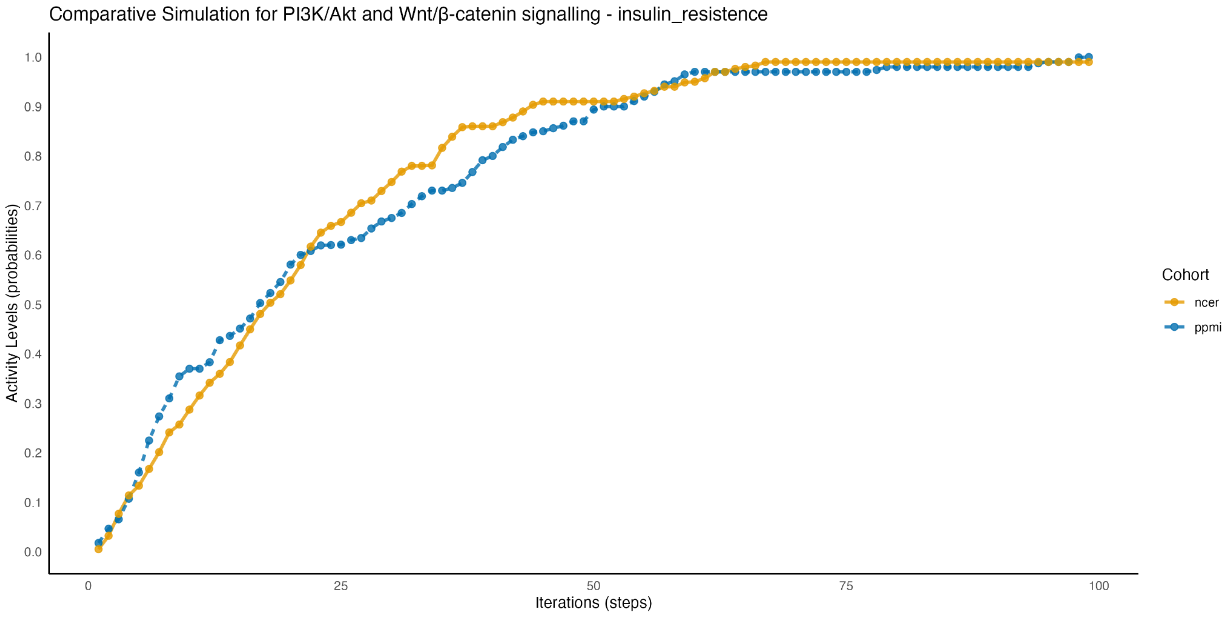


**Figure 31**: Comparative simulation of insulin resistance activity in the PI3K/Akt and Wnt/β-catenin signaling pathway between the NCER and PPMI cohorts. The plot

demonstrates that both cohorts exhibit an increase in insulin resistance over time, with the NCER cohort showing a slightly more rapid progression, suggesting cohort-specific differences in metabolic signaling related to Parkinson's Disease.

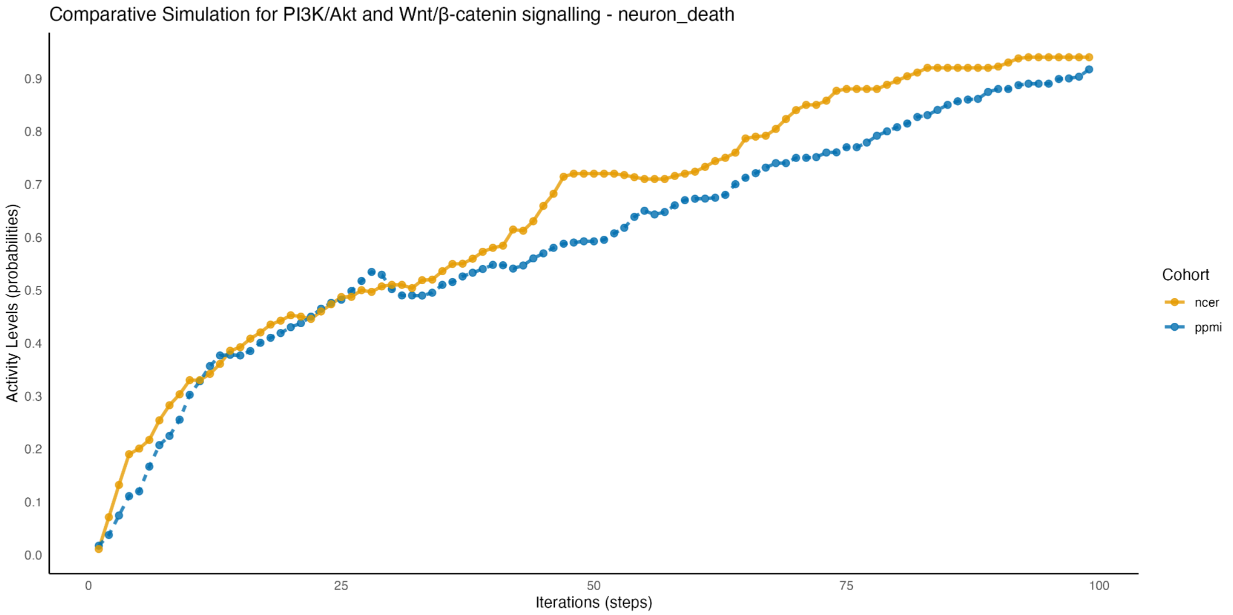


**Figure 32**: Comparative simulation of neuron death activity in the PI3K/Akt and Wnt/β-catenin signaling pathway for the NCER and PPMI cohorts. The graph highlights cohort-specific differences in neuron death dynamics, with the NCER cohort showing a higher and more sustained activation compared to PPMI. This may indicate variations in neurodegenerative processes across the cohorts.

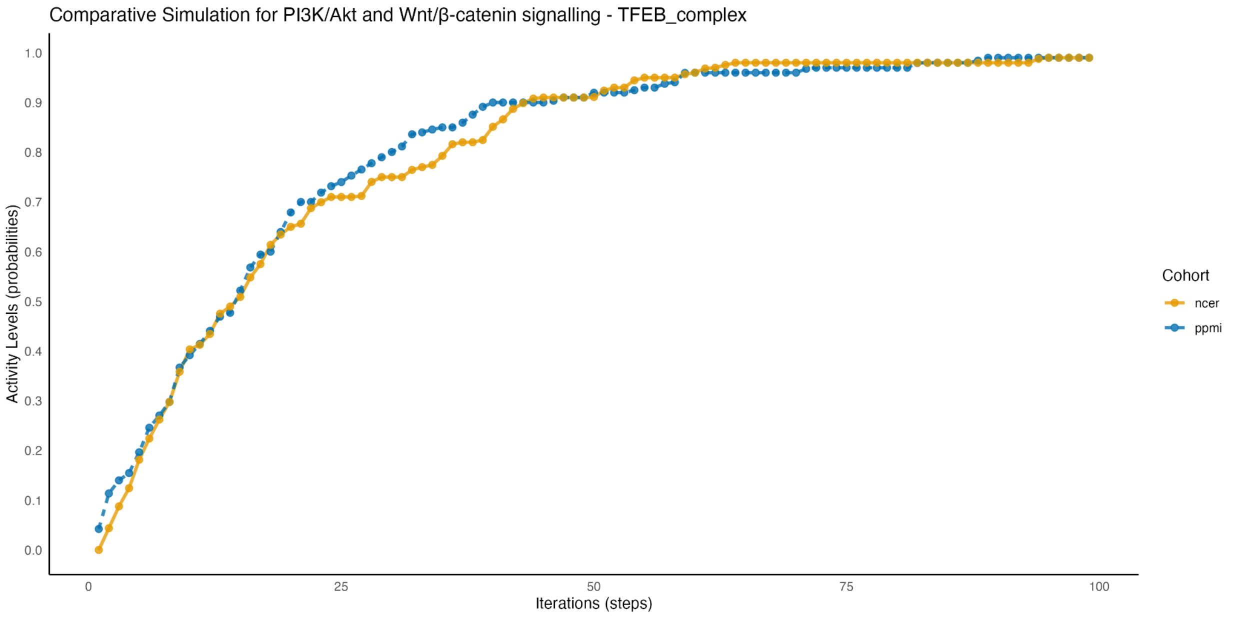

**Figure 33**: Comparative simulation of TFEB complex activity within the PI3K/Akt and Wnt/β-catenin signaling pathways between the NCER and PPMI cohorts. The plot indicates a gradual increase in TFEB complex activity for both cohorts, with the PPMI cohort showing a slightly earlier activation. This suggests cohort-specific nuances in transcription factor regulation linked to Parkinson's Disease progression.

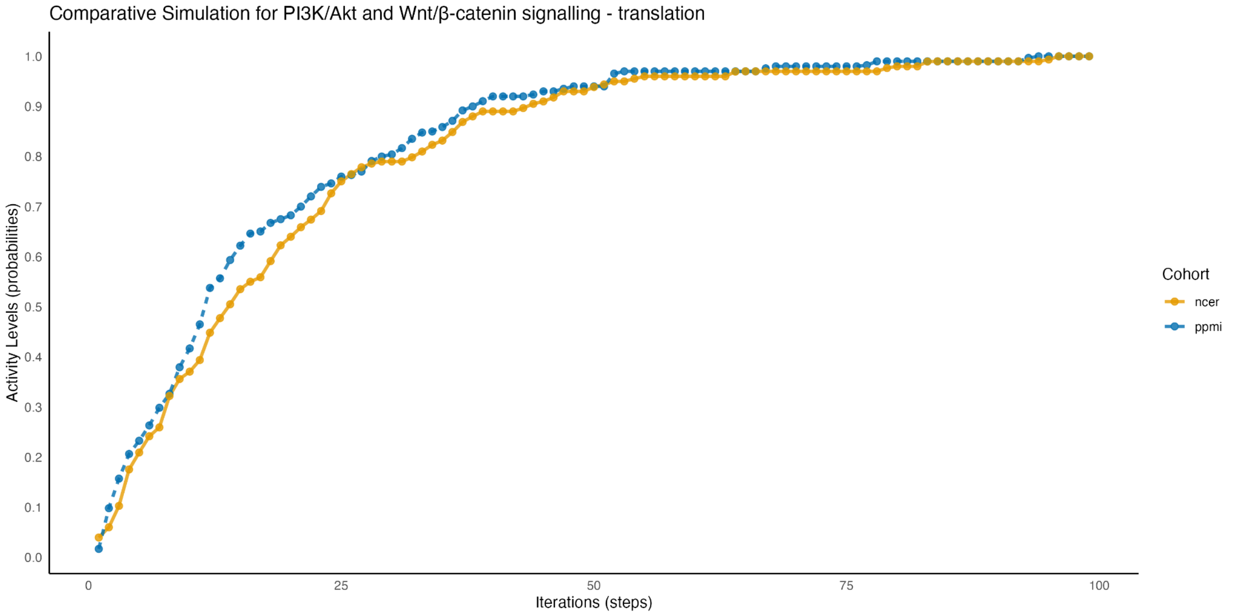


**Figure 34**: Comparative simulation of translation activity in the PI3K/Akt and Wnt/β-catenin signaling pathways for the NCER and PPMI cohorts. The results demonstrate similar trends in translation activation across both cohorts, with minor differences in the rate of progression, indicating a shared but slightly varied response in protein synthesis regulation.

Sources:

1. Sex-based Disparities in Brain Aging: A Focus on Parkinson's Disease (Beheshti et al., 2023)
2. Gender Differences in Motor and Non-Motor Symptoms in Parkinson's Disease (Abraham et al., 2023)
3. Differences in Brain Aging between Sexes in Parkinson's Disease (Beheshti et al., 2024)

This paragraph integrates your findings with relevant studies while citing important sources that support the gender-specific progression in PD. It avoids redundancy by smoothly connecting the molecular pathways with clinical observations.
